# Genome-wide Analysis of Insomnia (N=1,331,010) Identifies Novel Loci and Functional Pathways

**DOI:** 10.1101/214973

**Authors:** Philip R. Jansen, Kyoko Watanabe, Sven Stringer, Nathan Skene, Julien Bryois, Anke R. Hammerschlag, Christiaan A. de Leeuw, Jeroen Benjamins, Ana B. Muñoz-Manchado, Mats Nagel, Jeanne E. Savage, Henning Tiemeier, Tonya White, The 23andMe Research Team, Joyce Y. Tung, David A. Hinds, Vladimir Vacic, Patrick F. Sullivan, Sophie van der Sluis, Tinca J.C. Polderman, August B. Smit, Jens Hjerling-Leffler, Eus J.W. Van Someren, Danielle Posthuma

**Author notes:** These authors contributed equally to this work. Correspondence should be addressed to: Danielle Posthuma: Department of Complex Trait Genetics, VU University, De Boelelaan 1085, 1081 HV, Amsterdam, The Netherlands.

## Abstract

Insomnia is the second-most prevalent mental disorder, with no sufficient treatment available. Despite a substantial role of genetic factors, only a handful of genes have been implicated and insight into the associated neurobiological pathways remains limited. Here, we use an unprecedented large genetic association sample (N=1,331,010) to allow detection of a substantial number of genetic variants and gain insight into biological functions, cell types and tissues involved in insomnia complaints. We identify 202 genome-wide significant loci implicating 956 genes through positional, eQTL and chromatin interaction mapping. We show involvement of the axonal part of neurons, of specific cortical and subcortical tissues, and of two specific cell-types in insomnia: striatal medium spiny neurons and hypothalamic neurons. These cell-types have been implicated previously in the regulation of reward processing, sleep and arousal in animal studies, but have never been genetically linked to insomnia in humans. We found weak genetic correlations with other sleep-related traits, but strong genetic correlations with psychiatric and metabolic traits. Mendelian randomization identified causal effects of insomnia on specific psychiatric and metabolic traits. Our findings reveal key brain areas and cells implicated in the neurobiology of insomnia and its related disorders, and provide novel targets for treatment.

---

Insomnia is the second-most prevalent mental disorder^1^. One third of the general population reports insomnia complaints. The diagnostic criteria for Insomnia Disorder^2^ (i.e. difficulties with initiating or maintaining sleep with accompanying daytime complaints at least three times a week for at least three months, which cannot be attributed to inadequate circumstances for sleep^3^) are met by 10%, up to one third in samples of older age^4^. Insomnia contributes significantly to the risk and severity of cardiovascular, metabolic, mood, and neurodegenerative disorders^2^. Despite evidence of a considerable genetic component (heritability 38-59%^5^), only a small number of genetic loci moderating the risk of insomnia have thus far been identified^6,7^. Recent genome-wide association studies^6,7^ (GWAS) for insomnia complaints (N=113,006) demonstrated its polygenic architecture and implicated three genome-wide significant (GWS) loci and seven genes. A prominent role was reported for *MEIS1*, which showed pleiotropic effects for insomnia complaints and restless legs syndrome (RLS)^7^, yet the role of other genes was not unambiguously shown. We set out to substantially increase the sample size to allow the detection of more genetic risk variants for insomnia complaints, which may aid in understanding its neurobiological mechanisms. By combining data collected in the UK Biobank v2^8^ (UKB; N=386,533) and 23andMe, Inc., a personal genetics company^9,10^ (N=944,477), we obtained an unprecedented sample size of 1,331,010 individuals. Insomnia complaints were measured using questionnaire data, and the specific questions were validated to be good proxies of insomnia disorder, using an independent sample (The Netherlands Sleep Register, NSR)^11^ in which we had access to similar question data, as well as clinical interviews assessing insomnia disorder (see **Supplementary Methods 1.1-1.3**). We find 202 risk loci for insomnia, and extensive functional in silico analyses reveal the involvement of specific tissue and cell types, whereas secondary statistical analyses reveal causal effects of insomnia on metabolic and psychiatric traits.

## Meta-analysis yields 202 risk loci

UKB assessed insomnia complaints (hereafter referred to as ‘insomnia’) using a touchscreen device while 23andMe research participants completed online surveys. Assessment of insomnia in both samples shows high accuracy (sensitivity=84-98%; specificity=80-96%) for Insomnia Disorder (see **Supplementary Methods 1.3**). The prevalence of insomnia in the UKBv2 sample was 28.3%, 30.5% in the 23andMe sample, and 29.3% in the combined sample, in keeping with previous estimates for people with advanced age in the UK^4^ and elsewhere^12,13^. Older people dominate the UKB sample (mean age=56.7, SD=8.0) and the 23andMe sample (two-thirds of the sample older than 45, one-third even older than 60 years of age). Prevalence was higher in females (34.6%) than males (24.5%), yielding an odds ratio (OR) of 1.6, close to the OR of 1.4 reported in a meta-analysis^14^.

Quality control was conducted separately per sample, following standardized, stringent protocols (see **Methods**). GWAS was run separately per sample (UKB; N=386,533, 23andMe, Inc.; N=944,477) (Extended Data Fig. 1), and then meta-analyzed using METAL^15^ by weighing SNP effects by sample size (see **Methods**). We first analyzed males and females separately (**Extended Data Fig. 2**, 3), and observed a high genetic correlation between the sexes (*r_g_*=0.92, SE=0.02, **Extended Data Table 1**), indicating strong overlap of genetic effects. Owing to the large sample size, the *r_g_* of 0.92 was significantly different from 1 (one-sided Wald test, *P*=2.54×10^-6^) suggesting a small role for sex-specific genetic risk factors, consistent with our previous report^7^. However, since sex-specific effects were relatively small, we here focus on identifying genetic effects important in both sexes and continued with the combined sample (**Supplementary Table 1**, **2** and **Supplementary Discussion 2.1** provide sex-specific results).

We observe significant polygenic signal in the GWAS (lambda inflation factor=1.808) which could not be ascribed to spurious association (LD Score intercept=1.075)^16^ (Extended Data Fig. 4a). Meta-analysis identified 11,990 genome-wide significant (GWS) SNPs (*P*<5×10^-8^), represented by 248 independent lead SNPs (*r*^2^<0.1), located in 202 genomic risk loci (Fig. 1a, **Supplementary Fig. 1** and **Supplementary Table 3, 4**). All lead SNPs showed concordant signs of effect in both samples (Extended Data Fig. 4b). We confirm two (chr2:66,785,180 and chr5:135,393,752) out of six previously reported loci for insomnia ^6,7^ (**Supplementary Table 5**). Polygenic score (PGS) prediction in three randomly selected hold-out samples (N=3×3,000) estimated the current results to explain up to 2.6% of the variance in insomnia (Fig. 1b, Extended Data Fig. 5 and **Supplementary Table 6**). The SNP-based heritability (*h*^2^_*SNP*_) was estimated at 7.0% (SE=0.002). Partitioning the heritability by functional categories of SNPs (see **Methods**) showed the strongest enrichment of *h*^2^_*SNP*_ in conserved regions (enrichment=15.8, *P*=1.57×10^-14^). In addition, *h*^2^_SNP_ was enriched in common SNPs (MAF > 0.3) and depleted in more rare SNPs (MAF<0.01; Fig. 1c and **Supplementary Table 7**).

**Fig. 1a-e.**
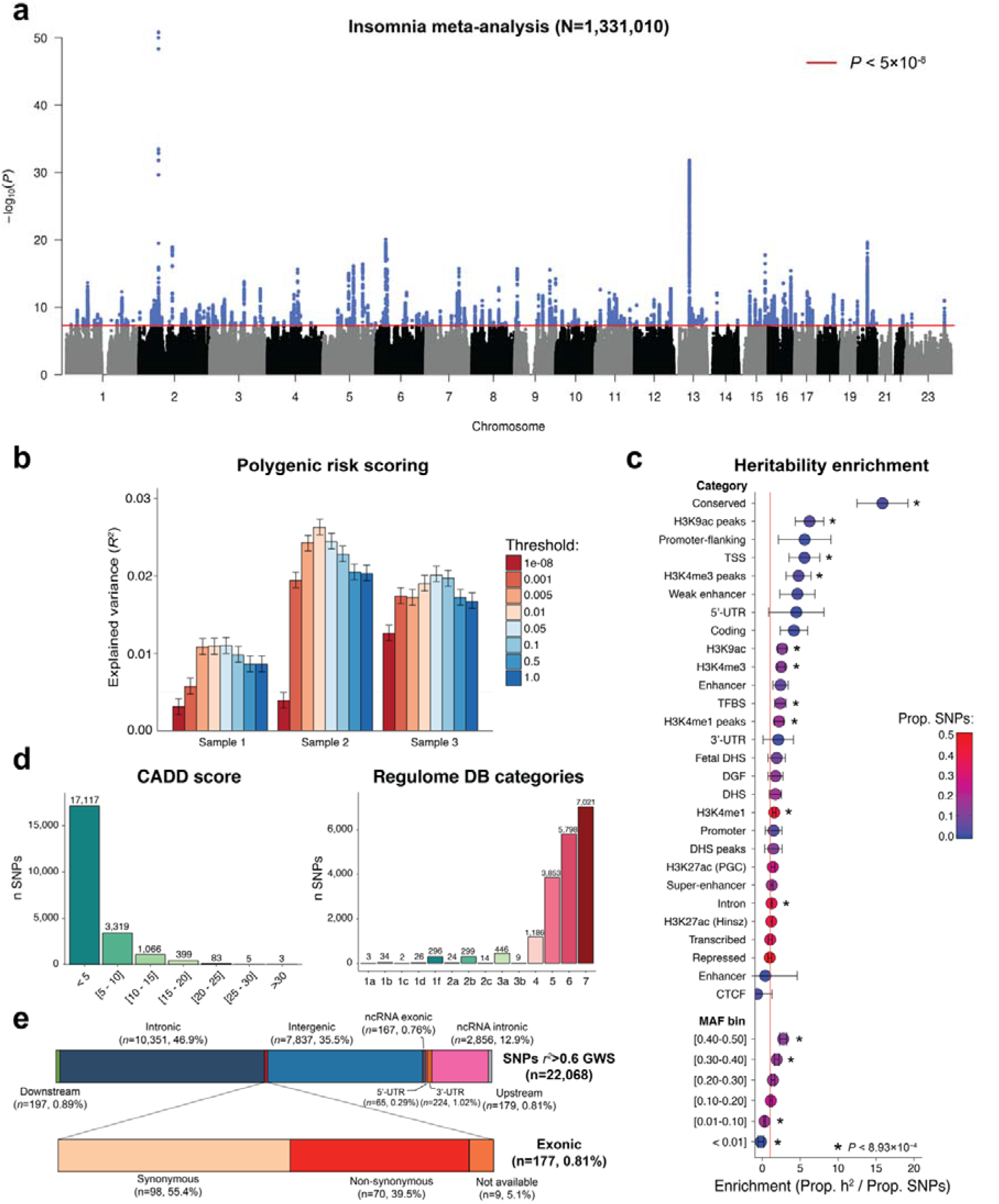
SNP-based results from the GWAS meta-analysis on insomnia (N=1,331,010). (**a**) Manhattan plot of the GWAS of insomnia, showing the -log10-transformed *P*-value for each SNP **(b)** Heritability enrichment for functional SNP categories and minor allele frequency bins (MAF). Enrichment was calculated by dividing the proportion of heritability for each category by the proportion of SNPs in that category, significant enrichments after Bonferroni correction (28 functional categories + 6 MAF bins + 22 chromosomes) are indicated by an asterisk (*P*<0.05/56=8.93×10^-4^) (**c**) Polygenic score (PGS) prediction in three hold-out samples (N=3,000), showing the increase in explained variance in insomnia (Nagelkerke’s pseudo *r*^2^) and 95% confidence interval for each *P*-value threshold. All *P*-value thresholds were statistically significant. (**d**) Distribution of CADD scores and RegulomeDB category of all annotated SNPs in LD (*r*^2^>=0.6) with one of the GWS SNPs (*n*=22,068) and (**e**) functional consequences of these SNPs.

We used FUMA^17^ to functionally annotate all 22,068 SNPs in the risk loci that were in LD (*r*^2^≥ 0.6) with one of the independent significant SNPs (see **Methods**). The majority of these SNPs (76.8%) were in open chromatin regions^18^ as indicated by a minimum chromatin state of 1-7 (Fig. 1d and **Supplementary Table 8**). In line with findings for other traits^7,19^, about half of these SNPs were in intergenic (35.5%) or non-coding RNA (13.0%) regions (Fig. 1e), and of these, 0.72% were highly likely to have a regulatory function as indicated by a RegulomeDB Score < 2 (see **Methods**). However, of these 51.5% were located inside a protein coding gene and 0.81% were exonic. Of the 177 exonic SNPs, 71 were exonic non-synonymous (ExNS, **Supplementary Table 9**). *WDR90* included four ExNS (rs7190775, rs4984906, rs3752493, and rs3803697) all in high LD with the same independent significant SNP (rs3184470). There were two ExNS SNPs with extremely high Combined Annotation Dependent Depletion (CADD) scores^20^ suggesting a strong deleterious effect on protein function: rs13107325 in *SLC39A8* (locus 56, *P*=8.31×10^-16^) with the derived allele T (MAF=0.03) associated with an increased risk of insomnia, and rs35713889 in *LAMB2 (*locus 42, *P*=1.77×10^-7^), where the derived allele T of rs35713889 (MAF=0.11) was also associated with an increased risk of insomnia complaints. **Supplementary Table 10** and **Supplementary Discussion 2.2** provide a detailed overview of the functional impact of all variants in the genomic risk loci.

## Genes implicated in insomnia

To obtain insight into (functional) consequences of individual GWS SNPs we used FUMA^17^ to apply three strategies to map associated variants to genes (see **Methods**). Positional gene-mapping aligned SNPs to 412 genes by location. Expression Quantitative Trait Loci (eQTL) gene-mapping matched cis-eQTL SNPs to 594 genes whose expression levels they influence. Chromatin interaction mapping annotated SNPs to 159 genes based on three-dimensional DNA-DNA interactions between genomic regions of the GWS SNPs and nearby or distant genes (**Supplementary Fig. 2, Supplementary Table 11** and **Supplementary Discussion 2.3**). 91 genes were mapped by all three strategies (**Supplementary Table 12**) and 336 genes were physically located outside of the risk loci but were implicated by eQTL associations (306 genes), chromatin interactions (16 genes) or both (14 genes). Several genes were implicated by GWS SNPs originating from two distinct risk loci on the same chromosome (Fig. 2a and **2b**): *MEIS1*, located on chromosome 2 in the strongest associated locus (locus 20), was positionally mapped by 51 SNPs and mapped by chromatin interactions in 10 tissue types including cross-loci interactions from locus 21, and is a known gene involved in insomnia^7^. *LRGUK*, located on chromosome 7 in locus 106, was positionally mapped by 22 SNPs and chromatin interactions in 3 tissue types including cross-loci interactions from locus 105. *LRGUK* was also implicated by eQTLs associations of 125 SNPs in 14 general tissue types. *LRGUK* was previously implicated in type 2 diabetes^21^ and autism spectrum disorder^22^ – disorders with prominent insomnia – but not yet directly implicated in sleep-related phenotypes, and is the most likely candidate to explain the observed association in loci 105 and 106.

**Fig. 2a-d.**
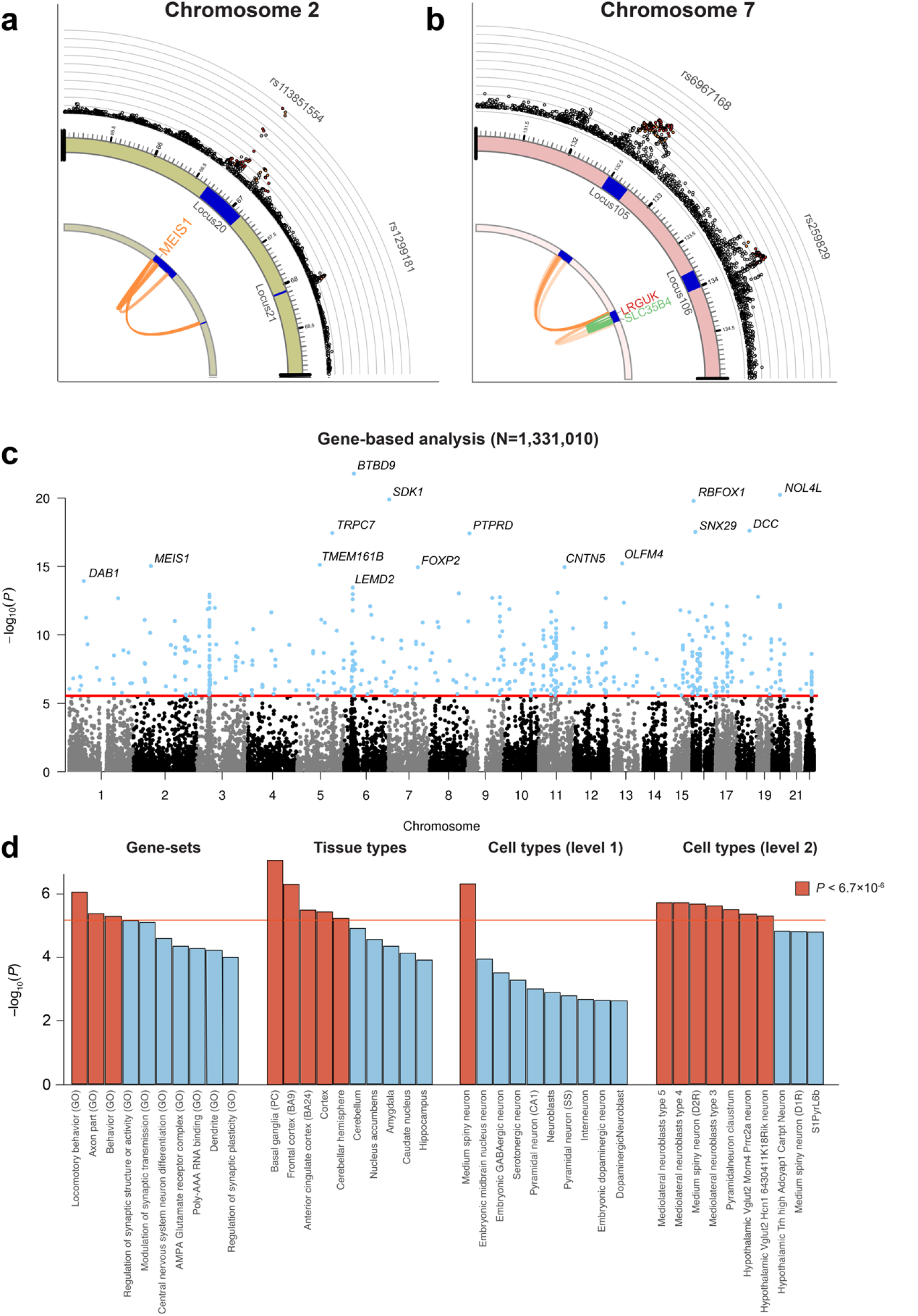
Gene-based and gene-set analyses of insomnia. Zoomed-in circos plots showing genes implicated by two genomic risk loci on chromosome 2 **(a**) and chromosome 7 **(b**), genomic risk loci indicated as blue areas, eQTL associations in green, chromatin interactions in orange. Genes mapped by by both eQTL and chromatin interactions are red. The outer layer shows a Manhattan plot containing the negative log10-transformed *P*-value of each SNP in the GWAS meta-analysis of insomnia. Full circos plots of all autosomal chromosomes are provided in **Supplementary Fig. 2**. **(c**) Genome-wide gene-based analysis (GWAS) of 18,185 genes that were tested for association with insomnia in MAGMA. The y-axis shows the negative log_10_-transformed *P*-value of the gene-based test, the x-axis shows the starting position on the chromosome. The red line indicates the Bonferroni corrected threshold for genome-wide significance (*P*=0.05/18,185=2.75×10^-6^). The top 15 most significant genes are highlighted. **(d**) Gene-set analysis of top 20 for each of the MsigDB pathways, tissue expression of GTEx tissue types, and cell types from single-cell RNA sequencing. Gene-set analyses were performed using MAGMA. The red line shows the Bonferroni significance threshold (*P*<0.05/7,473=6.7×10^-6^), correcting for the total number of tested gene-sets. Red bars indicated significant gene-sets.

Apart from linking individual associated genetic variants to genes, we conducted a genome-wide gene-based association analysis (GWGAS) using MAGMA^23^. GWGAS provides aggregate association *P-*values based on all variants located in a gene, and complements the three FUMA mapping strategies (see **Methods**). GWGAS identified 517 associated genes (Fig. 2c and **Supplementary Table 13**). The top gene *BTBD9* (*P*=8.51×10^-23^) on chromosome 6 in locus 81 was also mapped by positional and eQTL mapping (tissue type: left ventricle of the heart), and is part of a pathway regulating circadian rhythms. *BTBD9* has been associated with RLS, periodic limb movement disorder^24,25^ and Tourette Syndrome^26^. Involvement in sleep regulation was shown in *Drosophila^27^*, and mouse mutants show fragmented sleep^28^ and increased levels of dynamin 1^29^, a protein that mediates the increased sleep onset latency following pre-sleep arousal^30^.

Of the 517 MAGMA-based associated genes, 222 were outside of the GWAS risk loci, and 309 were also mapped by FUMA. In total, 956 unique genes were mapped by at least one of the three FUMA gene mapping strategies or by MAGMA (Extended Data Fig. 6). Of these, *MEIS1*, *MED27, IPO7* and *ACBD4* confirmed previous results^6,7^ (**Supplementary Table 14**). Sixty-two genes were implicated by all four mapping strategies indicating that apart from a GWS gene-based *P*-value, there were (i) GWS SNPs located inside these genes, (ii) GWS SNPs associated with differential expression of these genes and (iii) GWS SNPs that were involved in genomic regions interacting with these genes. We note that genes that were indicated by positional mapping and GWS gene-based *P*-values, but not via eQTL or chromatin interaction mapping (N=54 genes), may be of equal importance, yet are more likely to exert their influence on insomnia via structural changes in the gene products (i.e. at the protein level) and not via quantitative changes in the availability of the gene products.

## Implicated pathways, tissues and cell-types

To test whether GWS genes converged in functional gene-sets and pathways, we conducted gene-set analyses using MAGMA (see **Methods**). We tested associations of 7,473 gene-sets: 7,246 sets derived from the MsigDB^31^, gene expression values from 54 tissues from the GTEx database^32^, and cell-specific gene expression in 173 types of brain cells (Fig. 2d, **Supplementary Table 15**). Competitive testing was used and a Bonferroni corrected threshold of *P*<6.7×10^-6^ (0.05/7,473) to correct for multiple testing. Of the MsigDB gene-sets, three Gene Ontology (GO) gene-sets survived multiple testing: GO:*locomotory behavior* (*P*=8.95×10^-7^), GO:*behavior* (*P*=5.23×10^-6^), and GO:*axon part* (*P*=4.25×10^-6^). Twelve genes (*LRRK2, CRH, DLG4, DNM1, DRD1, DRD2, DRD4, GRIN1, NTSR1, SNCA, CNTN2*, and *CALB1*) were included in all of these gene-sets and two of these (*SNCA, DNM1*) had a GWS gene-based *P*-value (**Supplementary Table 16**). *SNCA* encodes alpha-synuclein and has been implicated in REM sleep behavior disorder^33^ and Parkinson’s disease^34^. Altered expression in mice changes sleep and wake EEG spectra^35^ along the same dimensions that have been implicated in insomnia disorder^36^. *DNM1* encodes the synaptic neuronal protein dynamin 1, which is increased in *BTBD9* mutant mice^29^ and mediates the sleep-disruptive effect of pre-sleep arousal (see above; *BTBD9* is the top associated gene). Conditional gene-set analyses suggested that the association with the gene-set *behavior* is almost completely explained by the association of *locomotory behavior*, and that the effect of *axon part* is independent of this (**Supplementary Discussion 2.4**). GO:*locomotory behavior* includes 175 genes involved in stimulus-evoked movement^37^. This set included 16 GWS genes: *BTBD9*, *MEIS1*, *DAB1, SNCA, GNAO1 ATP2B2, NEGR1*, *SLC4A10, GIP, DNM1, GPRC5B, GRM5, NRG1, PARK2, TAL1*, and *OXR1*). GO:*axon part* reflects a very general cellular component representing 219 genes, of which 14 were GWS (*KIF3B, SNCA, GRIA1, CDH8, ROBO2, DNM1, RANGAP1, GABBR1, P2RX3, NRG1, POLG, DAG1, NFASC*, and *CALB2*). Tissue specific gene-set analyses showed strong enrichment of genetic signal in genes expressed in the brain. Correcting for overall expression, four specific brain tissues reached the threshold for significance: overall cerebral cortex (*P*=3.68×10^-6^), Brodmann area 9 (BA9) of frontal cortex (*P*=5.04×10^-7^), BA24 of the anterior cingulate cortex (*P*=3.25×10^-6^), and cerebellar hemisphere (*P*=5.93×10^-6^)^1^. Several other brain tissues also showed strong enrichment just below threshold, including three striatal basal ganglia (BG) structures (nucleus accumbens, caudate nucleus, putamen). To test whether genes expressed in all three BG structures together would show significant enrichment of low *P*-values, we used the first principal component (BG_PC_) of these BG structures and found significant enrichment (*P*=8.33×10^-8^). When conditioning the three top cortical structures on the BG_PC_, they were no longer significantly associated after multiple testing correction (minimum *P*=0.03), which was expected given that the BG_PC_ correlated strongly with gene-expression in cortical (and other) areas (*r*>0.96). Similar results were obtained vice versa, i.e. using the first principal component of all cortical areas and conditioning the three BG structures on this resulted in no evidence of enrichment of low *P*-values for BG structures (minimum *P*=0.53). These results show that (i) genes expressed in brain are important in insomnia, (ii) genes expressed in cortical areas are more strongly associated than genes expressed in BG, (iii) there is a strong correlation between gene expression patterns across brain tissues, which suggests involvement of general cellular signatures more than specific brain tissue structures.

Brain cell type-specific gene-set analyses was first carried out on 24 broad cell-type categories. Cell type-specific gene expression was quantified using single cell RNA-sequencing of disassociated cells from somatosensory cortex, hippocampus, hypothalamus, striatum and midbrain from mouse (see **Methods**), which closely resembles gene-expression in humans^38^. Results indicated that genes expressed specifically in the medium spiny neurons (MSN, *P*=4.83×10^-7^) were associated with insomnia, and no other broad cell-types specific gene-set survived our strict threshold of *P*<6.7×10^-6^. MSNs represent 95% of neurons within the human striatum, which is one of the four major nuclei of the subcortical BG. Specifically, the striatum consists of the ventral (nucleus accumbens and olfactory tubercle) and dorsal (caudate nucleus and putamen) subdivisions. The association with MSNs thus likely explains the observed association of the BG striatal structures (nucleus accumbens, caudate nucleus, putamen).

Using broad cell classes risks not detecting associations that are specific to distinctive yet rare cell types; to account for this we then tested 149 specific brain cell-type categories, and found significant associations with 7 specific cell types: medio-lateral neuroblasts type 3, 4 and 5 (*P*=2.36×10^-6^, *P*=1.88×10^-6^, and *P*=1.87×10^-6^), D2 type medium spiny neurons (*P*=2.12×10^-6^), claustrum pyramidal neurons (*P*=3.09×10^-6^), hypothalamic Vglut2 Morn4 Prrc2a neurons (*P*=4.36×10^-6^), and hypothalamic Vglut2 Hcn16430411 K18 Rik neurons (*P*=4.98×10^-6^), known to have the densest number of melatonin receptors. These results suggest a role of distinct mature and developing cell types in the midbrain and hypothalamus. The hypothalamus contains multiple nuclei that are key to the control of sleep and arousal, including the suprachiasmatic nucleus (SCN) that accommodates the biological clock of the

## Low genetic overlap with sleep traits

Other sleep-related traits may easily be confounded with specific symptoms of insomnia, like early morning awakening, difficulties maintaining sleep, and daytime sleepiness. The most recent genome-wide studies for other sleep-related traits included 59,128 to 128,266 individuals, and assessed genetic effects on morningness^6,40,41^ (i.e. being a morning person), sleep duration ^6,41^, and daytime sleepiness/dozing ^41^. Using increased sample sizes for each of these sleep-related traits (max N=434,835), we here investigated to what extent insomnia and other sleep-related traits are genetically distinct or overlapping. We performed GWAS analyses for the following six sleep-related traits: morningness, sleep duration, ease of getting up in the morning, naps during the day, daytime dozing, and snoring (**Supplementary Methods 1.1-1.2**, Extended Data Fig. 7, 9). Of the 202 risk loci for insomnia, 39 were also GWS in at least one of the other sleep-related traits (Fig. 3, **Supplementary Table 17**). The strongest overlap in loci was found with sleep duration, with 14 out of 49 sleep duration loci overlapping with insomnia. Insomnia showed the highest genetic correlation with sleep duration (-0.47, SE=0.02; **Supplementary Table 18**) compared to other sleep-related traits, which was not surprising given that insomnia also shared the most risk loci with sleep duration (See further discussion sleep phenotypes in **Supplementary Discussion 2.5**).

**Fig. 3a-f.**
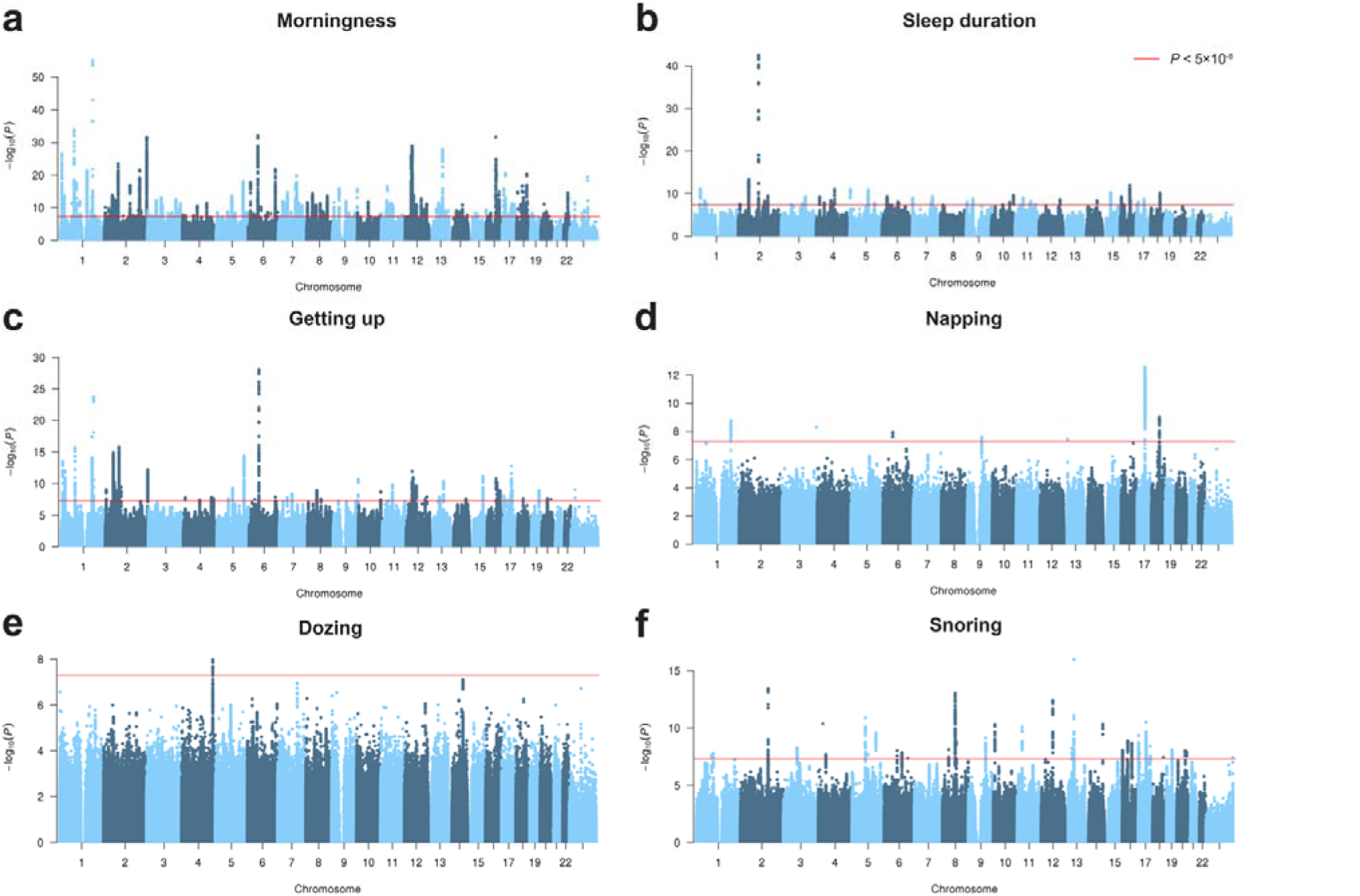
Genome-wide analyses of six sleep-related traits. Manhattan plots of the genome-wide association analyses of (**a**) Morningness (N=434,835). (**b**) Sleep duration (N=384,317) (**c**) Ease of getting up (N=385,949) (**d**) Napping (N=386,577) (**e**) Daytime dozing (N=385,333) and (**f**) Snoring (N=359,916). The y-axis shows the negative log_10_-transformed SNP *P*-value, the x-axis the base pair position of the SNPs on each chromosome. The red line indicates the Bonferroni corrected significance threshold (*P*<5×10 ^8^).

Gene-mapping of SNP associations of sleep-related traits resulted in 973 unique genes (Extended Data Fig. 9, **Supplementary Table 19-23**). Gene-based analysis showed that of the 517 GWS genes for insomnia, 120 were GWS in at least one of the other sleep-related traits, and one gene (*RBFOX1*) was GWS in all except napping and dozing (**Supplementary Table 24**). The largest proportion of overlap in GWS genes for insomnia was again with sleep duration, with 37 of the 135 (27%) GWS genes for sleep duration also GWS for insomnia. There was overlap in tissue enrichment in cortical structures and basal ganglia between insomnia and both morningness and sleep duration. On the single cell level, the medium spiny neurons were also implicated for morningness and sleep duration, but not for the other sleep-related traits (**Supplementary Table 25**). Taken together, these results suggest that at a genetic level, insomnia shows partial overlap with sleep duration, but minimal overlap with other sleep-related traits. Consistent short sleep across nights occurs only in a minor part of insomnia patients, even in a clinical sample^42^.

## Strong overlap between insomnia and psychiatric and metabolic traits

We confirm previously reported genetic correlations between insomnia and neuropsychiatric and metabolic traits^6,7^ (**Supplementary Table 26**), and also identify several GWS SNPs for insomnia that have previously been associated with these traits (**Supplementary Table 27**). The strongest correlations were with depressive symptoms (*r_g_*=0.64, SE=0.04 *P*=1.21×10^-71^), followed by anxiety disorder (*r_g_*=0.56, SE=0.11 *P*=1.40×10^-7^), subjective well-being (*r_g_*=-0.51, SE=0.03 *P*=4.93×10^-52^), major depression (*r_g_*=0.50, SE=0.07 *P*=8.08×10^-12^) and neuroticism (*r_g_*=0.48, SE=0.02 *P*=8.72×10^-80^). Genetic correlations with metabolic traits ranged between 0.09-0.20. The genetic correlations between insomnia and psychiatric traits were also stronger than the correlations between insomnia and the other sleep-related traits. Since a similar high reliability has been reported for both sleep and psychiatric phenotypes, the findings suggest that genetically insomnia more closely resembles neuropsychiatric traits than it resembles other sleep-related traits (Fig. 4). To infer directional associations between insomnia and these correlated traits, we performed bidirectional Multi-SNP Mendelian Randomization (MR) analysis^43^ (see **Methods**). Results support a direct risk effect of insomnia on metabolic syndrome phenotypes including BMI (*b_xy_*=0.36, SE=0.05, *P*=1.25×10^-12^) type 2 diabetes (*b_xy_*=0.62, SE=0.11, *P*=2.29×10^-8^), and coronary artery disease (*b_xy_*=0.61, SE=0.09, *P*=2.88×10^-12^). In addition, insomnia was bidirectionally associated with educational attainment, with a stronger effect from insomnia on educational attainment (*b_xy_*=-0.32, SE=0.02, *P*=1.68×10^-77^) (i.e. a higher risk for insomnia leads to lower educational attainment) than vice versa (*b_xy_*=-0.10, SE=0.01, *P*=2.27×10^-23^), the same pattern was observed for intelligence. We also found risk effects of insomnia on several psychiatric traits, including schizophrenia (*b_xy_*=0.68, SE=0.10, *P*=5.12×10^-11^), ADHD (*b_xy_*=0.77, SE=0.06, *P*=2.50×10^-45^), neuroticism (*b_xy_*=0.46, SE=0.03, *P*=3.92×10^-53^) and anxiety disorder (*b_xy_*=0.47, SE=0.10, *P*=4.11×10^-6^), with no evidence of large reverse effects, except for a small risk effect of neuroticism on insomnia (*b_xy_*=0.09, SE=0.02, *P*=1.24×10^-6^) and depressive symptoms (*b_xy_*=0.09, SE=0.02, *P*=1.24×10^-6^)^2^. Overall, there was only a small proportion of SNPs showing pleiotropy between insomnia and other traits (**Supplementary Table 28** and **Supplementary Discussion 2.6**).

**Fig. 4.**
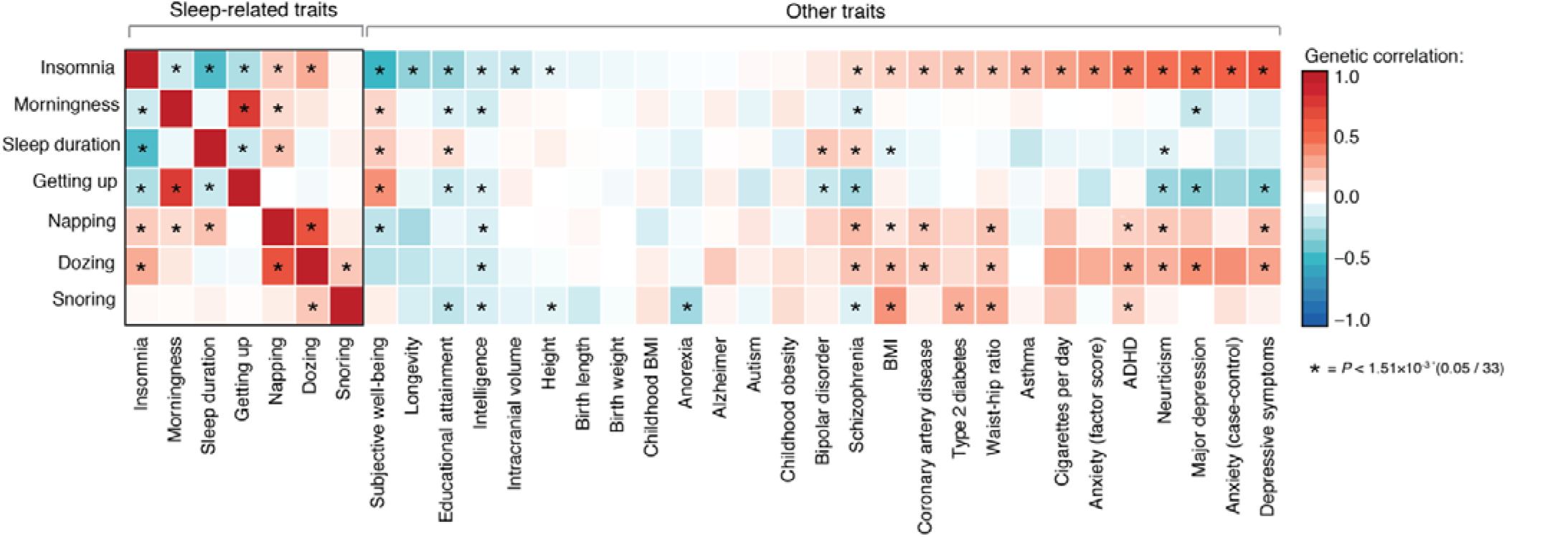
Genetic overlap of insomnia with other sleep-related traits and psychiatric and metabolic traits. Heatmap of genetic correlations between insomnia, sleep-related phenotypes and neuropsychiatric and metabolic traits studies that were calculated using LD Score regression. Red color indicates a positive *r_g_* while green indicates negative *r_g_*. Correlations that were significant after Bonferroni correction (*P<*0.05/33=1.5110^-3^) are indicated with an asterisk (see also **Supplementary Table 18, 26**).

## Discussion

In the largest GWAS study to date of 1,331,010 participants we identified 202 genomic risk loci for insomnia. Using extensive functional annotation of associated genetic variants, we demonstrated that the genetic component of insomnia points towards a role of genes involved in locomotory behavior, and genes expressed in specific cell types from the claustrum, hypothalamus and striatum, and specifically in MSNs (Fig. 5). MSNs are GABAergic inhibitory cells and represent 95% of neurons in the human striatum, one of the four major nuclei of the BG (for reviews, see ^44-46^). MSNs receive massive excitatory glutamatergic input from the cerebral cortex and the thalamus, and are targets of dopamine neurons in substantia nigra and the ventral tegmental area. In addition, they receive inhibitory inputs from striatal GABAergic interneurons. MSNs themselves are GABAergic output neurons with exceptionally long projections to globus pallidus (GP), substantia nigra and ventral pallidum, and control the activity of thalamocortical neurons. Previous studies during the natural sleep-wake cycle, *in vitro*, and from anesthetized *in vivo* preparations have shown that MSNs show fast, synchronized cyclic firing, i.e. the so-called Up and Down states, during slow-wave sleep and irregular pattern of action potentials during wakefulness. In fact, MSNs were the first neurons in which the Up and Down states characteristic of slow wave sleep were described^47^. Cell body-specific striatal lesions of the rostral striatum induce a profound sleep fragmentation, which is most characteristic of insomnia. A role for BG in sleep regulation is also suggested by the high prevalence of insomnia in neurodegenerative disorders, such as Parkinson’s Disease and Huntington’s disease in which the BG are affected. Vetrivelan et al.^44^ proposes a cortex-striatum-GP_external_-cortex network involved in the control of sleep–wake behavior and cortical activation, in which midbrain dopamine disinhibits the GP_external_ and promotes sleep through activation of D2 receptors in this network. Furthermore, brain imaging studies have suggested the caudate nucleus of the striatum as a key node in the neuronal network imbalance of insomnia^48^, and also reported abnormal function in the cortical areas we found to be most enriched (BA9^49^, BA24^50^). Our results support the involvement of the striato-cortical network in insomnia, by showing enrichment of risk genes for insomnia in cortical areas as well as the striatum, and specifically in MSNs. We recently showed that, along with several other cell types, MSNs also mediate the risk for mood disorders^51^ and schizophrenia^38^. MSNs are strongly implicated in reward processing and future work could address whether the genetic overlap between insomnia and mood disorders is mediated by gene function in MSNs.

**Fig. 5.**
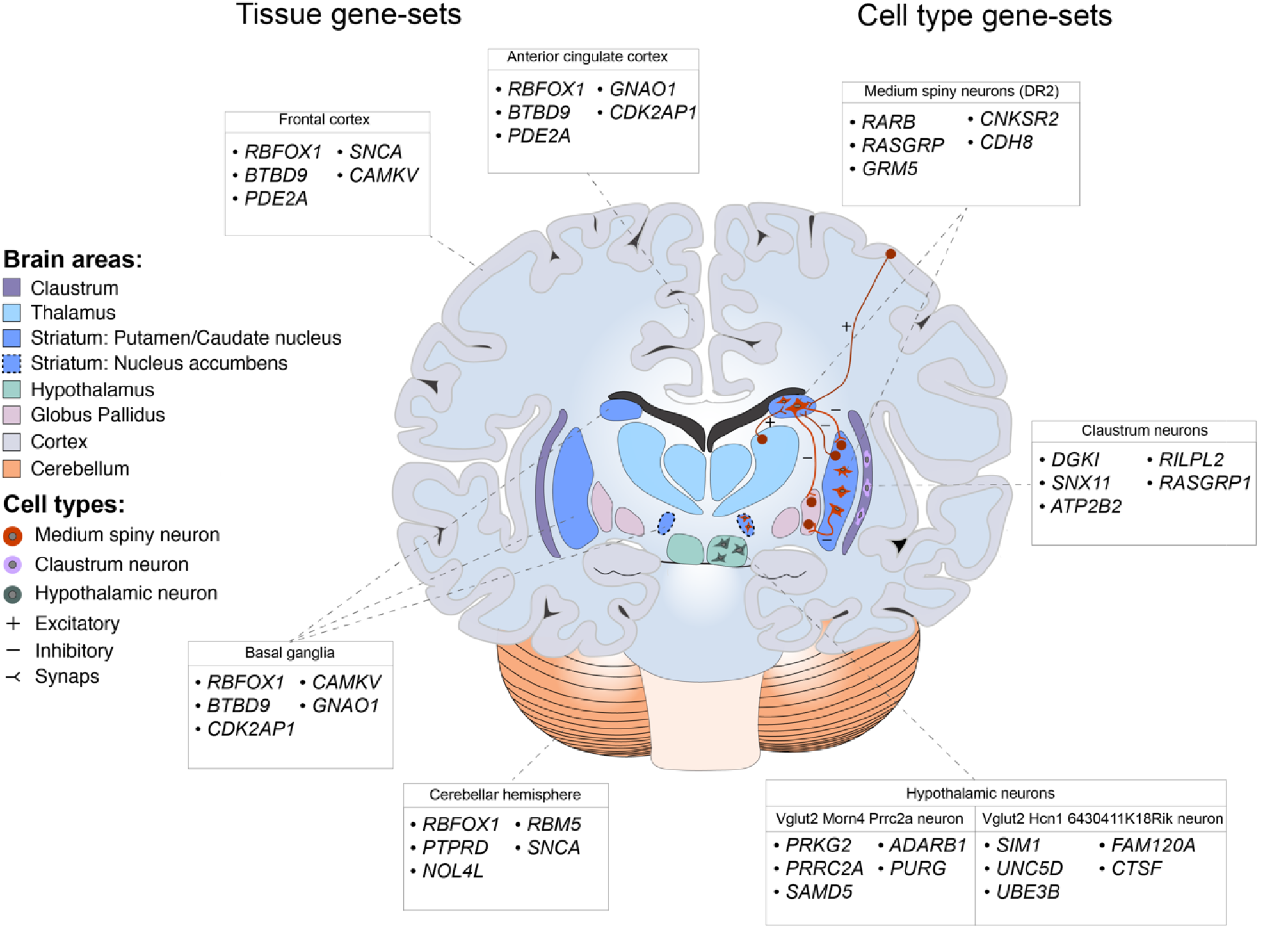
Overview of brain tissues and cell types associated with insomnia based on GWAS results from 1,331,010 individuals. For each associated gene-set, the top 5 genes driving the association are reported for each brain area and cell-type. Results for GTEx brain tissue type gene-sets are shown on the left side of the figure, while results from the level 2 single-cell gene expression are shown on the right.

Our results also showed enrichment of insomnia genes in pyramidal neurons of the claustrum. This subcortical brain region is structurally closely associated with the amygdala and has been implicated in salience coding of incoming stimuli and binding of multisensory information into conscious percepts^52^. These functions are highly relevant to insomnia, because the disorder is characterized by increased processing of incoming stimuli^53^ and by ongoing consciousness even during sleep, a phenomenon known as sleep state misperception^54^. We also found enrichment of insomnia genes in mediolateral neuroblasts from the embryonic midbrain and in two hypothalamic cell types. The role of the mediolateral neuroblasts is less clear; although they were obtained from the embryonic midbrain, it is at present unknown what type of mature neurons they differentiate into. We note that the midbrain is similar on a bulk transcriptomic level to the pons^55^, and lacking cells from that region we cannot conclusively say that midbrain cell-types are enriched. The current findings provide novel insight into the causal mechanism of insomnia, implicating specific cell types, brain areas and biological functions. These findings are starting points for the development of new therapeutic targets for insomnia and may also provide valuable insights for other, genetically related disorders.

## Methods

### Meta-analysis

A meta-analysis on the GWAS results of insomnia and morningness in UKB and 23andMe cohorts was performed using fixed-effects meta-analysis METAL^15^, using SNP *P-*values weighted by sample size. To investigate sex-specific genetic effects, we ran the meta-analysis between UKB and 23andMe datasets for males and females separately.

### Genomic risk loci definition

We used FUMA^17^ (http://fuma.ctglab.nl/), an online platform for functional mapping and annotation of genetic variants, to define genomic risk loci and obtain functional information of relevant SNPs in these loci. FUMA provides comprehensive annotation information by combining several external data sources. We first identified *independent significant SNPs* that have a genome-wide significant *P*-value (<5×10^-8^) and are independent from each other at *r*^2^<0.6. These SNPs were further represented by *lead SNPs*, which are a subset of the independent significant SNPs that are in approximate linkage equilibrium with each other at *r*^2^<0.1. We then defined associated *genomic risk loci* by merging any physically overlapping lead SNPs (linkage disequilibrium [LD] blocks <250kb apart). Borders of the genomic risk loci were defined by identifying all SNPs in LD (*r*^2^?0.6) with one of the independent significant SNPs in the locus, and the region containing all these *candidate SNPs* was considered to be a single independent genomic risk locus. LD information was calculated using the UK Biobank genotype data as a reference. Risk loci were defined based on evidence from independent significant SNPs that were available in both 23andMe and UKB. We note that SNPs that were GWS but only available in the 23andMe dataset were not included when defining genomic risk loci and were not included in any follow-up annotations or analyses, because there was no external replication in the UKB sample. If such SNPs were located in a risk locus, they are displayed in Locuszoom plots (grey, as there is no LD information in UKB). When risk loci contained GWS SNPs based solely on 23andMe, we did not count that risk locus, as there were no other SNPs available in both samples that supported these GWS SNPs.

### Gene-based analysis

SNP-based *P*-values from the meta-analysis were used as input for the gene-based genome-wide association analysis (GWGAS). 18,182 to 18,185 protein-coding genes (each containing at least one SNP in the GWAS, the total number of tested genes can thus be slightly different across phenotypes) from the NCBI 37.3 gene definitions were used as basis for GWGAS in MAGMA ^23^. Bonferroni correction was applied to correct for multiple testing (*P*<2.73×10^-6^).

### Gene-set analysis

Results from the GWGAS analyses were used to test for association in three types of 7,473 predefined gene-sets:

1. 7,246 curated gene-sets representing known biological and metabolic pathways derived from 9 data resources, catalogued by and obtained from the MsigDB version 6.0^56^ (http://software.broadinstitute.org/gsea/msigdb/collections.jsp)
2. Gene expression values from 54 (53 + 1 calculated 1^st^ PC of three tissue subtypes) tissues obtained from GTEx^32^, log2 transformed with pseudocount 1 after winsorization at 50 and averaged per tissue
3. Cell-type specific expression in 173 types of brain cells (24 broad categories of cell types, ‘level 1’ and 129 specific categories of cell types ‘level 2’), which were calculated following the method described in ^38^. Briefly, brain cell-type expression data was drawn from single-cell RNA sequencing data from mouse brains. For each gene, the value for each cell-type was calculated by dividing the mean Unique Molecular Identifier (UMI) counts for the given cell type by the summed mean UMI counts across all cell types. Single-cell gene-sets were derived by grouping genes into 40 equal bins based on specificity of expression. Mouse cell gene-expression was shown to closely approximate gene-expression in post-mortem human tissue^38^.

These gene-sets were tested using MAGMA. We computed competitive *P*-values, which represent the test of association for a specific gene-set compared with genes not in the gene-set to correct for baseline level of genetic association in the data^57^. The Bonferroni-corrected significance threshold was 0.05/7,473 gene-sets=6.7×10^-6^. Conditional analyses were performed as a follow-up using MAGMA to test whether each significant association observed was independent of all others. The association between each gene-set in each of the three categories was tested conditional on the most strongly associated set, and then, if any substantial (*P*<0.05/number of gene-sets) associations remained, by conditioning on the first and second most strongly associated set, and so on until no associations remained. Gene-sets that retained their association after correcting for other sets were considered to represent independent signals. We note that this is not a test of association per se, but rather a strategy to identify, among gene-sets with known significant associations and overlap in genes, which set (s) are responsible for driving the observed association.

### SNP-based heritability and genetic correlation

LD Score regression^16^ was used to estimate genomic inflation and SNP-based heritability of the phenotypes, and to estimate the cross-cohort genetic correlations. Pre-calculated LD scores from the 1000 Genomes European reference population were obtained from https://data.broadinstitute.org/alkesgroup/LDSCORE/.

### Genetic correlations

Genetic correlations between sleep-related traits, and between sleep-related traits and previously published GWAS studies of sufficient sample size were calculated using LD Score regression on HapMap3 SNPs only. Genetic correlations were corrected for multiple testing based on the total number of correlations (between 6 sleep-related phenotypes and 27 previous GWAS studies) by applying a Bonferroni corrected threshold of (*P<*0.05/33=1.51×10^-3^).

### Stratified heritability

To test whether specific categories of SNP annotations were enriched for heritability, we partitioned SNP heritability for binary annotations using stratified LD score regression^58^. Heritability enrichment was calculated as the proportion of heritability explained by a SNP category divided by the proportion of SNPs that are in that category. Partitioned heritability was computed by 28 functional annotation categories, by minor allele frequency (MAF) in six percentile bins and by 22 chromosomes. Annotations for binary categories of functional genomic characteristics (e.g. coding or regulatory regions) were obtained from the LD score website (https://github.com/bulik/ldsc). The Bonferroni-corrected significance threshold for 56 annotations was set at: *P*<0.05/56=8.93×10^-4^.

### Functional annotation of SNPs

Functional annotation of SNPs implicated in the meta-analysis was performed using FUMA^17^. We selected all candidate SNPs in genomic risk loci having an *r*^2^?0.6 with one of the independent significant SNPs (see above), a *P*-value (*P*<1×10^-5^), a MAF>0.0001 for annotations, and availability in both UKB and 23andMe datasets. Functional consequences for these SNPs were obtained by matching SNPs’ chromosome, base-pair position, and reference and alternate alleles to databases containing known functional annotations, including ANNOVAR^59^ categories, Combined Annotation Dependent Depletion (CADD) scores, RegulomeDB^20^ (RDB) scores, and chromatin states^60^. ANNOVAR categories identify the SNP’s genic position (e.g. intron, exon, intergenic) and associated function. CADD scores predict how deleterious the effect of a SNP is likely to be for a protein structure/function, with higher scores referring to higher deleteriousness. A CADD score above 12.37 is considered to be potentially pathogenic^20^. The RegulomeDB score is a categorical score based on information from expression quantitative trait loci (eQTLs) and chromatin marks, ranging from 1a to 7 with lower scores indicating an increased likelihood of having a regulatory function. Scores are as follows: 1a=eQTL + Transciption Factor (TF) binding + matched TF motif + matched DNase Footprint + DNase peak; 1b=eQTL + TF binding + any motif + DNase Footprint + DNase peak; 1c=eQTL + TF binding + matched TF motif + DNase peak; 1d=eQTL + TF binding + any motif + DNase peak; 1e=eQTL + TF binding + matched TF motif; 1f=eQTL + TF binding / DNase peak; 2a=TF binding + matched TF motif + matched DNase Footprint + DNase peak; 2b=TF binding + any motif + DNase Footprint + DNase peak; 2c=TF binding + matched TF motif + DNase peak; 3a=TF binding + any motif + DNase peak; 3b=TF binding + matched TF motif; 4=TF binding + DNase peak; 5=TF binding or DNase peak; 6=other;7=Not available. The chromatin state represents the accessibility of genomic regions (every 200bp) with 15 categorical states predicted by a hidden Markov model based on 5 chromatin marks for 127 epigenomes in the Roadmap Epigenomics Project^61^. A lower state indicates higher accessibility, with states 1-7 referring to open chromatin states. We annotated the minimum chromatin state across tissues to SNPs. The 15-core chromatin states as suggested by Roadmap are as follows: 1=Active Transcription Start Site (TSS); 2=Flanking Active TSS; 3=Transcription at gene 5’ and 3’; 4=Strong transcription; 5= Weak Transcription; 6=Genic enhancers; 7=Enhancers; 8=Zinc finger genes & repeats; 9=Heterochromatic; 10=Bivalent/Poised TSS; 11=Flanking Bivalent/Poised TSS/Enh; 12=Bivalent Enhancer; 13=Repressed PolyComb; 14=Weak Repressed PolyComb; 15=Quiescent/Low.

### Gene-mapping

Genome-wide significant loci obtained by GWAS were mapped to genes in FUMA^17^ using three strategies:

1. Positional mapping maps SNPs to genes based on physical distance (within a 10kb window) from known protein coding genes in the human reference assembly (GRCh37/hg19).
2. eQTL mapping maps SNPs to genes with which they show a significant eQTL association (i.e. allelic variation at the SNP is associated with the expression level of that gene). eQTL mapping uses information from 45 tissue types in 3 data repositories (GTEx^32^, Blood eQTL browser^60^, BIOS QTL browser^62^), and is based on cis-eQTLs which can map SNPs to genes up to 1Mb apart. We used a false discovery rate (FDR) of 0.05 to define significant eQTL associations.
3. Chromatin interaction mapping was performed to map SNPs to genes when there is a three-dimensional DNA-DNA interaction between the SNP region and another gene region. Chromatin interaction mapping can involve long-range interactions as it does not have a distance boundary. FUMA currently contains Hi-C data of 14 tissue types from the study of Schmitt et al ^63^. Since chromatin interactions are often defined in a certain resolution, such as 40kb, an interacting region can span multiple genes. If a SNP is located in a region that interacts with a region containing multiple genes, it will be mapped to each of those genes. To further prioritize candidate genes, we selected only interaction-mapped genes in which one region involved in the interaction overlaps with a predicted enhancer region in any of the 111 tissue/cell types from the Roadmap Epigenomics Project^61^, and the other region is located in a gene promoter region (250bp up and 500bp downstream of the transcription start site and also predicted by Roadmap to be a promoter region). This method reduces the number of genes mapped but increases the likelihood that those identified will indeed have a plausible biological function. We used a *P*-FDR < 1×10^-5^ to define significant interactions, based on previous recommendations^63^, modified to account for the differences in cell lines used here.

### GWAS catalog lookup

We used FUMA to identify SNPs with previously reported (*P*<5×10^-5^) phenotypic associations in published GWAS listed in the NHGRI-EBI catalog^64^, which matched with SNPs in LD with one of the independent significant SNPs identified in the meta-analysis.

### Polygenic risk scoring

To calculate the explained variance in insomnia by our GWAS results, we calculated polygenic scores (PGS) based on the SNP effect sizes in the meta-analysis. The PGS were calculated using two methods: LDpred^65^ and PRSice^66^, a script for calculating *P-*value thresholded PGS in PLINK. PGS were calculated using a leave-one-out method, where summary statistics were recalculated each time with one sample of N=3,000 from UKB excluded from the analysis. This sample was then used as a target sample for estimating the explained variance in insomnia by the PGS.

### Mendelian Randomization

To investigate causal associations between insomnia and genetically correlated traits, we analyzed direction of effects using Generalized Summary-data based Mendelian Randomization (GSMR^43^; http://cnsgenomics.com/software/gsmr/). This method uses effect sizes from GWAS summary statistics (standardized betas or log-transformed odds ratios) to infer causality of effects between two traits based on genome-wide significant SNPs. Built-in HEIDI outlier detection was applied to remove SNPs with pleiotropic effects on both traits, as these may bias the results. We tested for causal associations between insomnia and traits that were significantly genetically correlated with insomnia (*b_zx_*). In addition, we tested for bi-directional associations by using other traits as the determinant and insomnia as the outcome (*b_zy_*). We selected independent (*r*^2^<0.1) lead SNPs with a GWS *P*-value (<5×10^-8^) as instrumental variables in the analyses. For traits with less than 10 lead SNPs (i.e. the minimum number of SNPs on which GSMR can perform a reliable analysis) we selected independent SNPs (*r*^2^<0.1), with a *P*-value *<*1×10^-5^. If the outcome trait is binary, the estimated *b_zx_* and *b_zy_* are approximately equal to the natural log of the odds ratio (OR). An OR of 2 can be interpreted as a doubled risk compared to the population prevalence of a binary trait for every SD increase in the exposure trait. For quantitative traits, the *b_zx_* and *b_zy_* can be interpreted as a one standard deviation increase explained in the outcome trait for every SD increase in the exposure trait.

## Acknowledgements

This work was funded by The Netherlands Organization for Scientific Research (NWO Brain & Cognition 433-09-228, NWO MagW VIDI 452-12-04, NWO VICI 435-14-005 and 453-07-001, 645-000-003). P.R.J. was funded by the Sophia Foundation for Scientific Research (S14-27), E.J.W.V.S. was funded by the European Research Council (ERC-ADG-2014-671084 INSOMNIA), J.B. was funded by the Swiss National Science Foundation. Analyses were carried out on the Genetic Cluster Computer, which is financed by the Netherlands Scientific Organization (NWO: 480-05-003), by the VU University, Amsterdam, the Netherlands, and by the Dutch Brain Foundation, and is hosted by the Dutch National Computing and Networking Services SurfSARA. This research has been conducted using the UK Biobank Resource (application number 16406). We would like to thank the participants and researchers who collected and contributed to the data. We thank the 23andMe research participants and employees for making this work possible, including the following members of the 23andMe Research Team: Michelle Agee, Babak Alipanahi, Adam Auton, Robert K. Bell, Katarzyna Bryc, Sarah L. Elson, Pierre Fontanillas, Nicholas A. Furlotte, David A. Hinds, Karen E. Huber, Aaron Kleinman, Nadia K. Litterman, Jennifer C. McCreight, Matthew H. McIntyre, Joanna L. Mountain, Elizabeth S. Noblin, Carrie A. M. Northover, Steven J. Pitts, J. Fah Sathirapongsasuti, Olga V. Sazonova, Janie F. Shelton, Suyash Shringarpure, Chao Tian, and Catherine H. Wilson.

## Author contributions

D.P. and E.J.W.V.S. conceived the idea of the study. D.P. supervised the pre- and post gwas analysis pipeline. P.R.J. and K.W. performed the analyses. S.St. performed quality control on the UK Biobank data and wrote the analysis pipeline. K.W. wrote the online platform (FUMA) that was used for follow-up analyses. C.d.L conducted conditional gene-set analyses. J.B., N.S., A.M.M. and J.H.L contributed scRNA information. J.T., D.H., V.V. and the 23andMe Research Team contributed and analyzed the 23andMe cohort data. D.P., E.J.W.V.S. and P.R.J. wrote the paper. All authors discussed the results, and approved the final version of the paper.

## Materials & Correspondence

The data analyzed in the current study was partly provided by the UK Biobank Study (www.ukbiobank.ac.uk), received under the UK Biobank application number 16406. Our policy is to make genome-wide summary statistics (sumstats) publicly available. Sumstats from the GWAS’s conducted are available for download at https://ctg.cncr.nl/. Note that our freely available meta-analytic sumstats (insomnia and morningness) concern results excluding the 23andMe sample. This is a non-negotiable clause in the 23andMe data transfer agreement, intended to protect the privacy of the 23andMe research participants. To fully recreate our meta-analytic results for insomnia and morningness: (a) obtain insomnia and morningness sumstats from 23andMe (see below); (b) conduct a meta-analysis of our sumstats with the 23andMe sumstats. 23andMe participant data are shared according to community standards that have been developed to protect against breaches of privacy. Currently, these standards allow for the sharing of summary statistics for at most 10,000 SNPs. The full set of summary statistics can be made available to qualified investigators who enter into an agreement with 23andMe that protects participant confidentiality. Interested investigators should for more information.

## Author Information

V.V., D.H., and J.T. are employees of 23andMe. All other authors declare no competing financial interest. Correspondence and requests for materials should be addressed to.

## EXTENDED DATA

**Extended Data Table 1.** LD Score regression estimates of the sex-specific GWAS of insomnia. Results are shown for UK Biobank, 23andMe and the sex-specific meta-analyzed sample. H2=estimated SNP-heritability, intercept=LD Score regression intercept, rg=genetic correlation in the same study sample.

| Sample | Sex | N | *h*<sup>2</sup>(SE) | Mean chi<sup>2</sup> | Lambda | Intercept (SE) | *rg* male | *rg* female |
| --- | --- | --- | --- | --- | --- | --- | --- | --- |
| UK Biobank | male | 177.817 | 0.083<br>(0.007) | 1,157 | 1,143 | 1.001<br>(0.008) | 1 | 0.857<br>(0.051) |
|  | female | 208.716 | 0.092<br>(0.005) | 1,233 | 1,210 | 1.011<br>(0.008) | 0.857<br>(0.051) | 1 |
| 23andMe | male | 443.207 | 0.080<br>(0.004) | 1,385 | 1,317 | 1.016<br>(0.008) | 1 | 0.925<br>(0.022) |
|  | female | 501.270 | 0.090<br>(0.003) | 1,580 | 1,460 | 1.046<br>(0.009) | 0.925<br>(0.022) | 1 |
| Meta | male | 621.024 | 0.067<br>(0.003) | 1,460 | 1,382 | 1.024<br>(0.009) | 1 | 0.919<br>(0.018) |
|  | female | 709.986 | 0.078<br>(0.003) | 1,700 | 1,547 | 1.042<br>(0.009) | 0.919<br>(0.018) | 1 |

**Extended Data Fig 1a-b.**
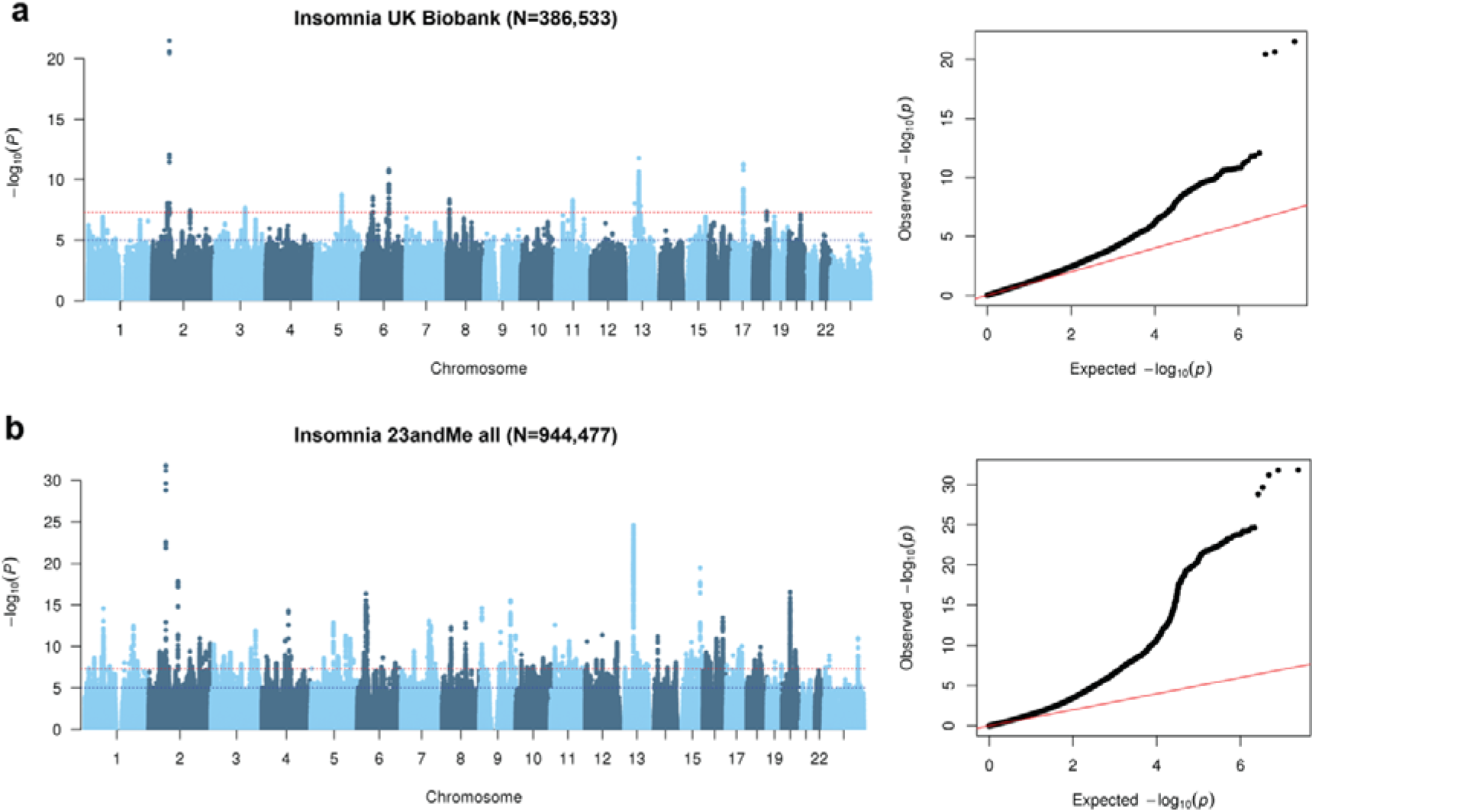
Manhattan and Q-Q plots of the genome-wide analysis of insomnia. Results are shown for the genome-wide analysis in (**a**) UK Biobank and (**b**) 23andMe.

**Extended Data Fig. 3a-c.**
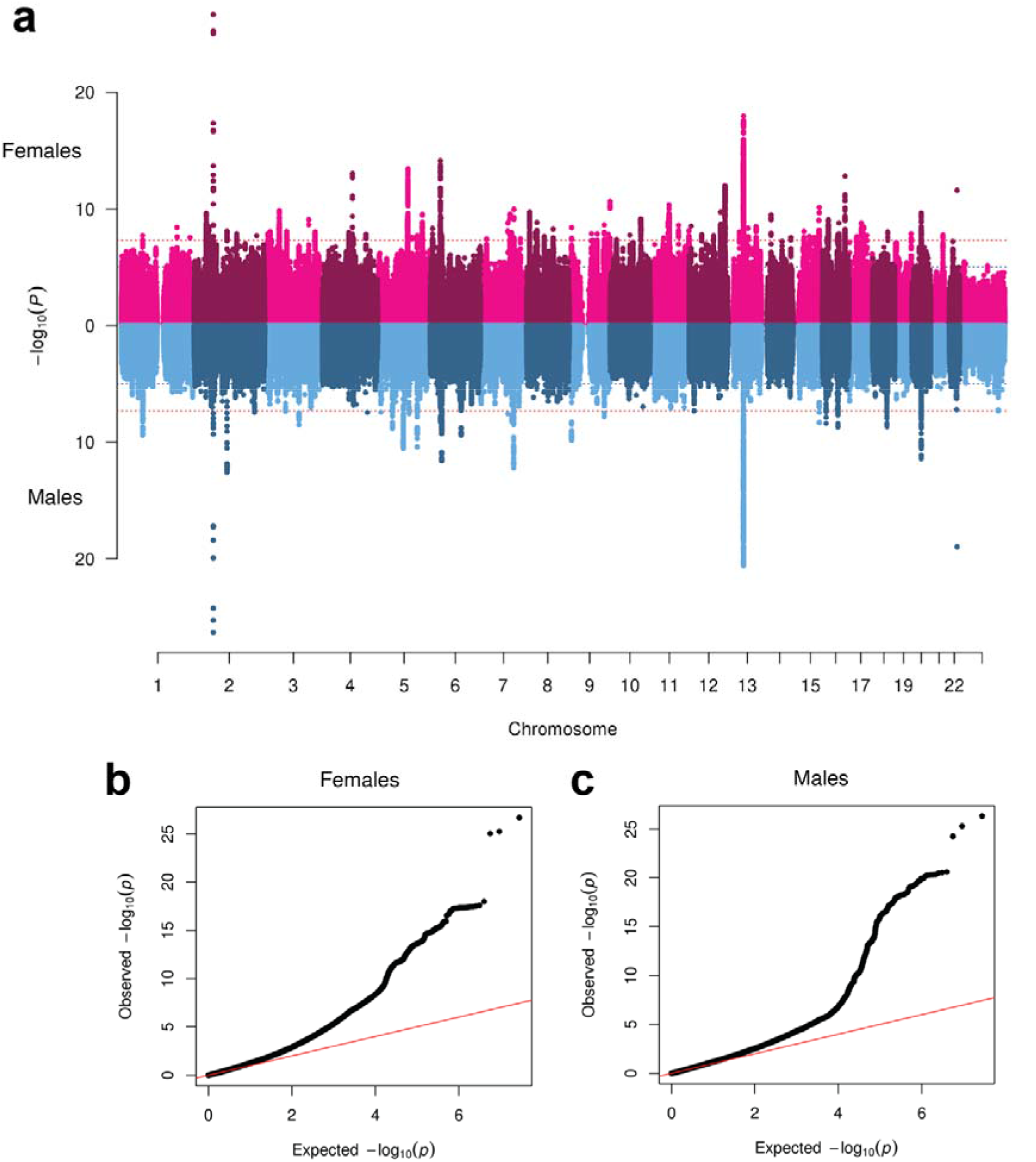
Sex-specific Manhattan plot and Q-Q plot of the insomnia meta-analysis in males and females (UK Biobank + 23andMe). (**a**) Miami plot showing sex-specific SNP association P-values for females on the upper side and males on the lower side. (**b**) Q-Q plot in females, and (**c**) in males.

**Extended Data Fig. 4a-b.**
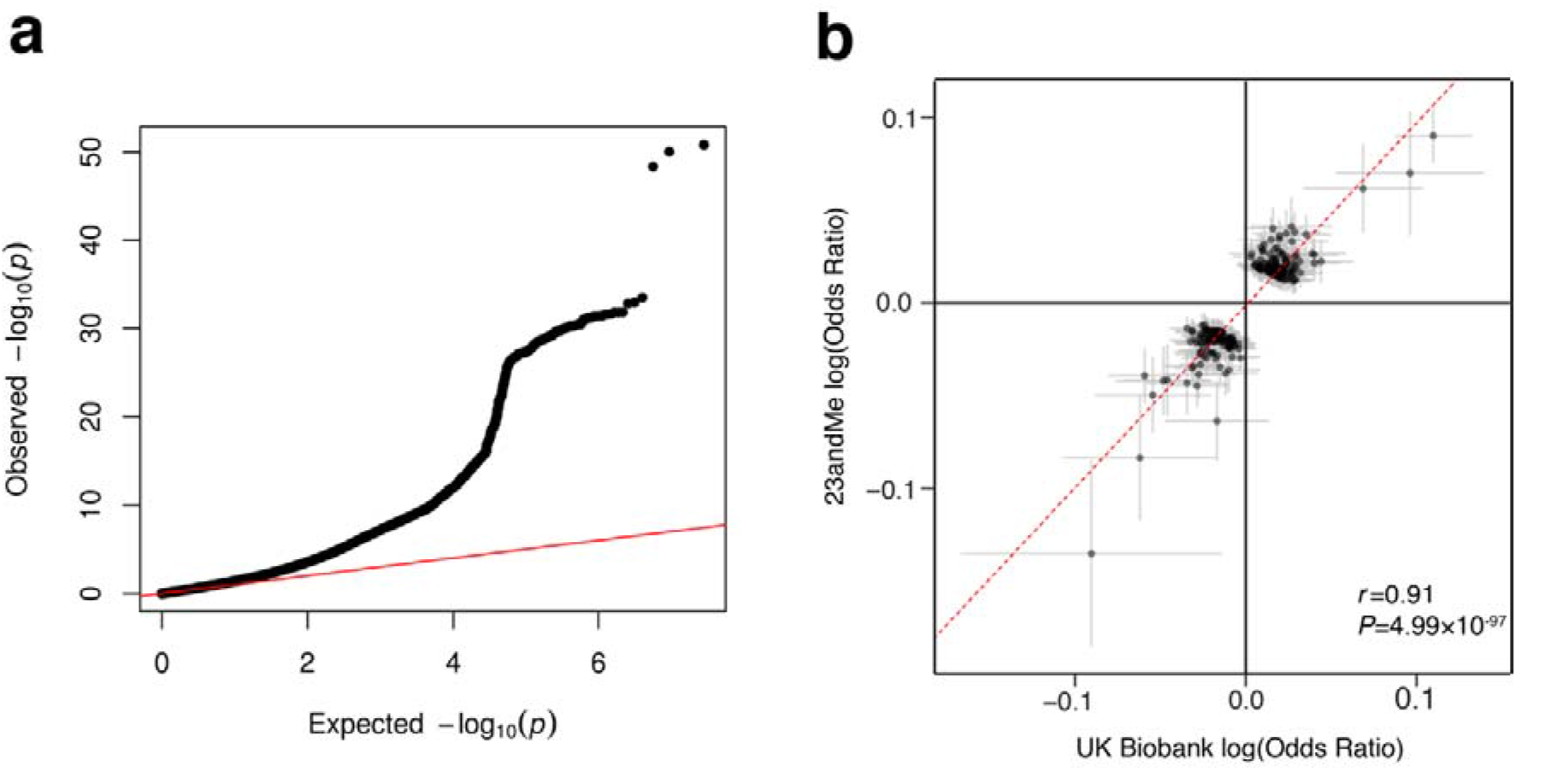
Q-Q plot and lead SNPs of the GWAS meta-analysis for insomnia. (**a**) QQ-plot of the insomnia meta-analysis showing the expected negative log10-transformed *P*-value distribution on the x-axis, and observed negative log10-transformed *P*-value on the y-axis, (**b**) effect size plot of the 248 lead SNP of the insomnia meta-analysis (log-transformed odds ratio and 95% confidence interval) in UK Biobank and 23andMe.

**Extended Data Fig. 5.**
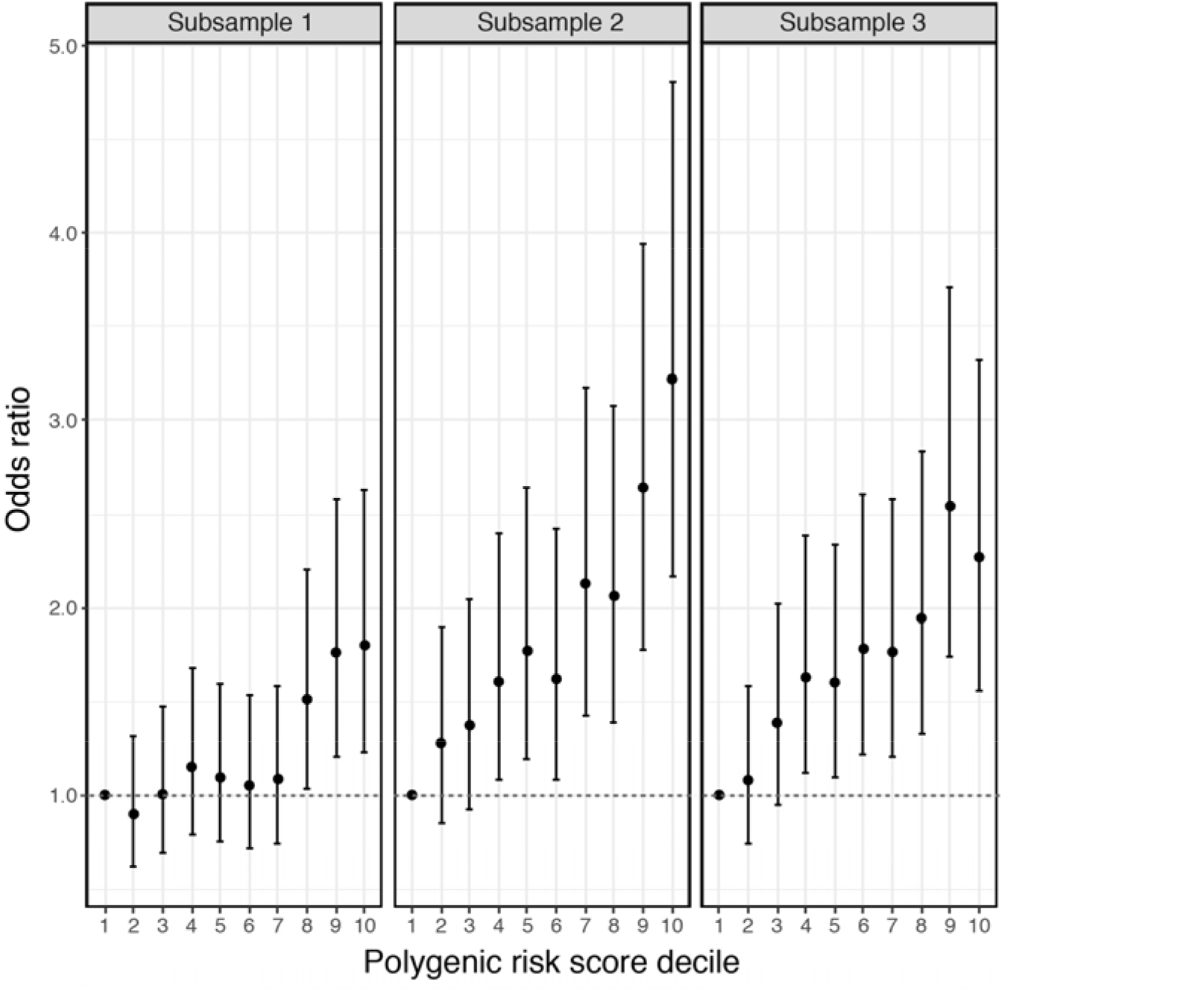
Risk of insomnia per polygenic risk score decile in three independent holdout samples (N=3×3000). Odds ratios and 95% confidence interval for deciles in polygenic risk score were calculated based on a logistic regression model, using the lowest polygenic risk score decile as the reference.

**Extended Data Fig. 6.**
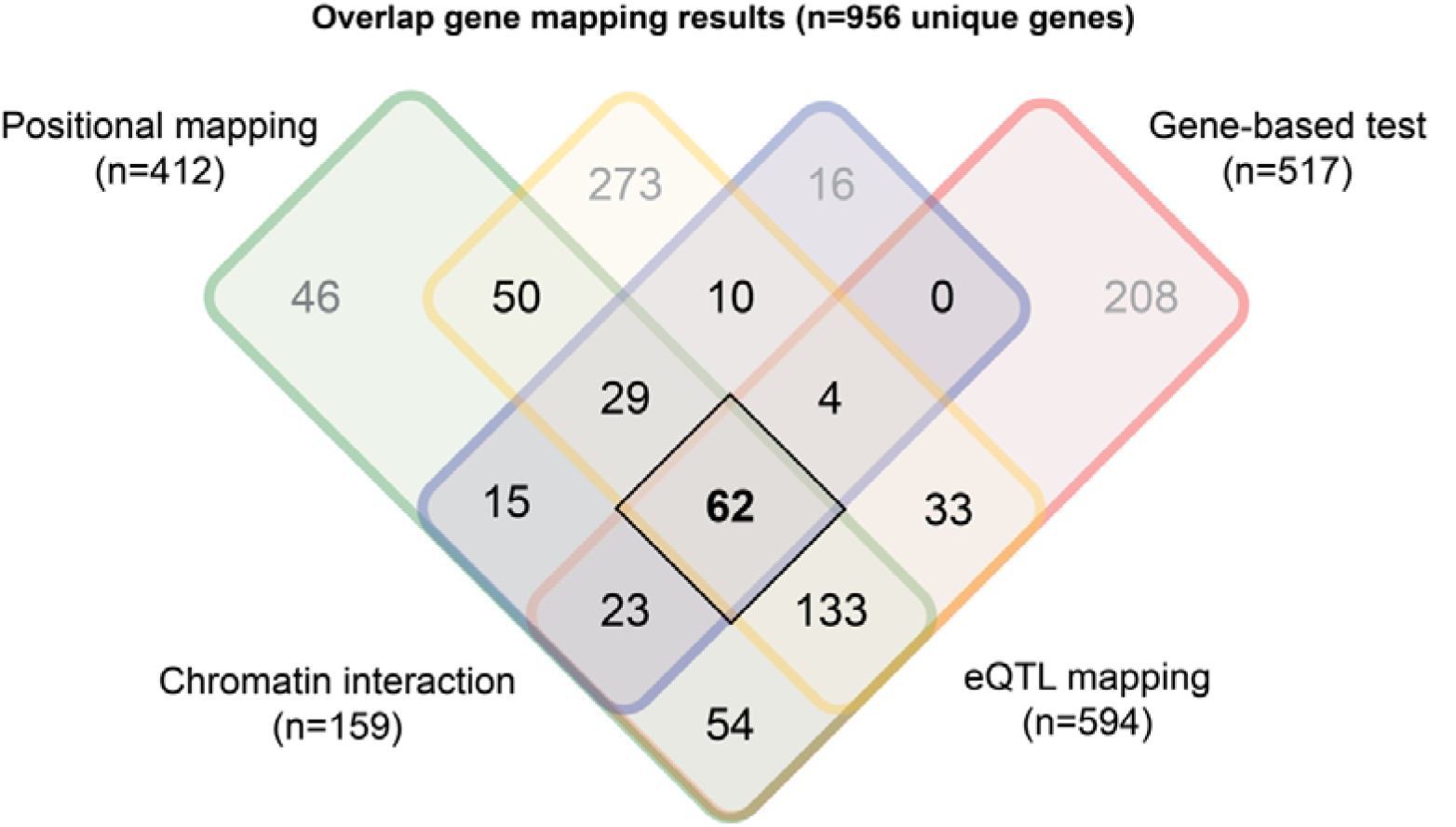
Venn diagram showing the number of genes that were mapped by four gene-mapping strategies. Each square shows the number of overlapping genes between three gene-mapping methods in FUMA (positional mapping, eQTL mapping and chromatin interaction mapping) and significant genes in gene-based tests in MAGMA. The number of genes in bold highlights the number of genes that were implicated by all four methods.

**Extended Data Fig. 7a-b.**
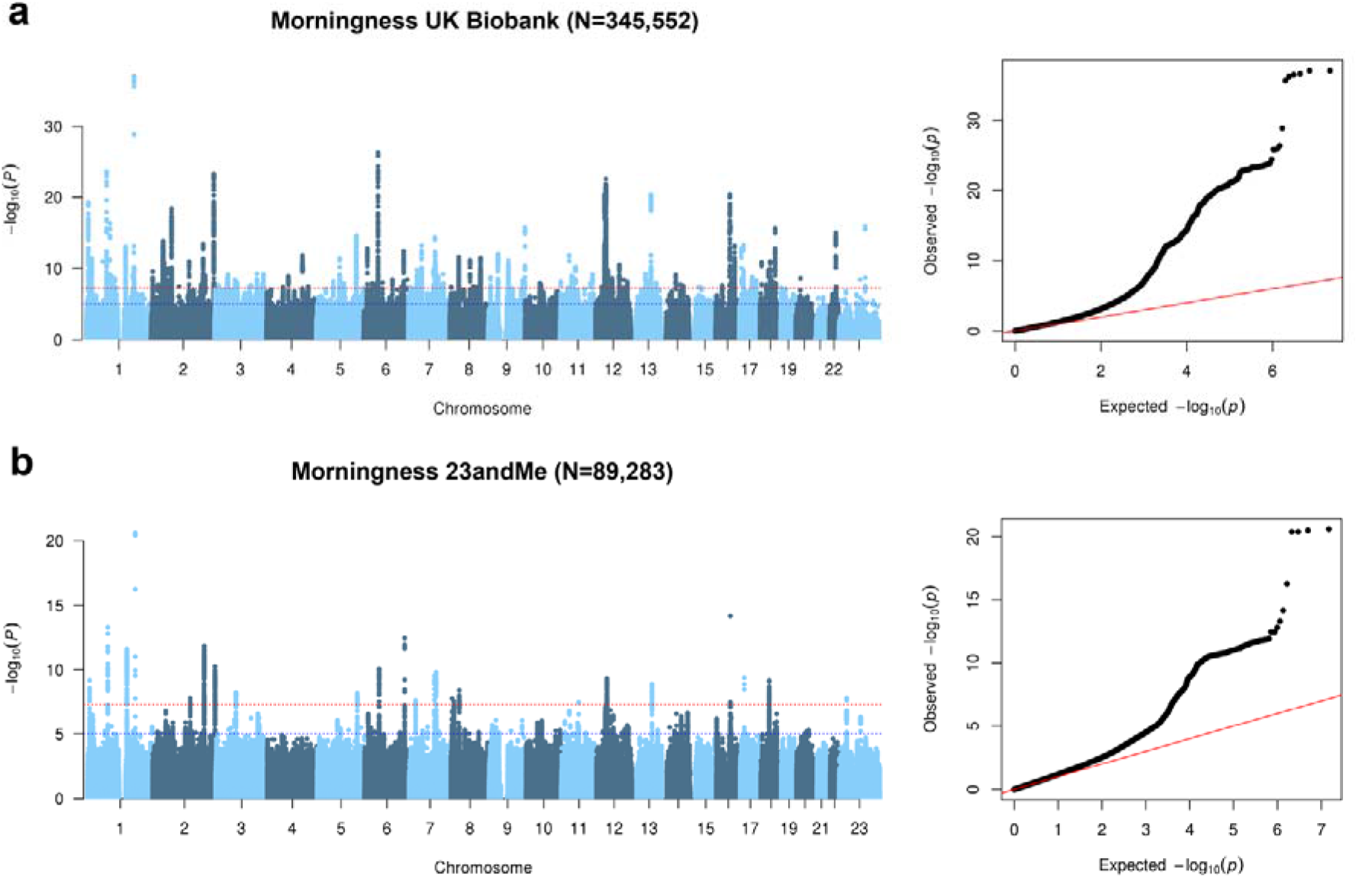
Manhattan plot and Q-Q plot of the genome-wide analysis of morningness in UK Biobank and 23andMe. Results are shown for (**a**) UK Biobank and (**b**) 23andMe.

**Extended Data Fig. 8a-f.**
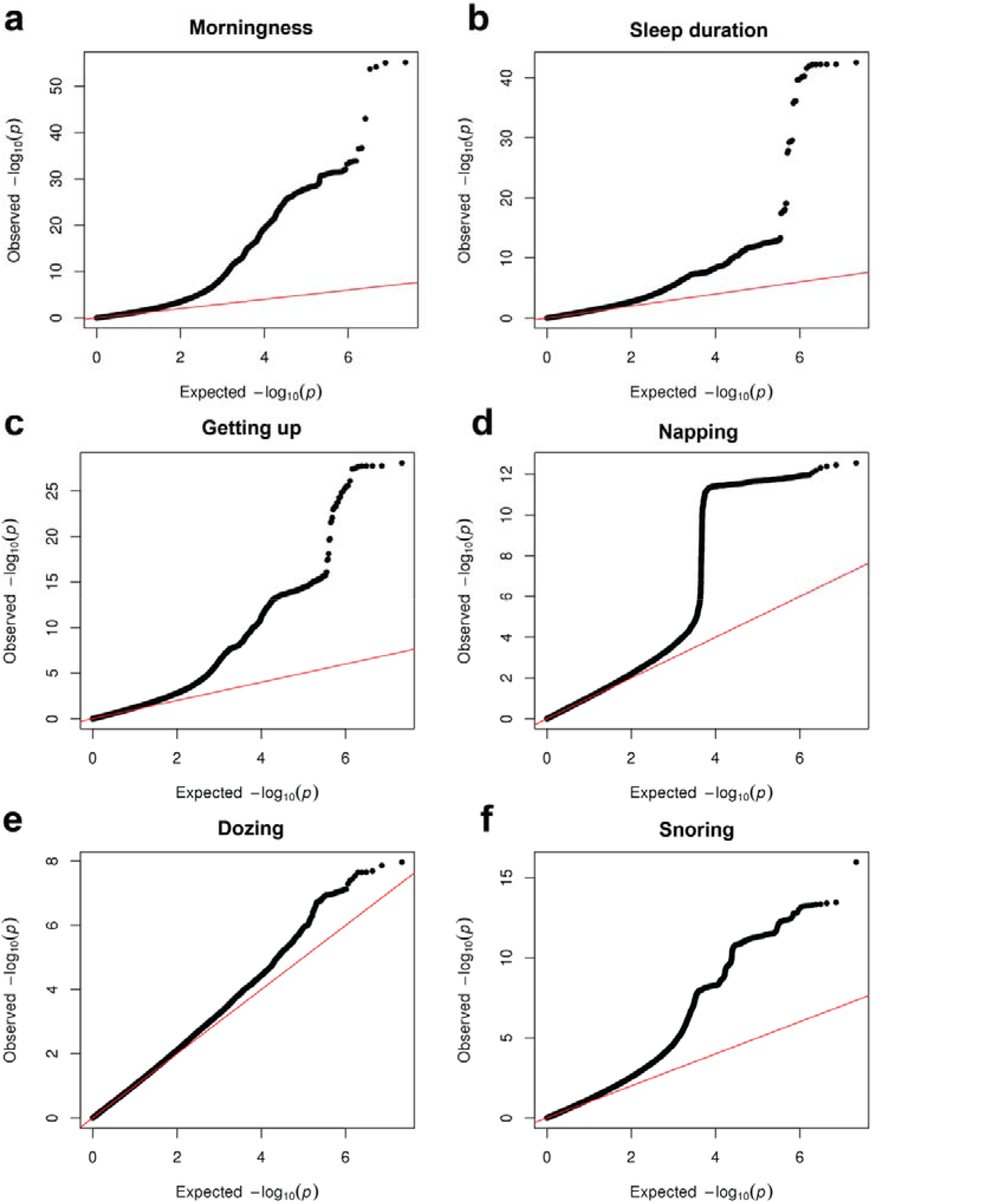
Q-Q plots of the genome-wide analysis of six sleep related traits. (**a**) morningness (including UKB and 23andMe), (**b**) sleep duration, (**c**) ease of getting up, (**d**) daytime napping, (**e**) daytime dozing, (**f**) snoring. Manhattan plots of the genome-wide analyses are shown in Fig. 3.

**Extended Data Fig. 9a-f.**
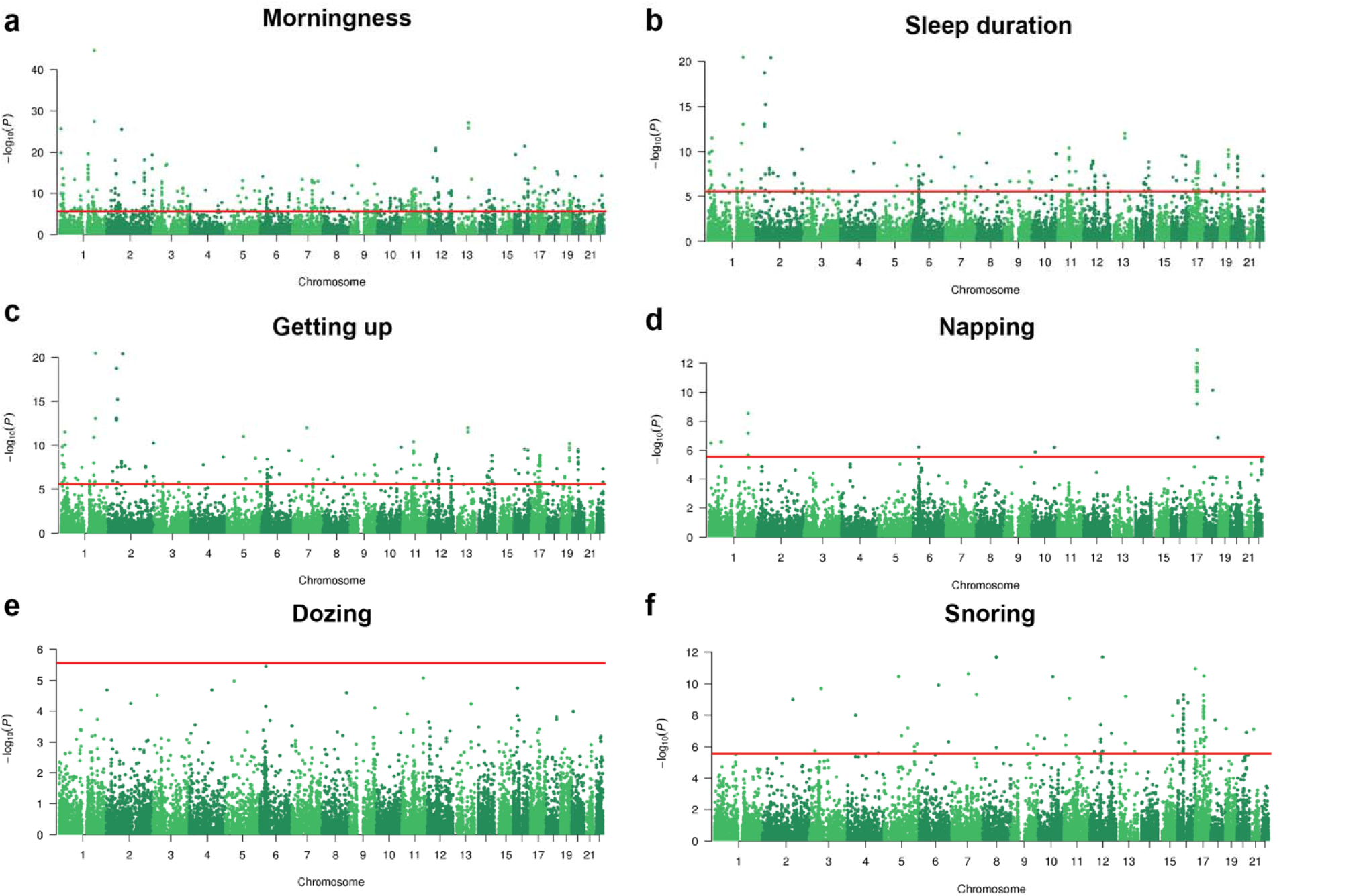
Genome-wide gene-based association analysis of six sleep-related phenotypes. Manhattan plots genome-wide gene-based analysis (GWGAS) results for (**a**) morningness (**b**) sleep duration (**c**) ease of getting up (**d**) daytime napping (**e**) daytime dozing (**f**) snoring. GWGAS was performed in MAGMA. The analysis of morningness was based on GWAS meta-analysis of UKB and 23andMe, while other sleep-related phenotypes were analysed in UKB. The red line indicates Bonferroni corrected significance threshold depending on the number of genes tested.

Supplementary Information includes:

**1. Supplementary Methods**

1.1 Sample description UK Biobank

1.2 Sample description 23andMe

1.3 Insomnia phenotype validation external sample

**2. Supplementary Discussion**

2.1. Sex-specific association results for insomnia

2.2. GWAS meta-analysis results for insomnia

2.3. Implicated genes for insomnia

2.4. Gene-set association results for insomnia

2.5 Results sleep-related phenotypes

2.6 Mendelian Randomization

**3. Supplementary Figures (1 to 2**)

**4. Supplementary Tables (1 to 28**)

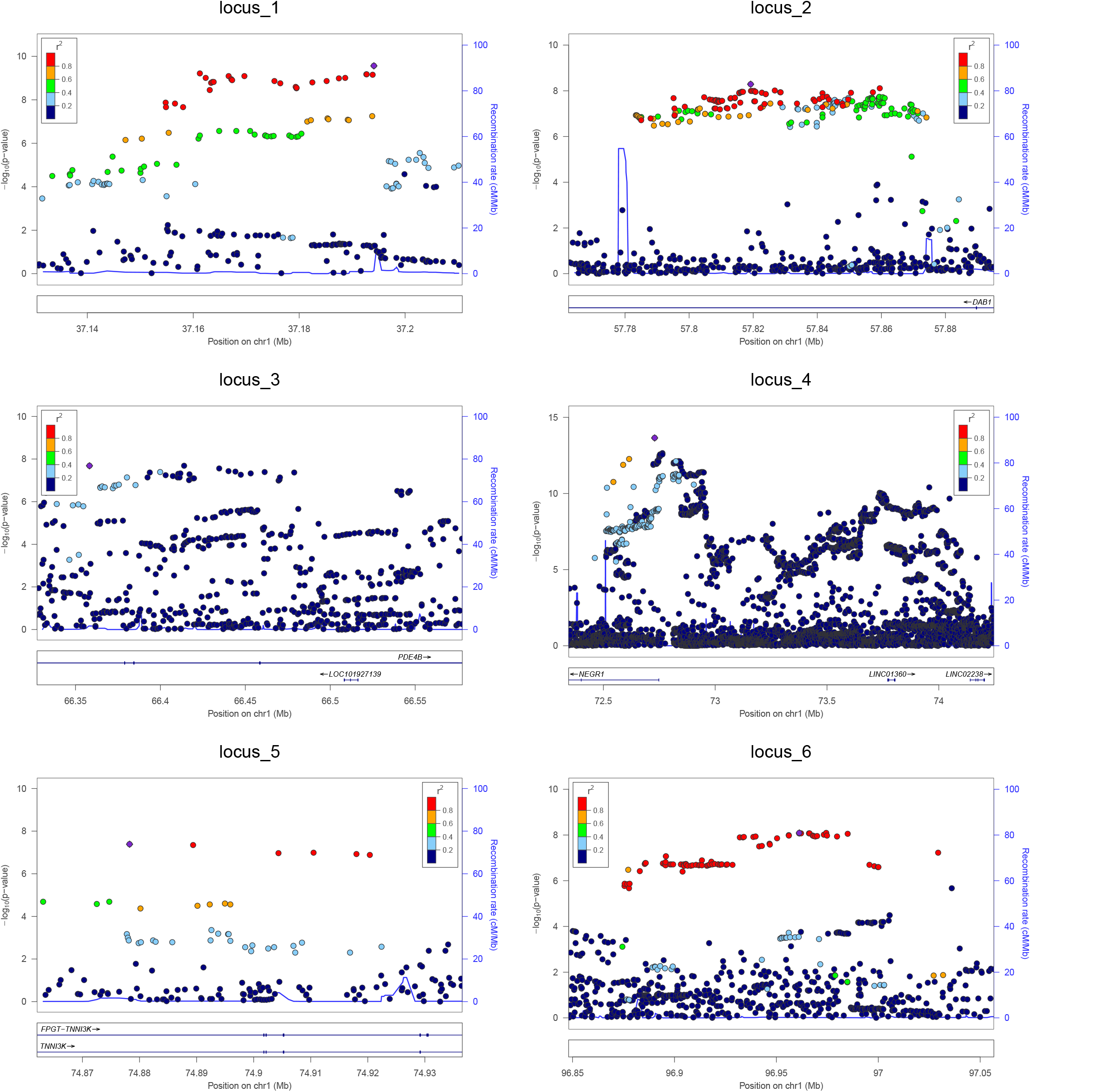
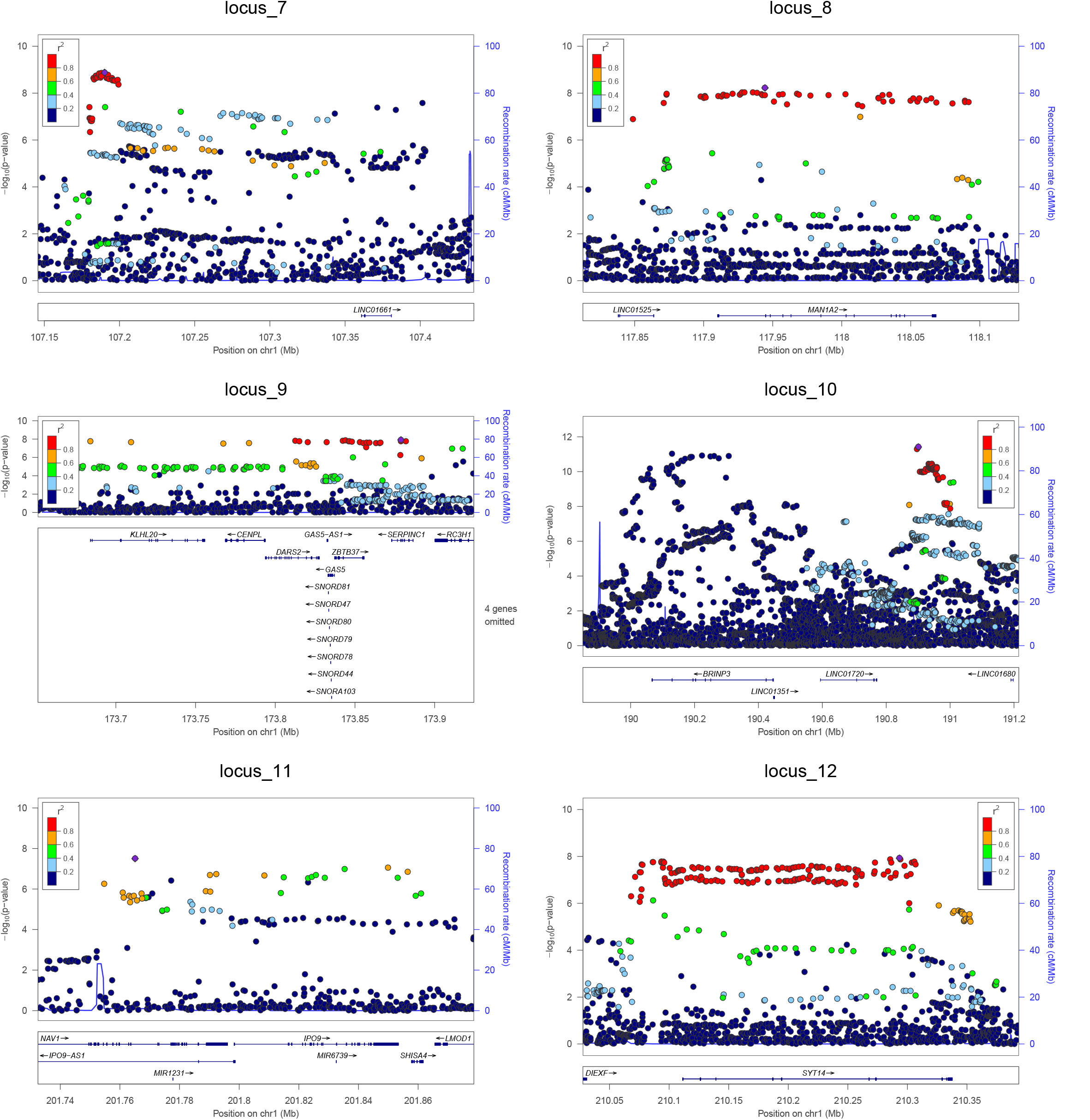
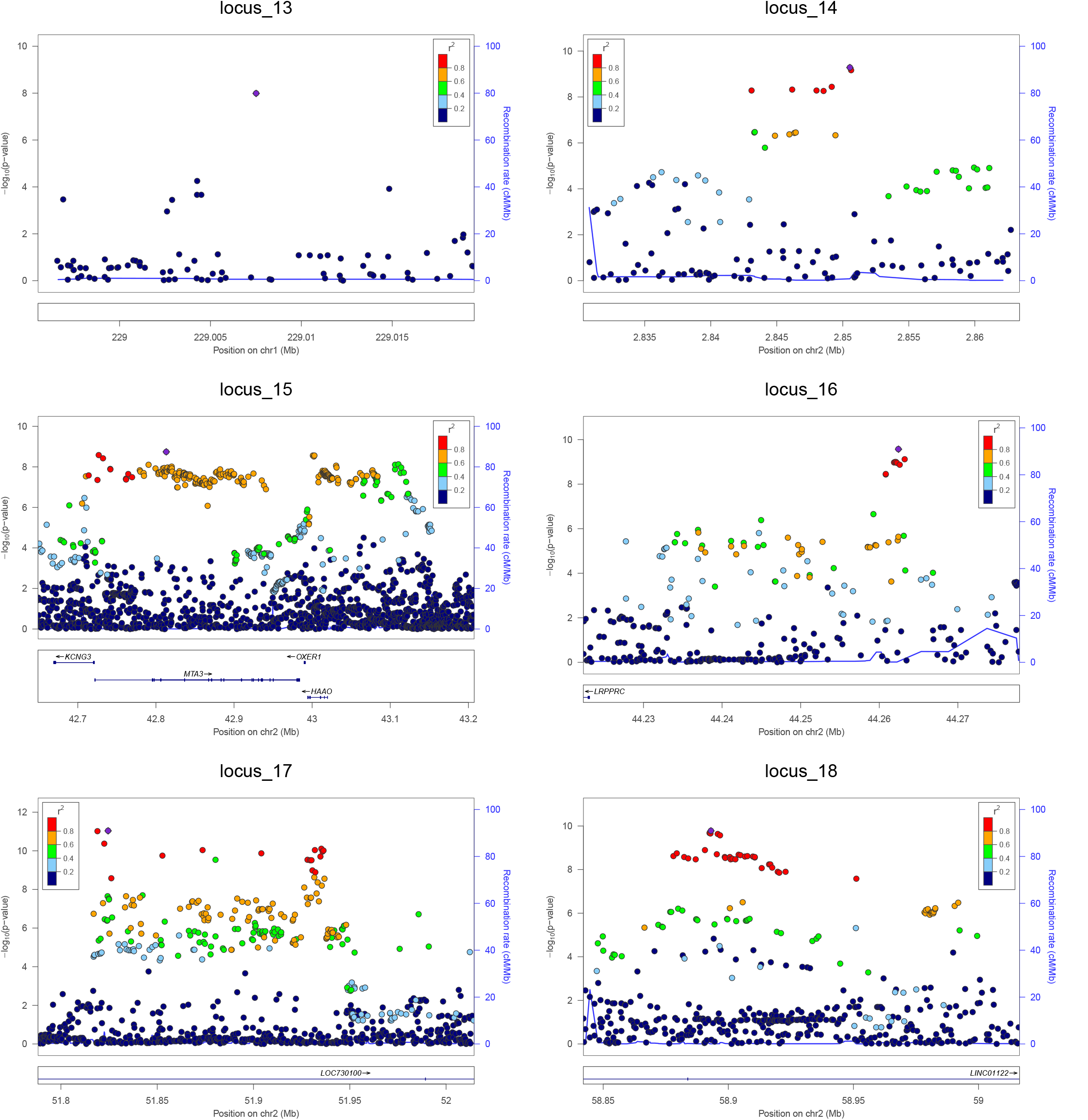
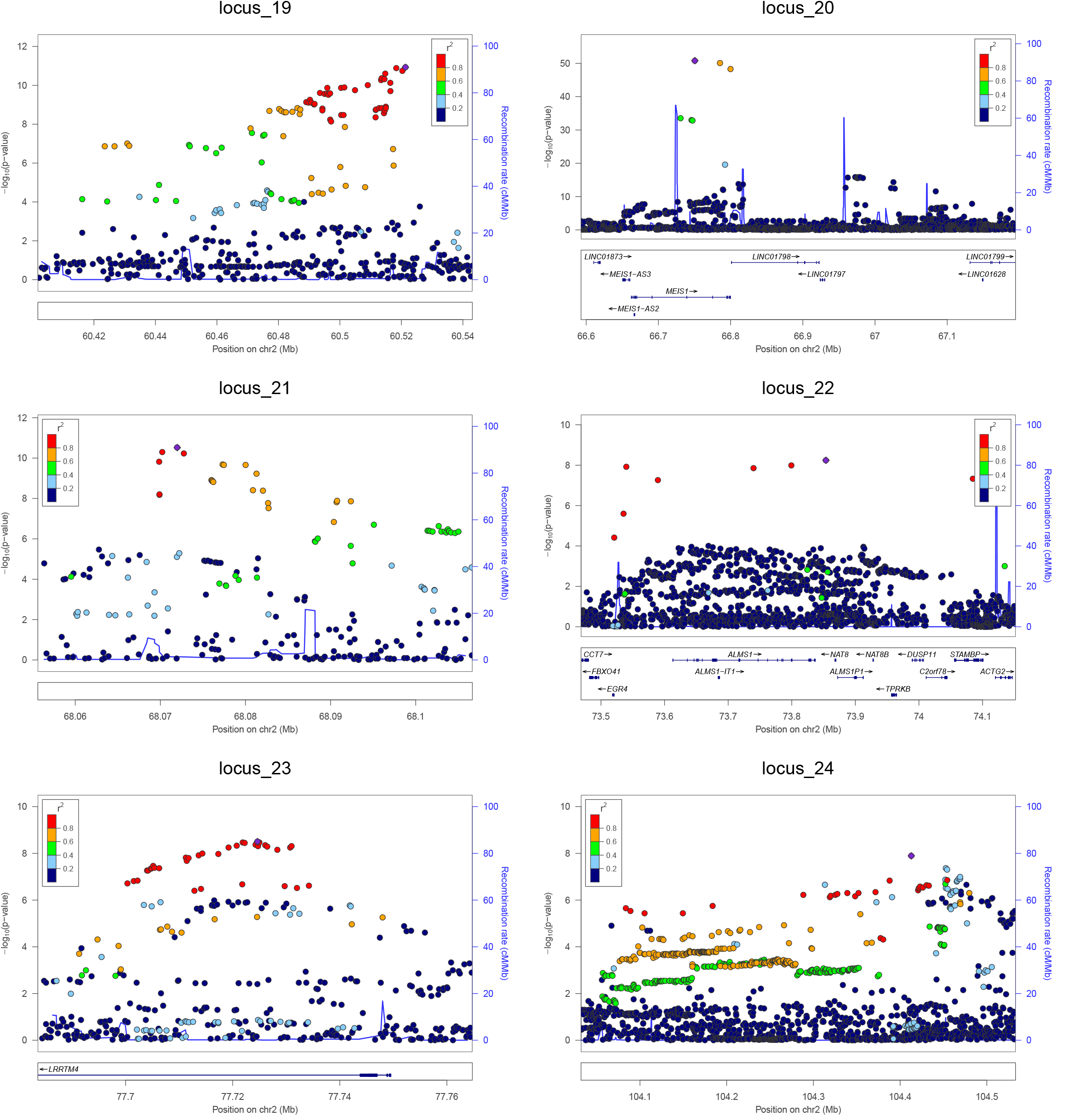
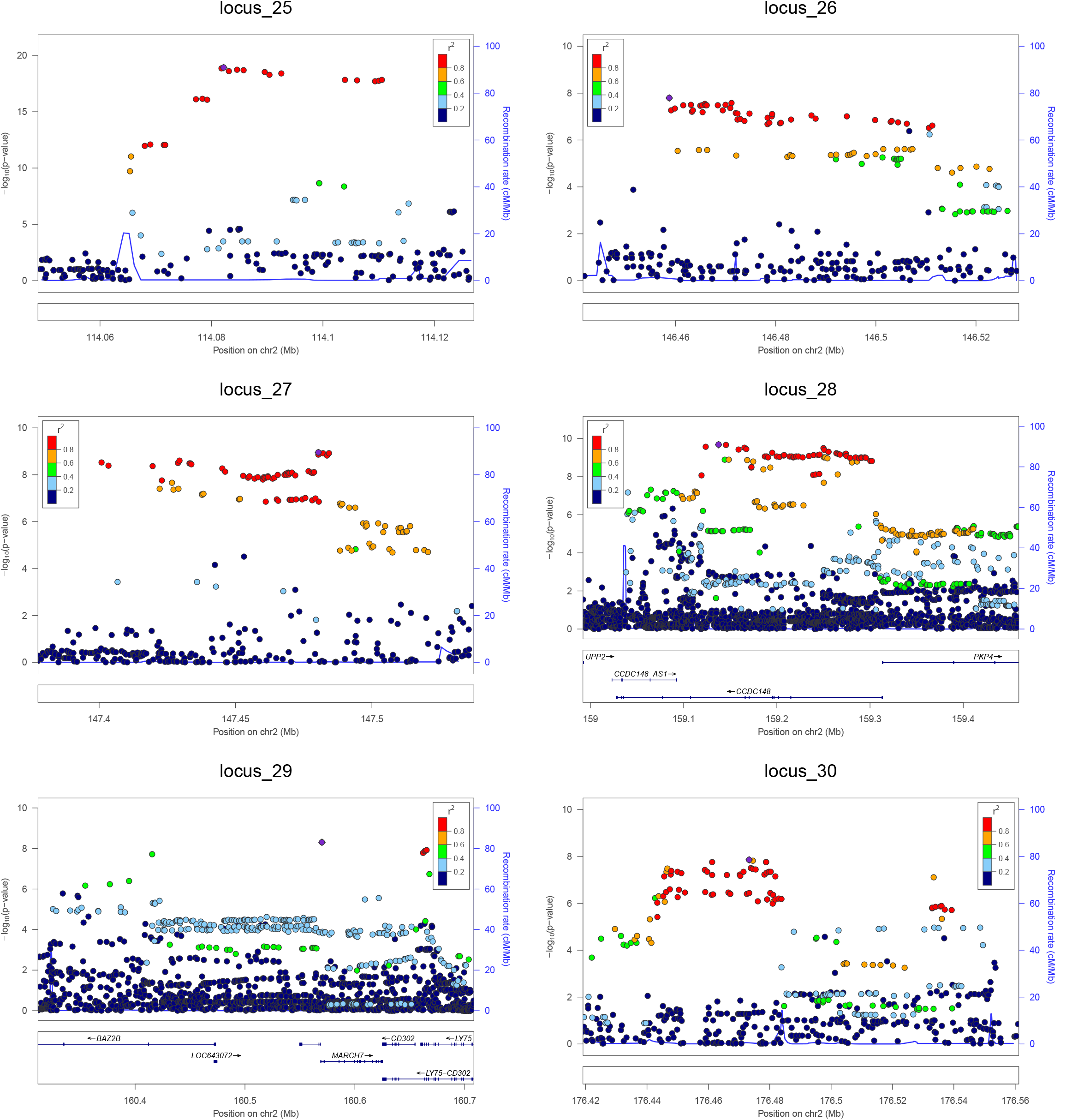
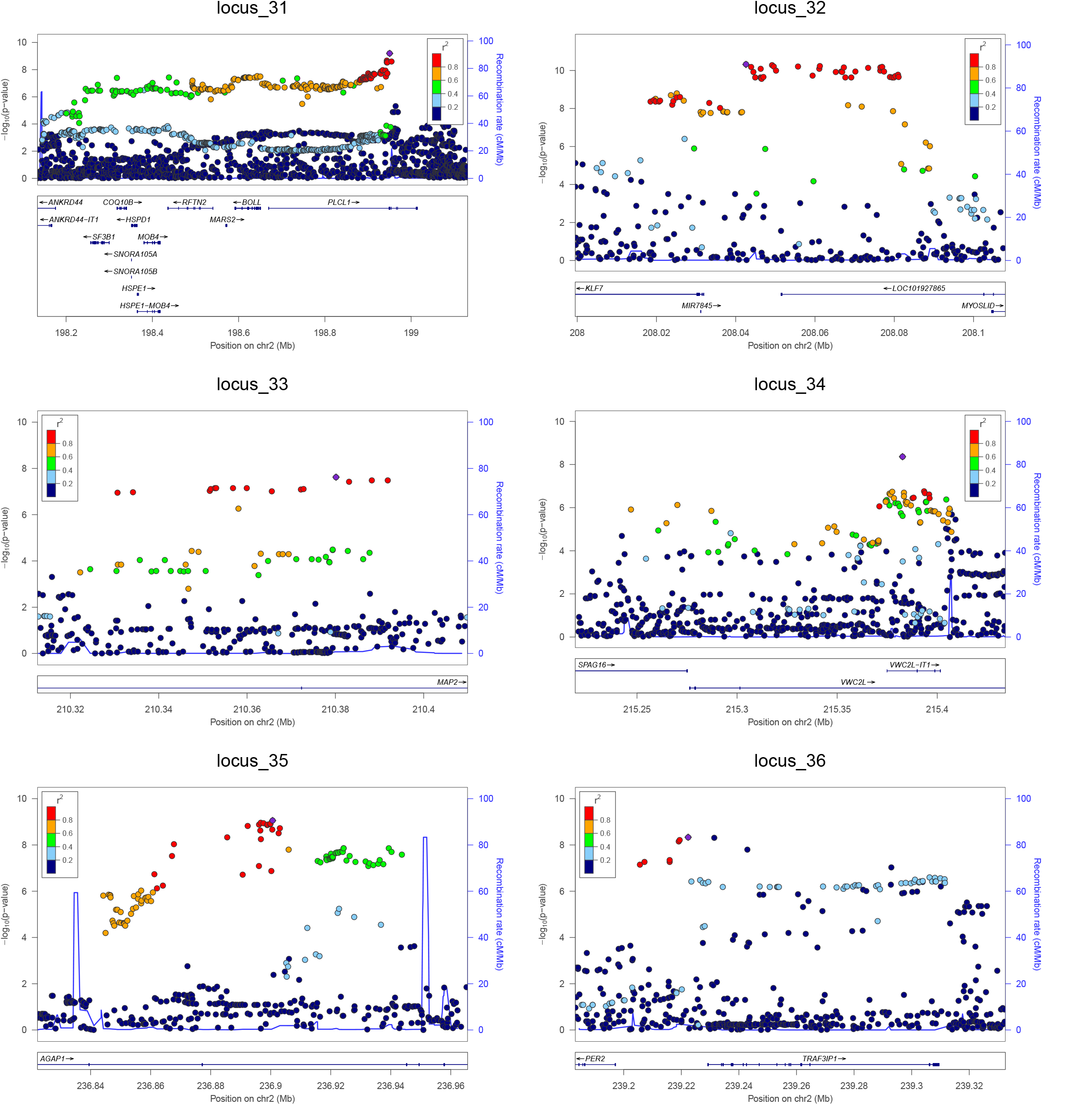
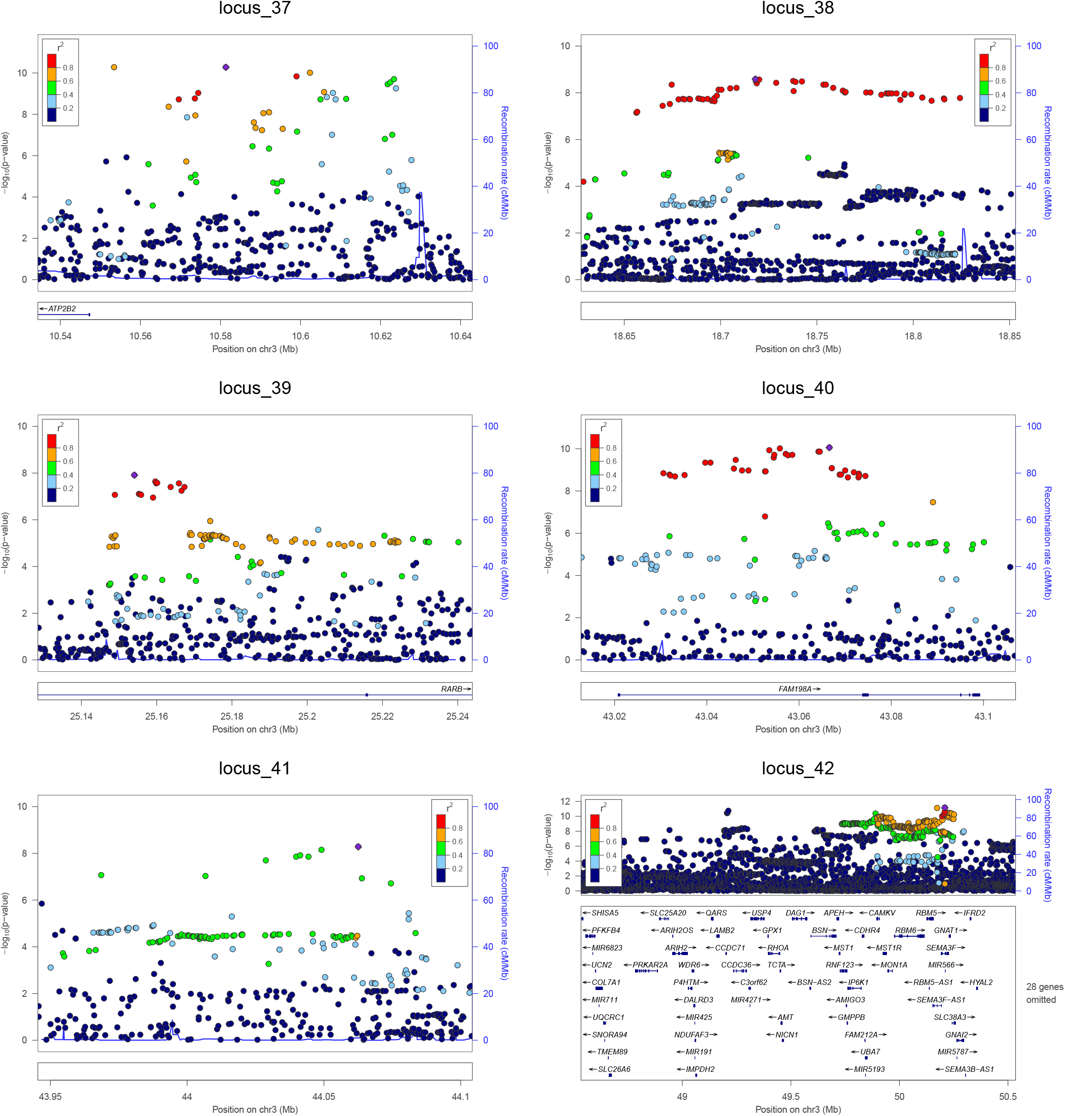
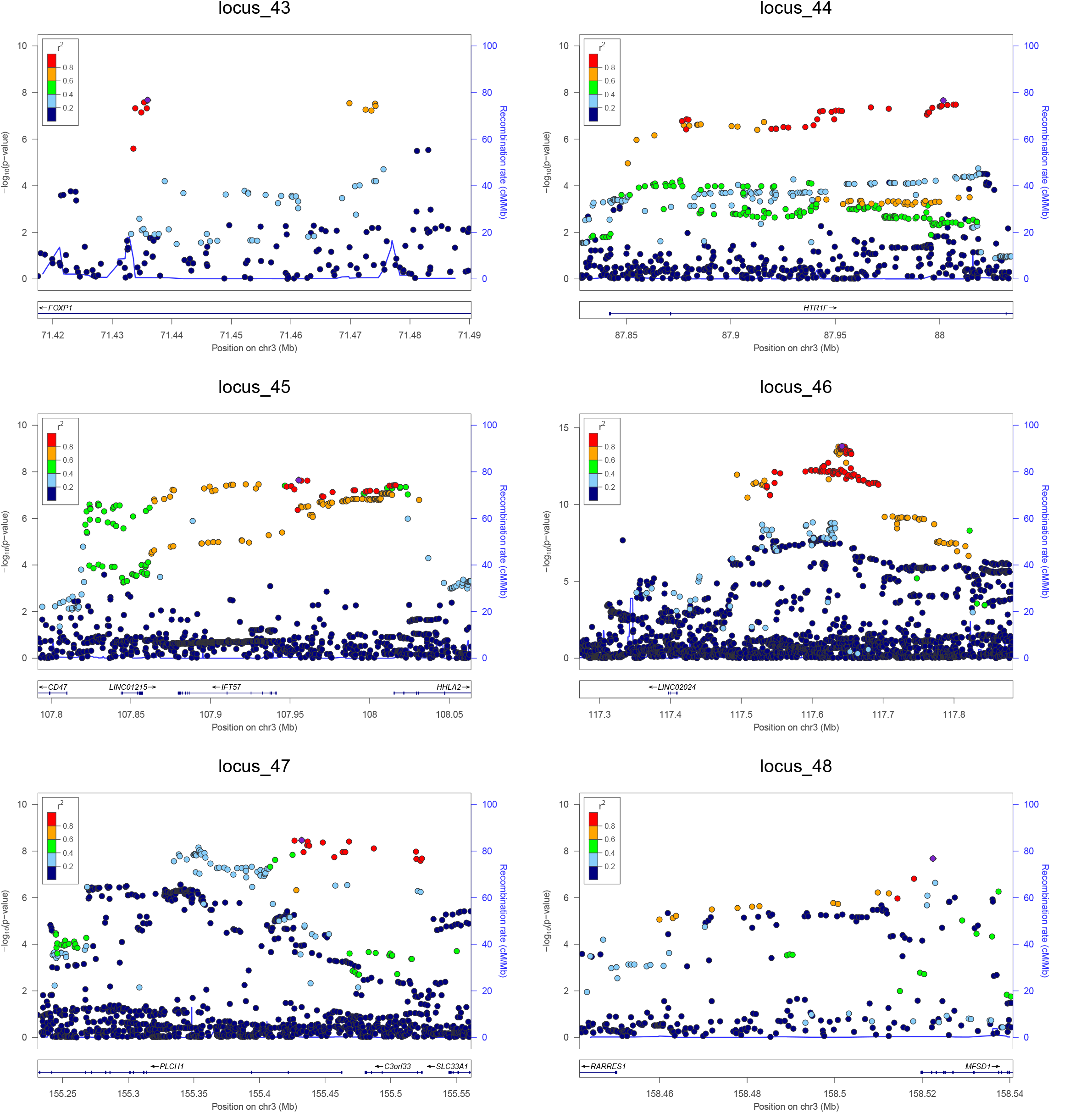
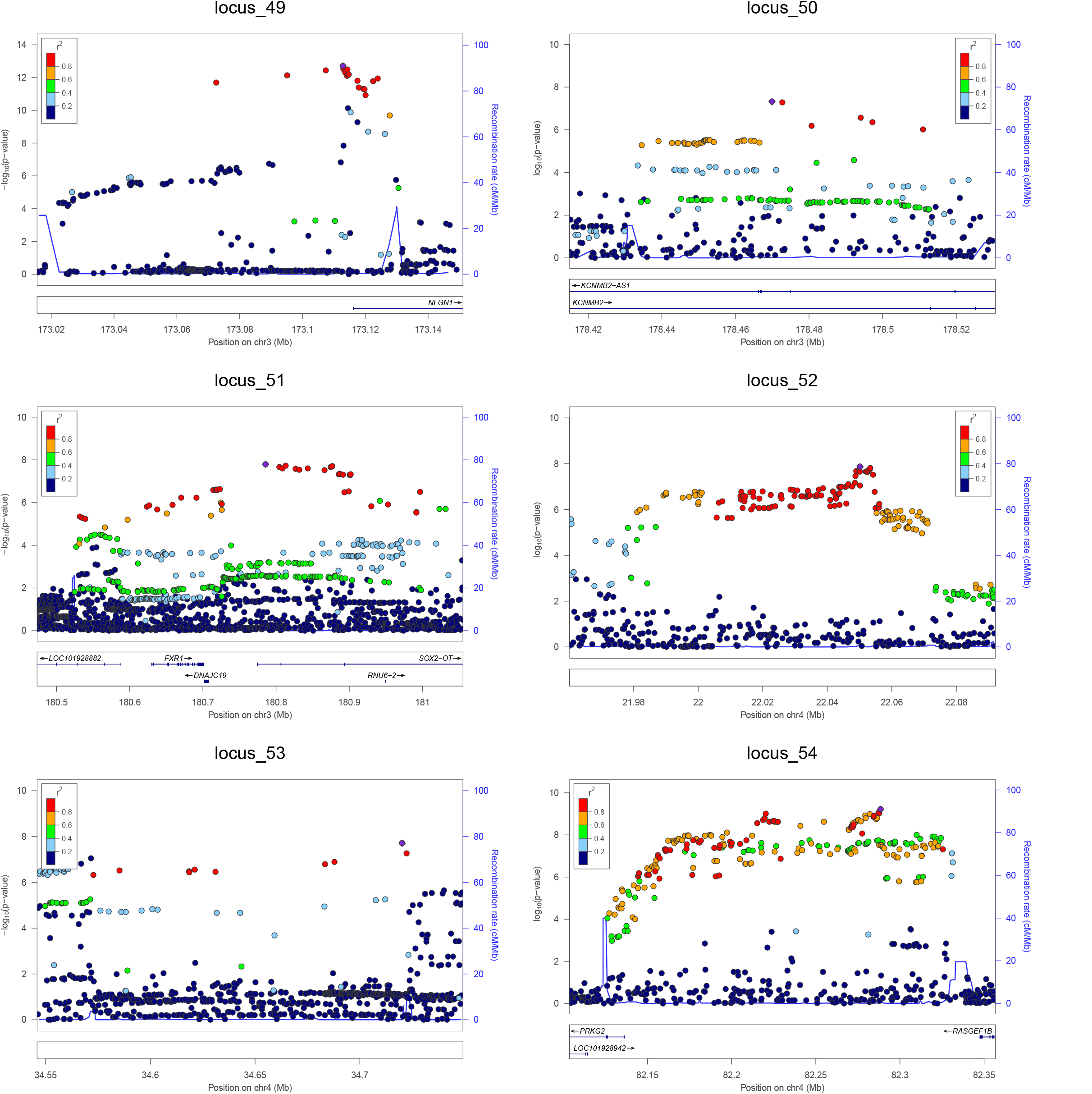
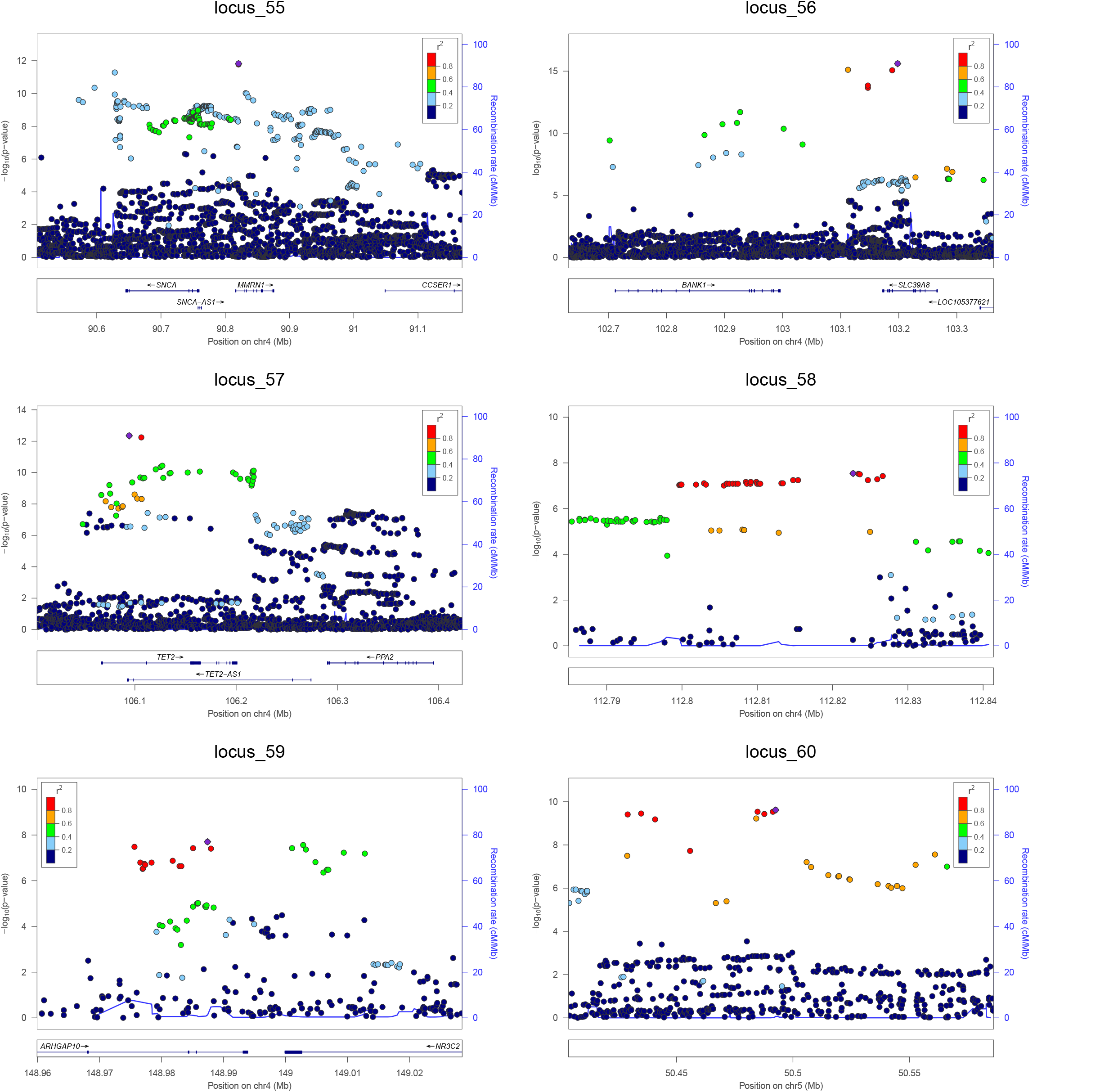
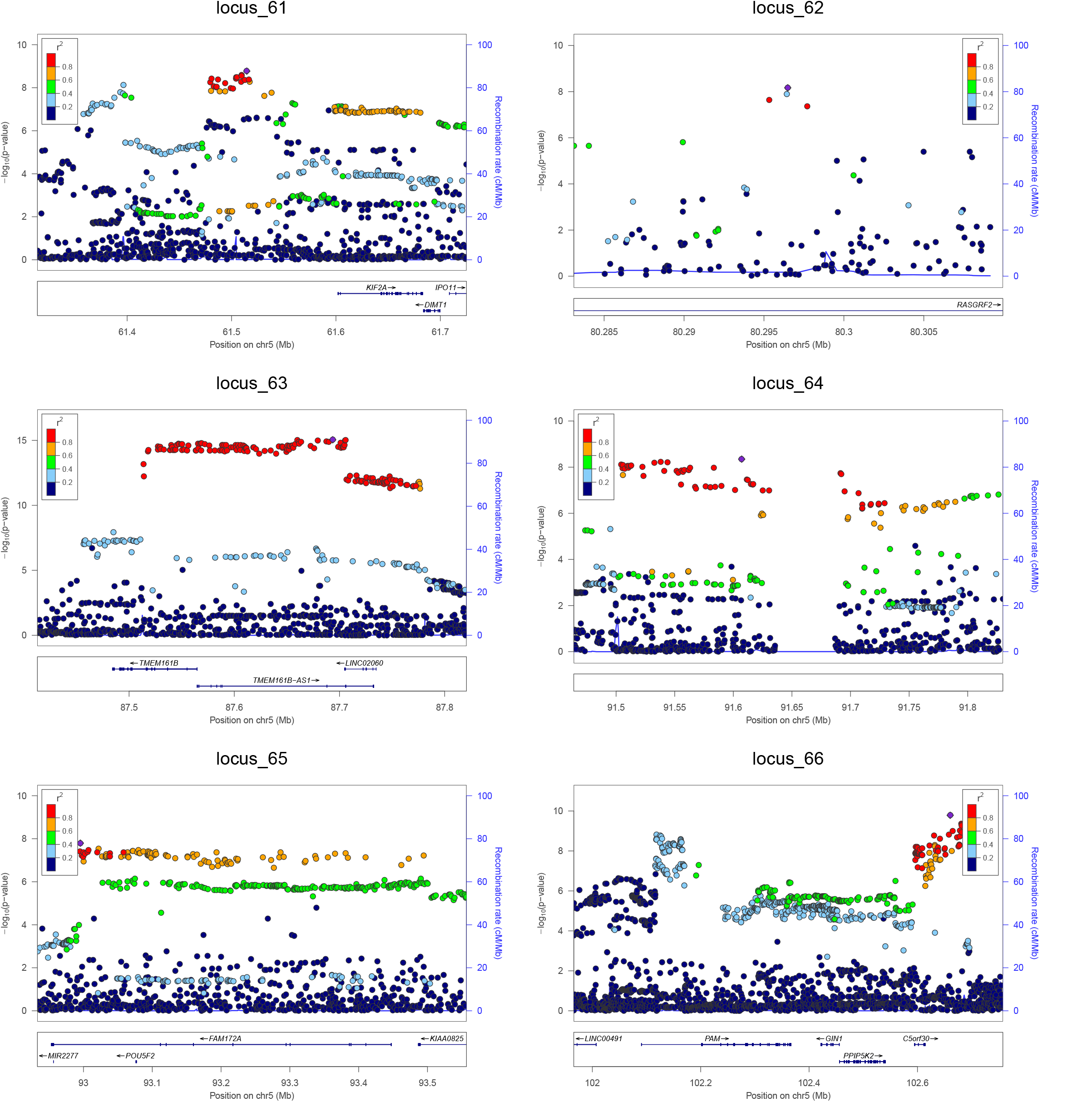
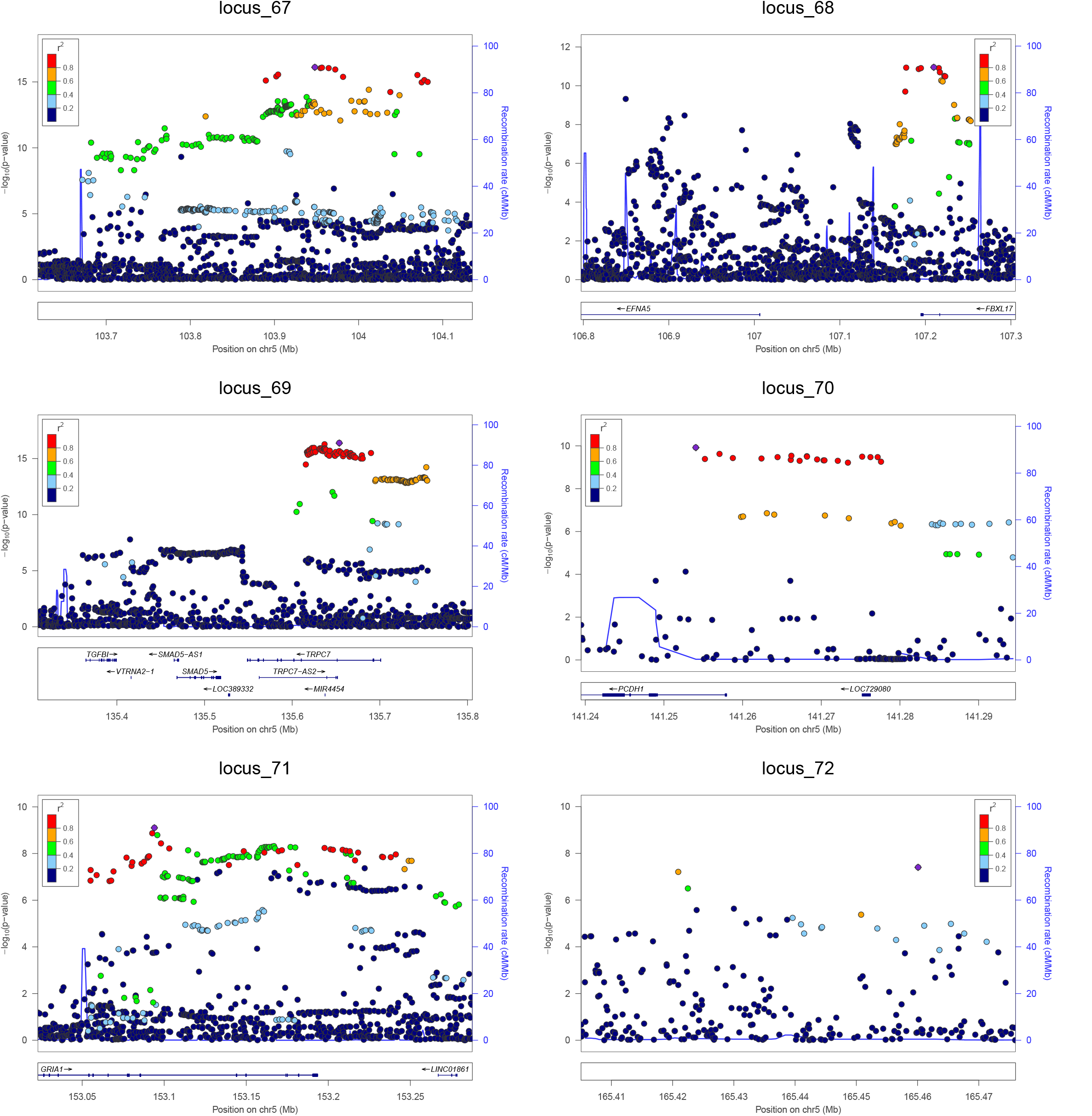
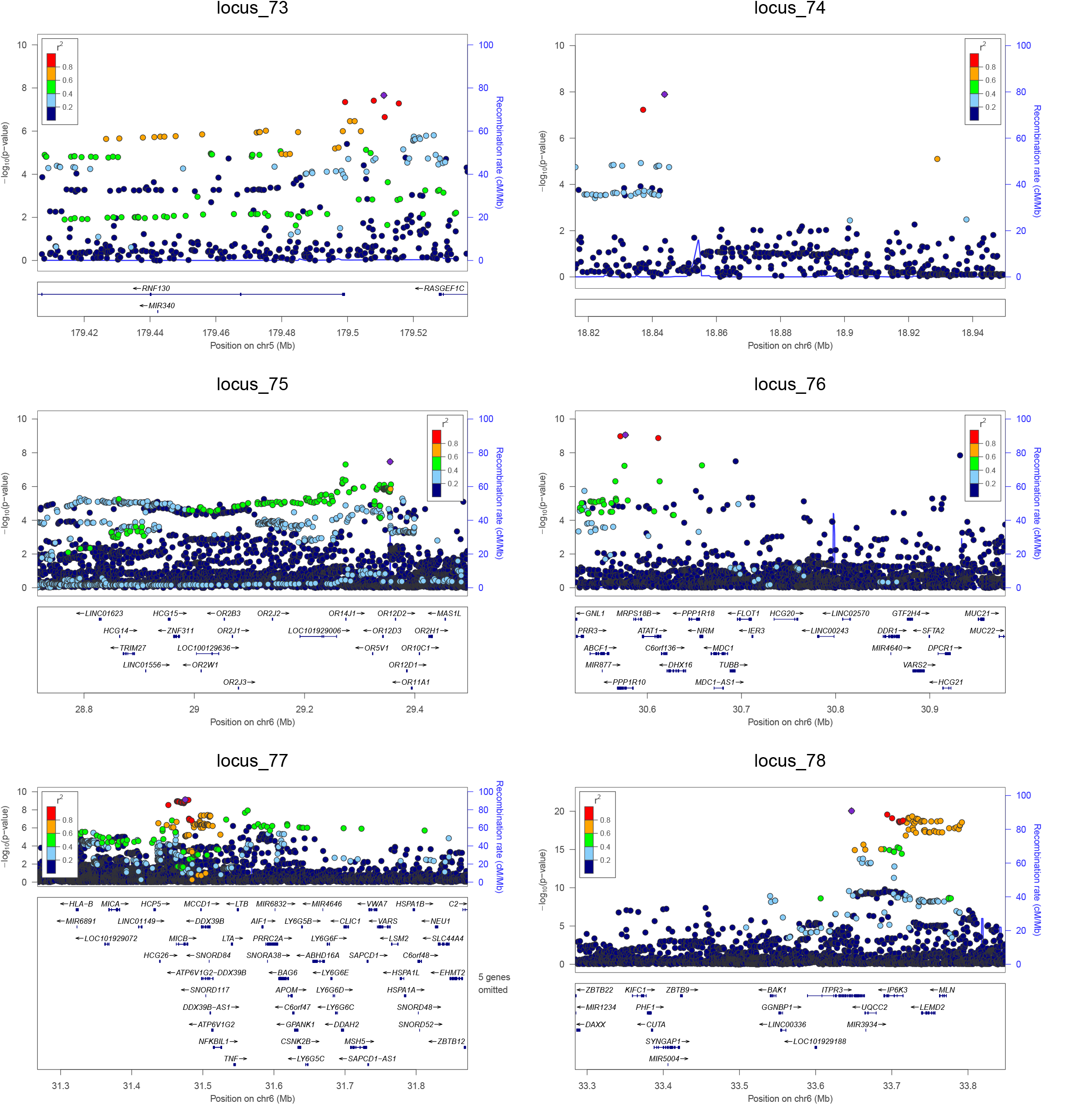
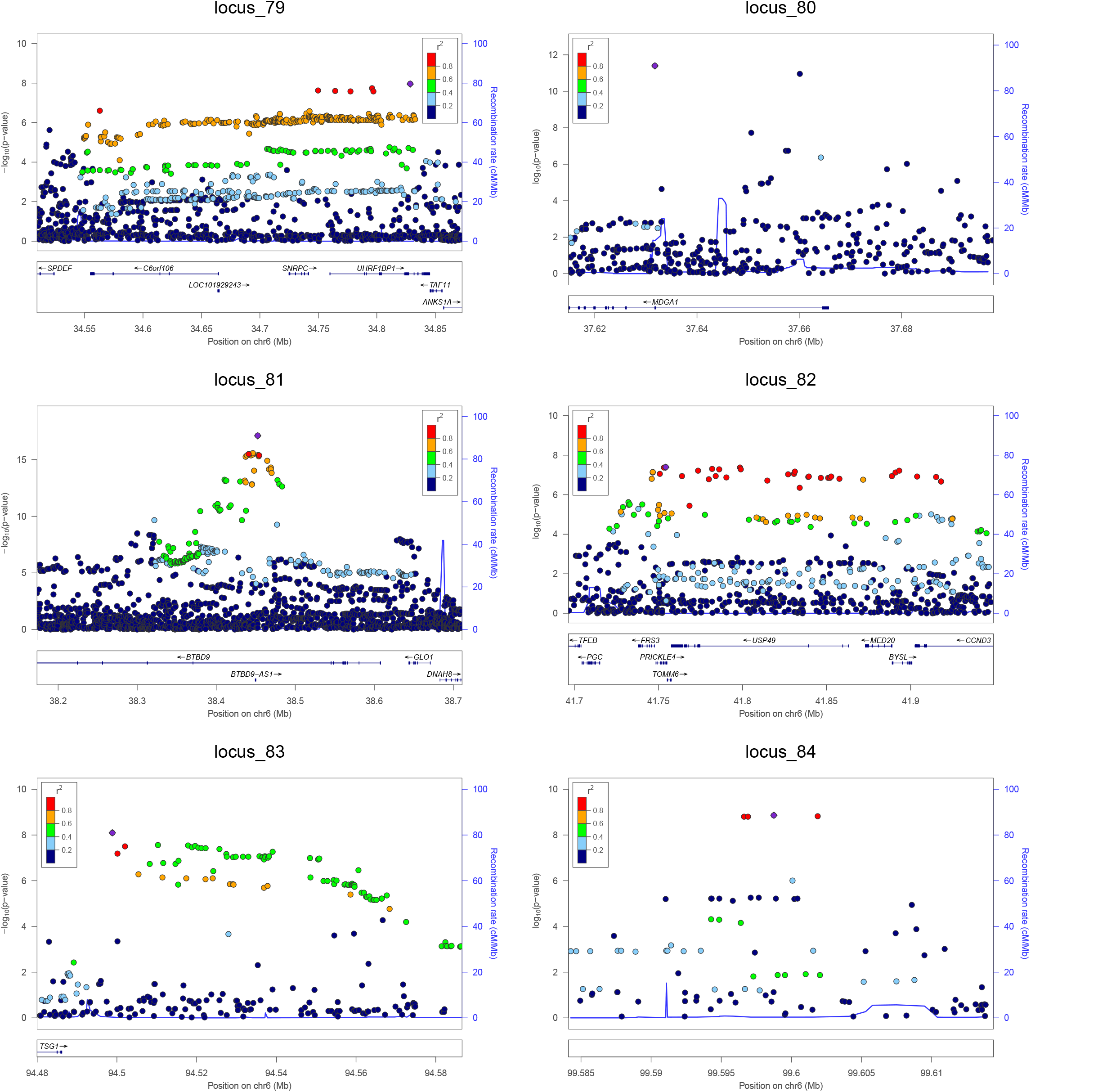
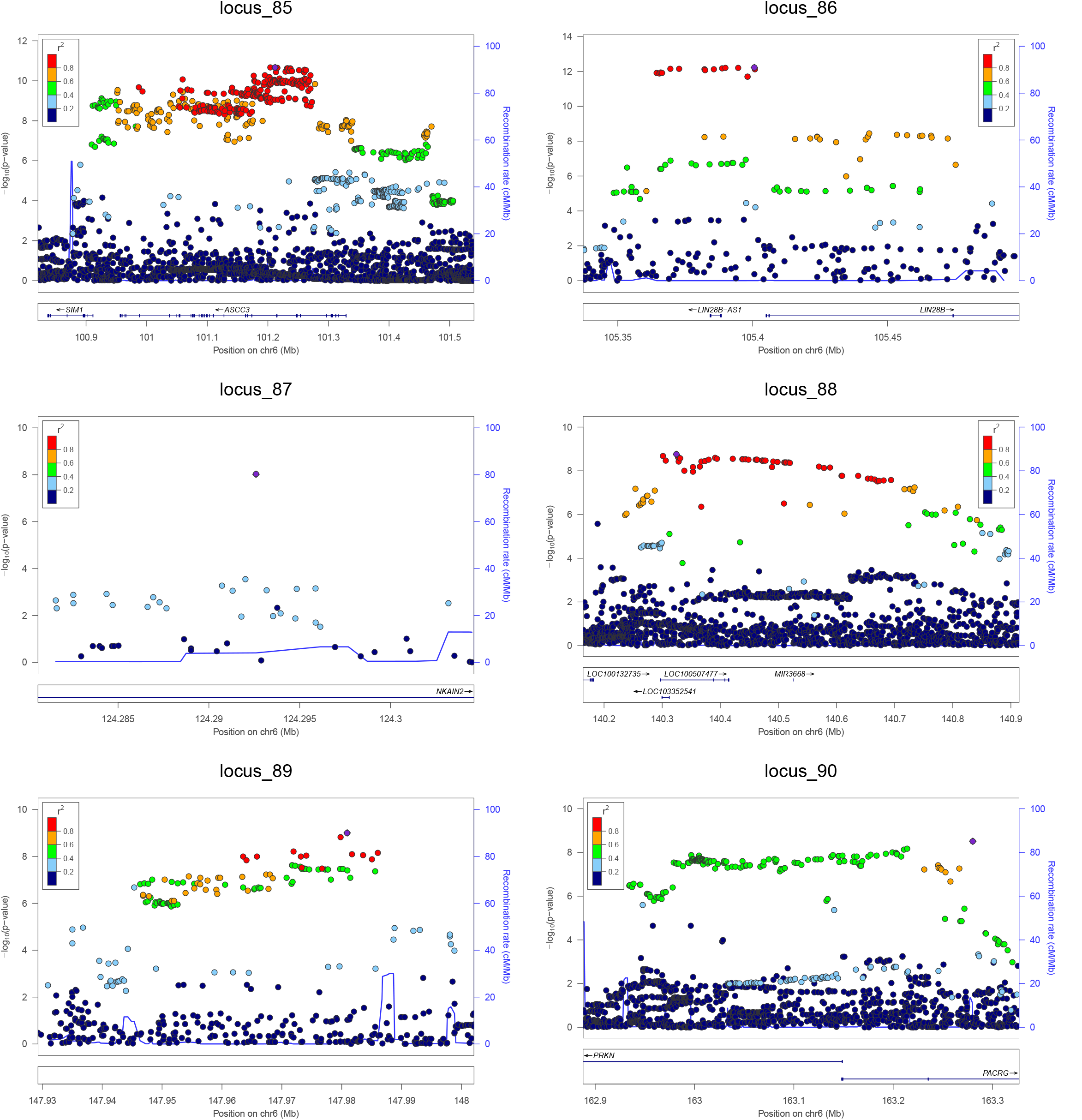
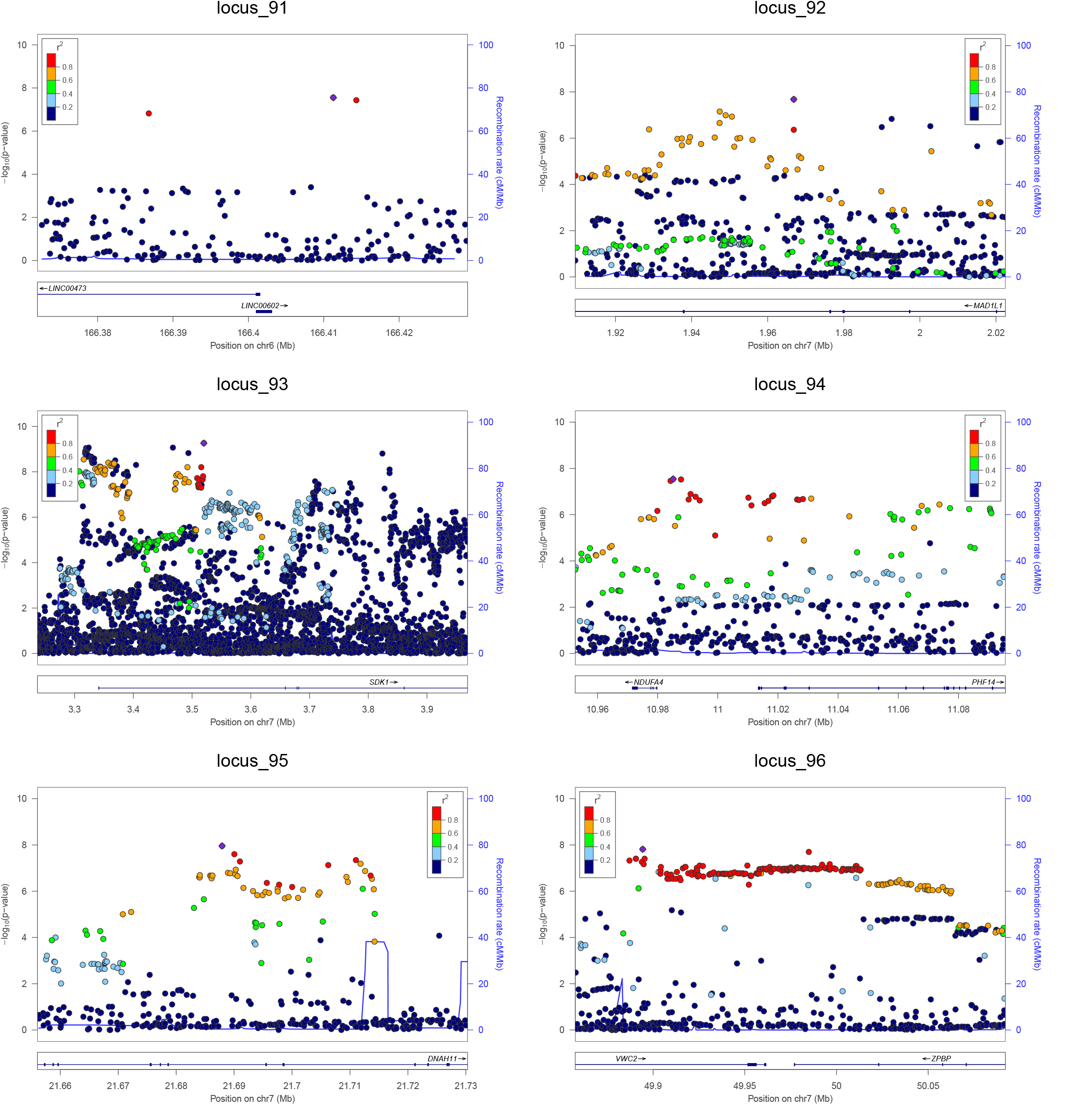
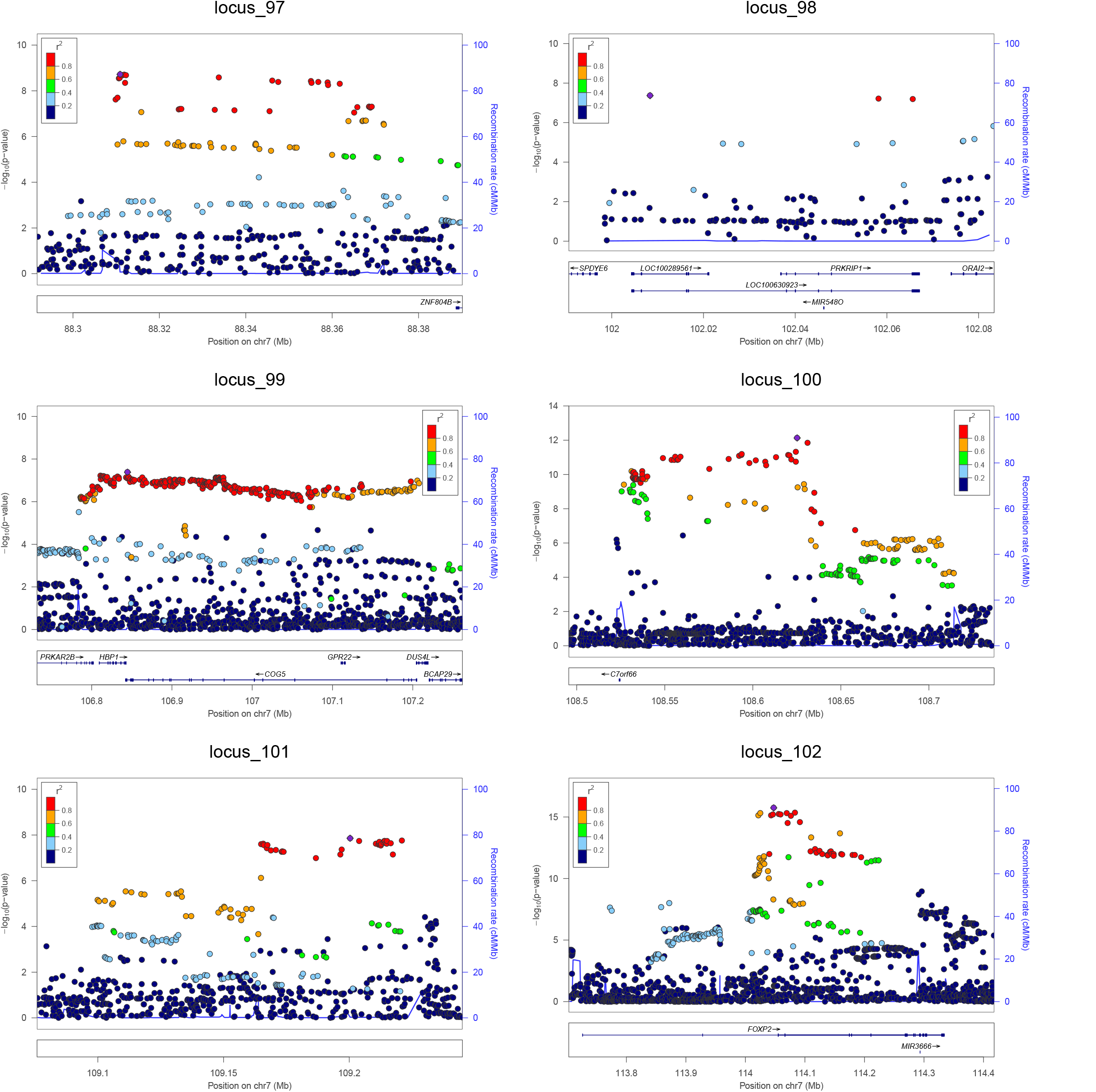
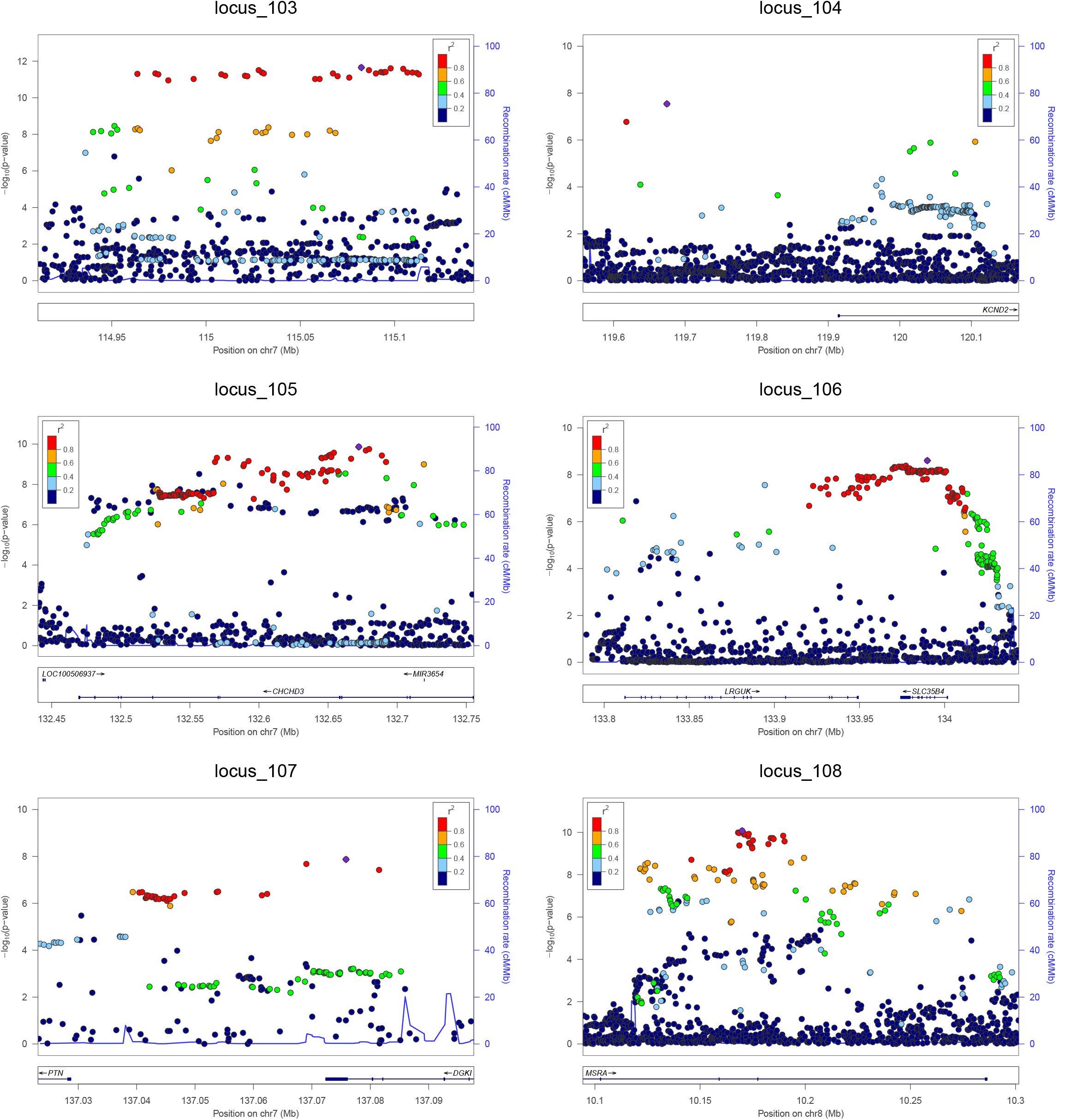
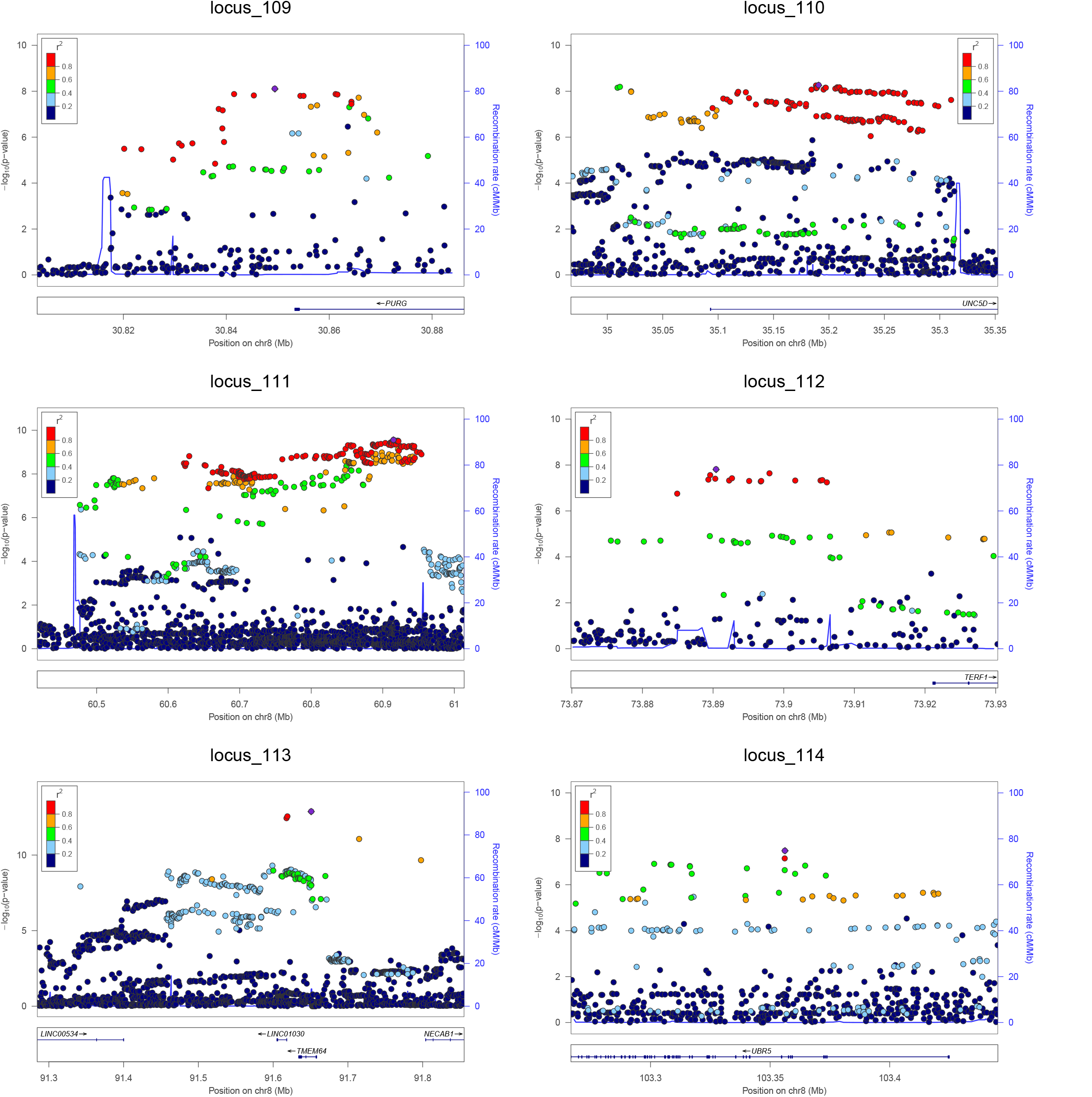
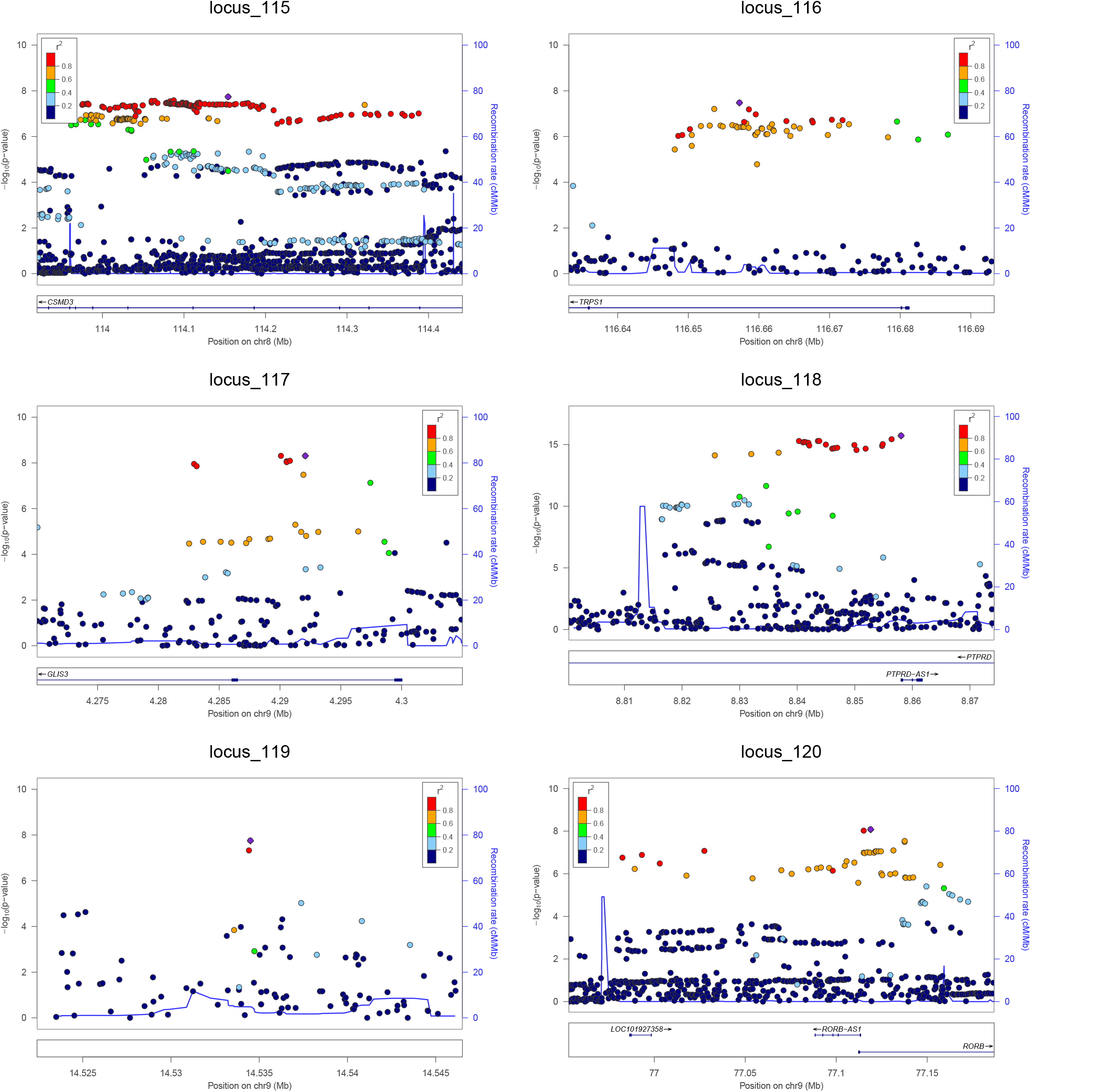
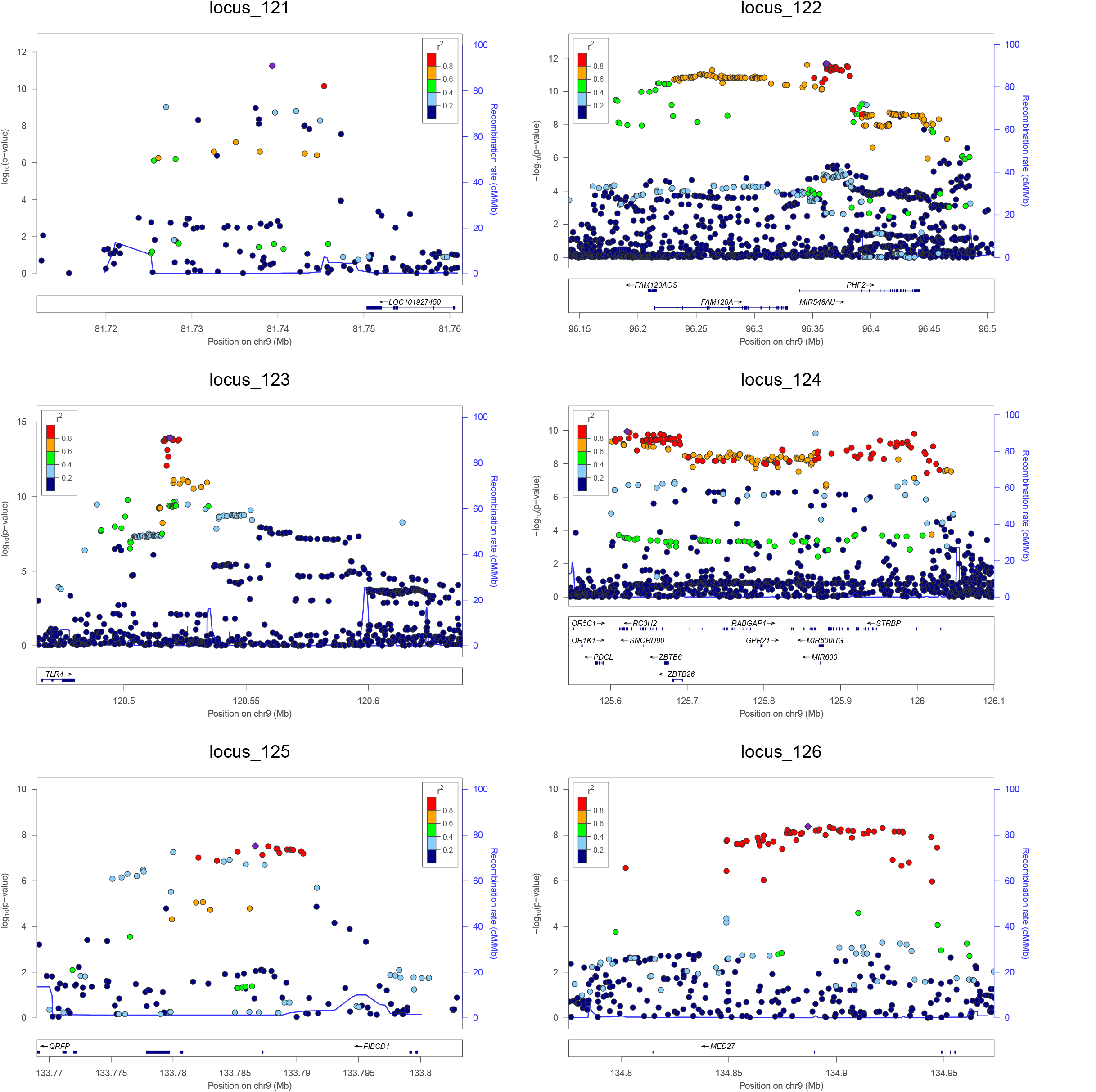
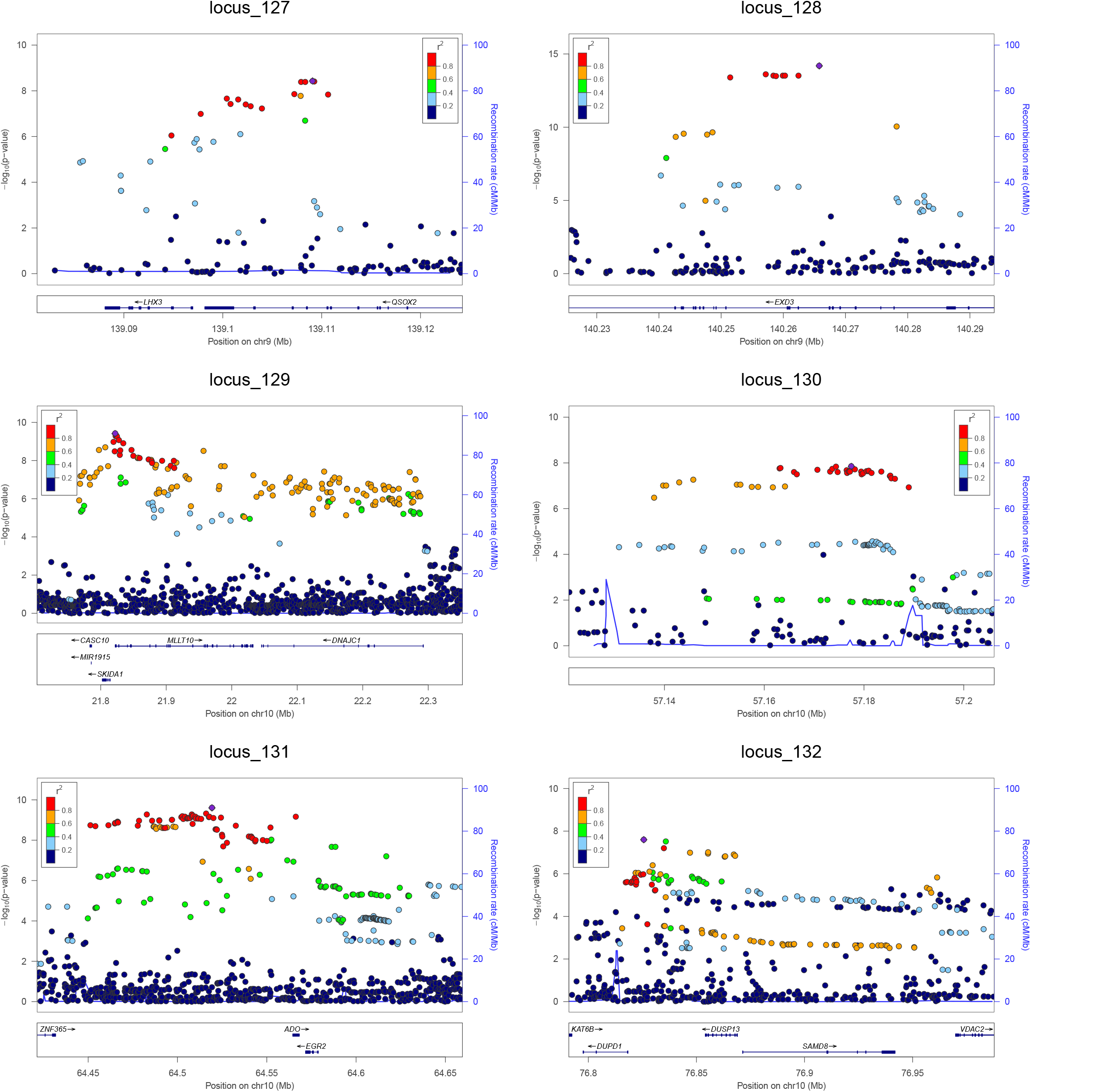
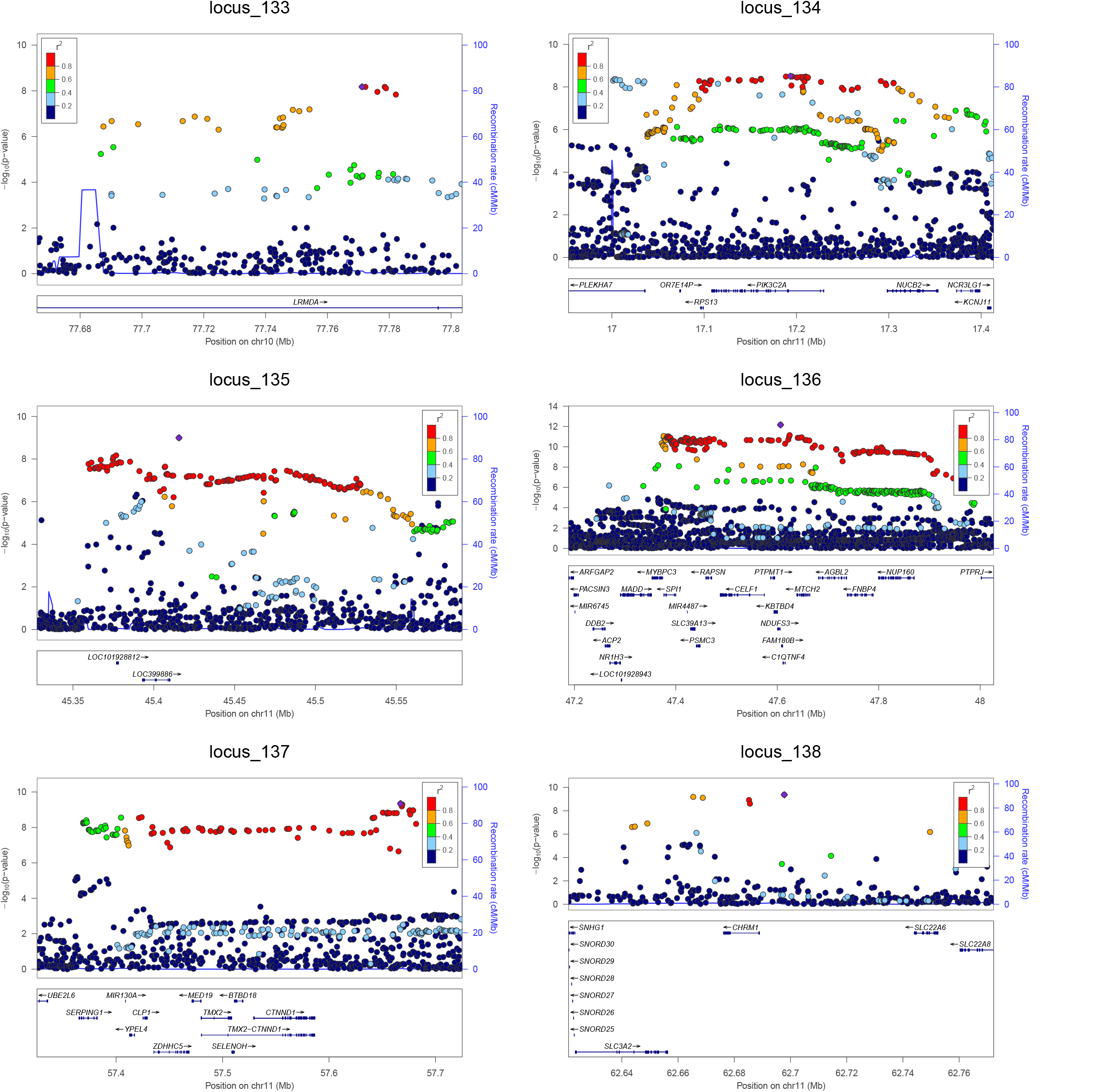
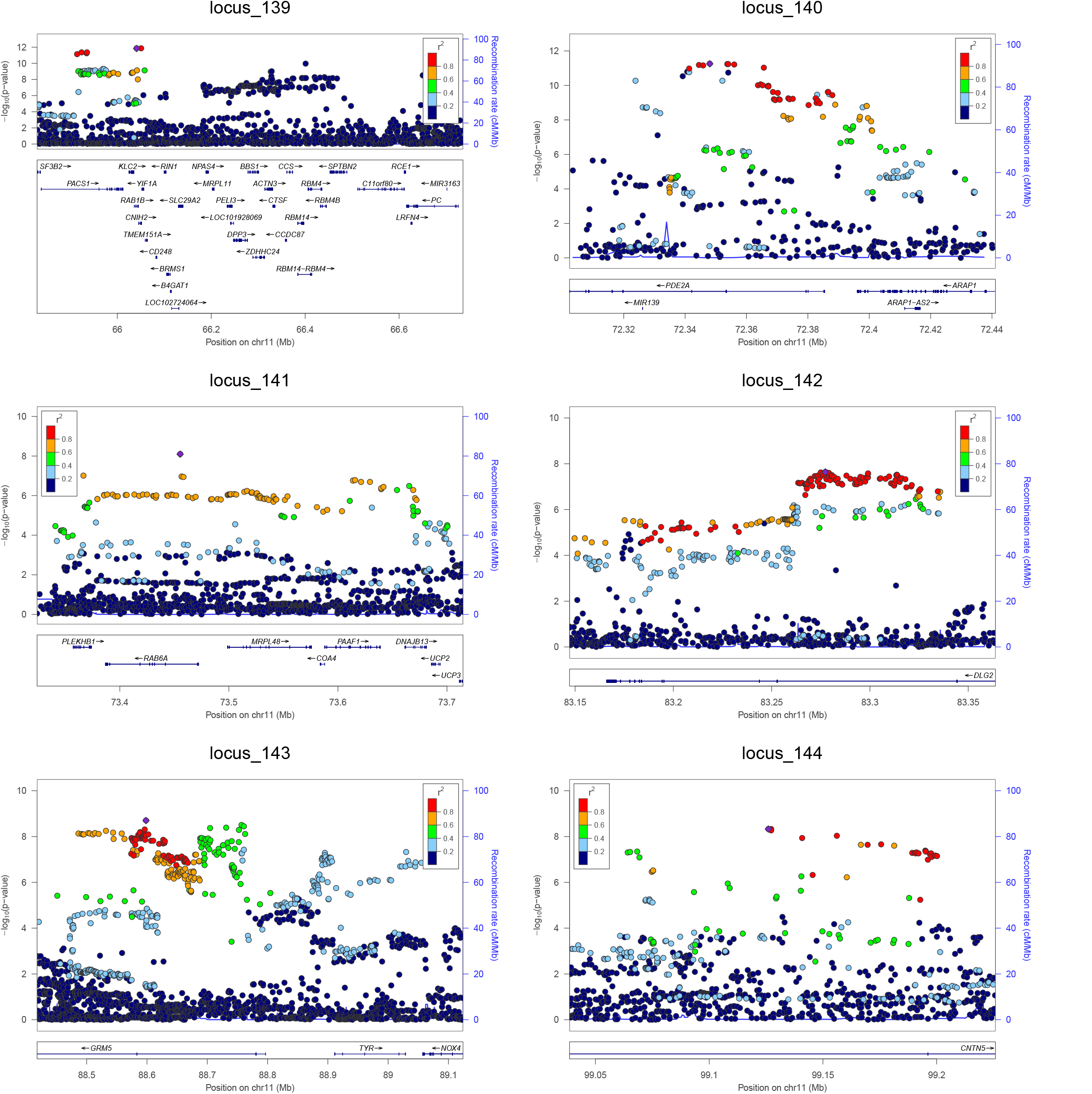
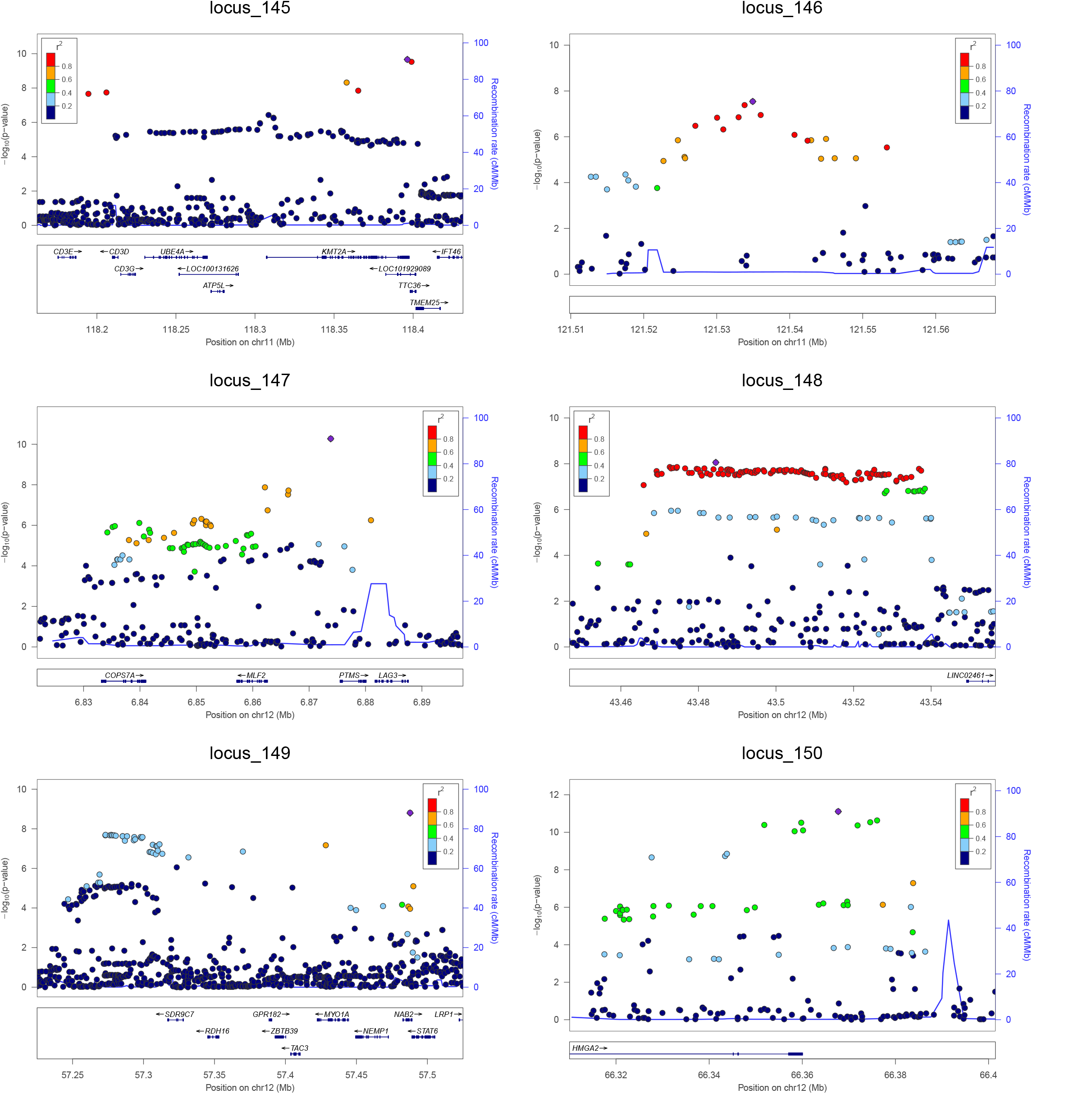
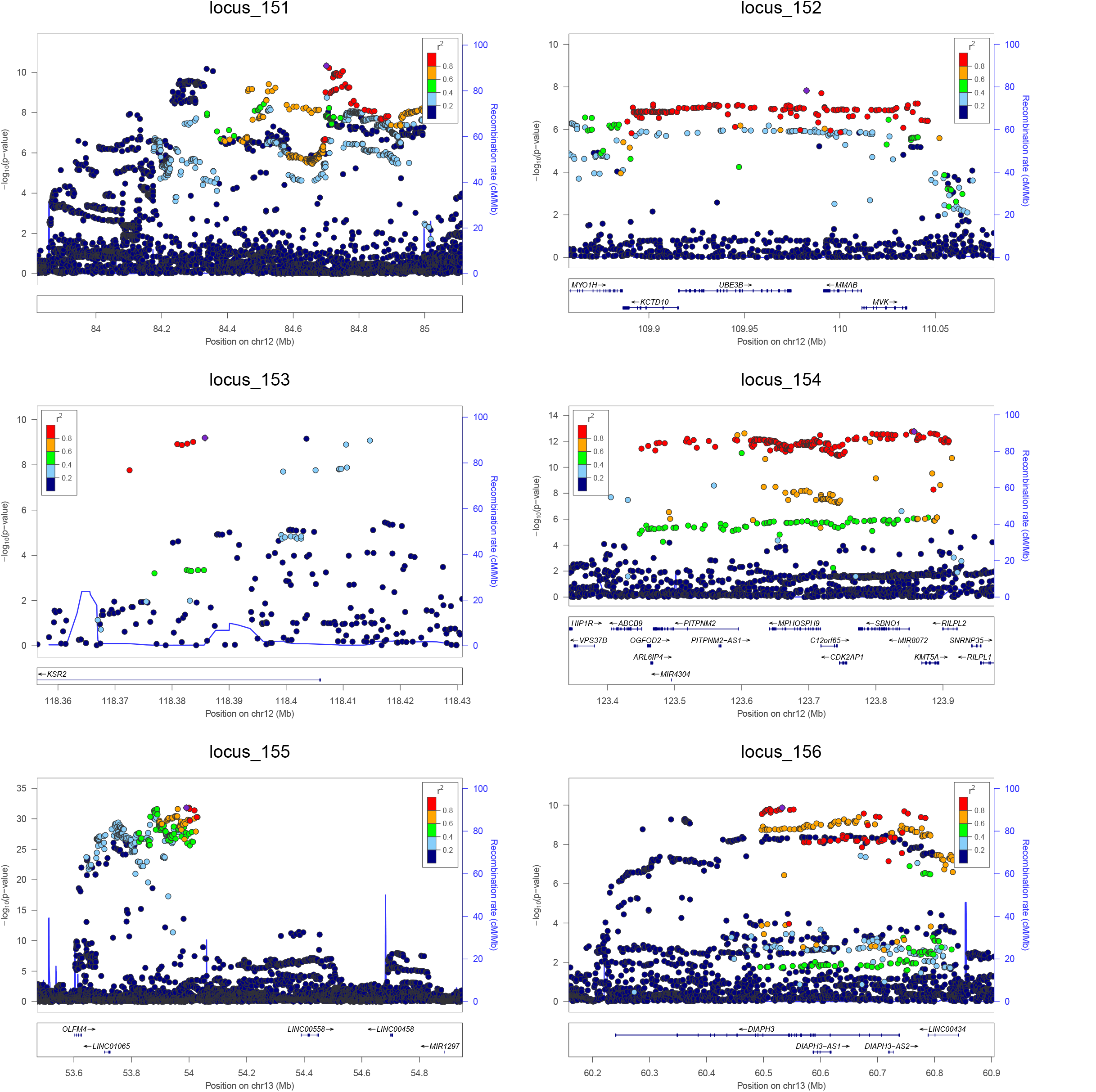
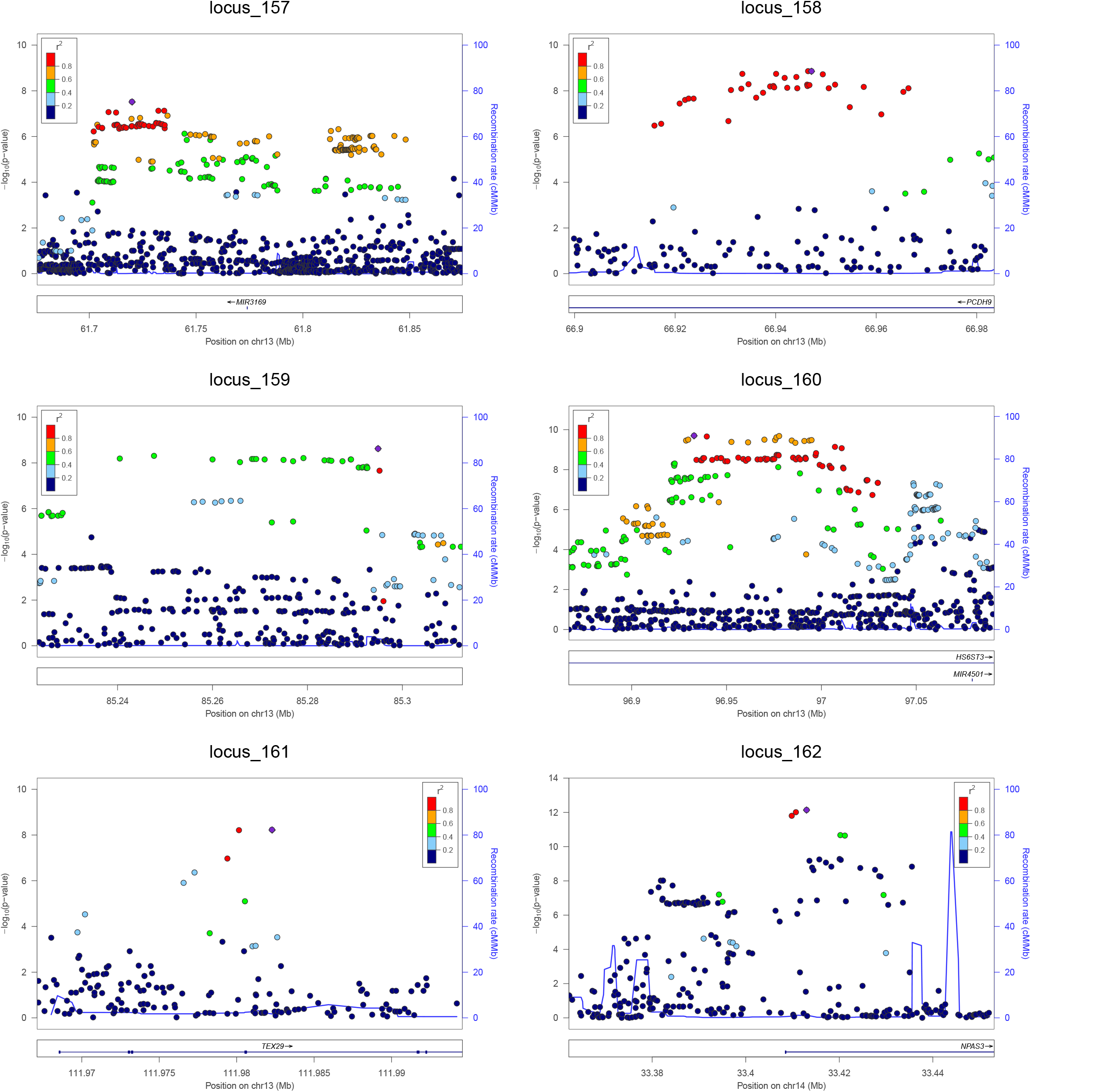
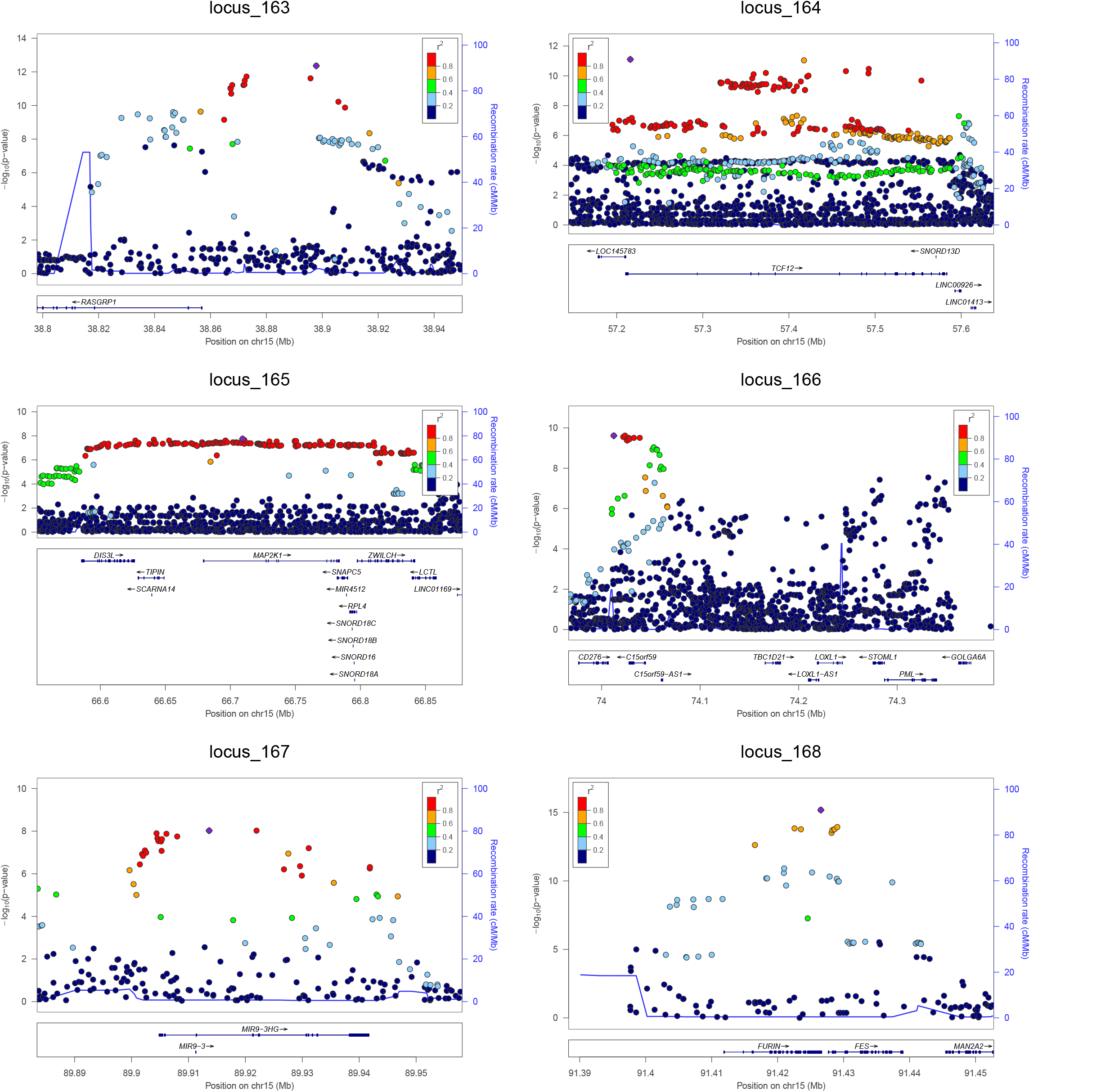
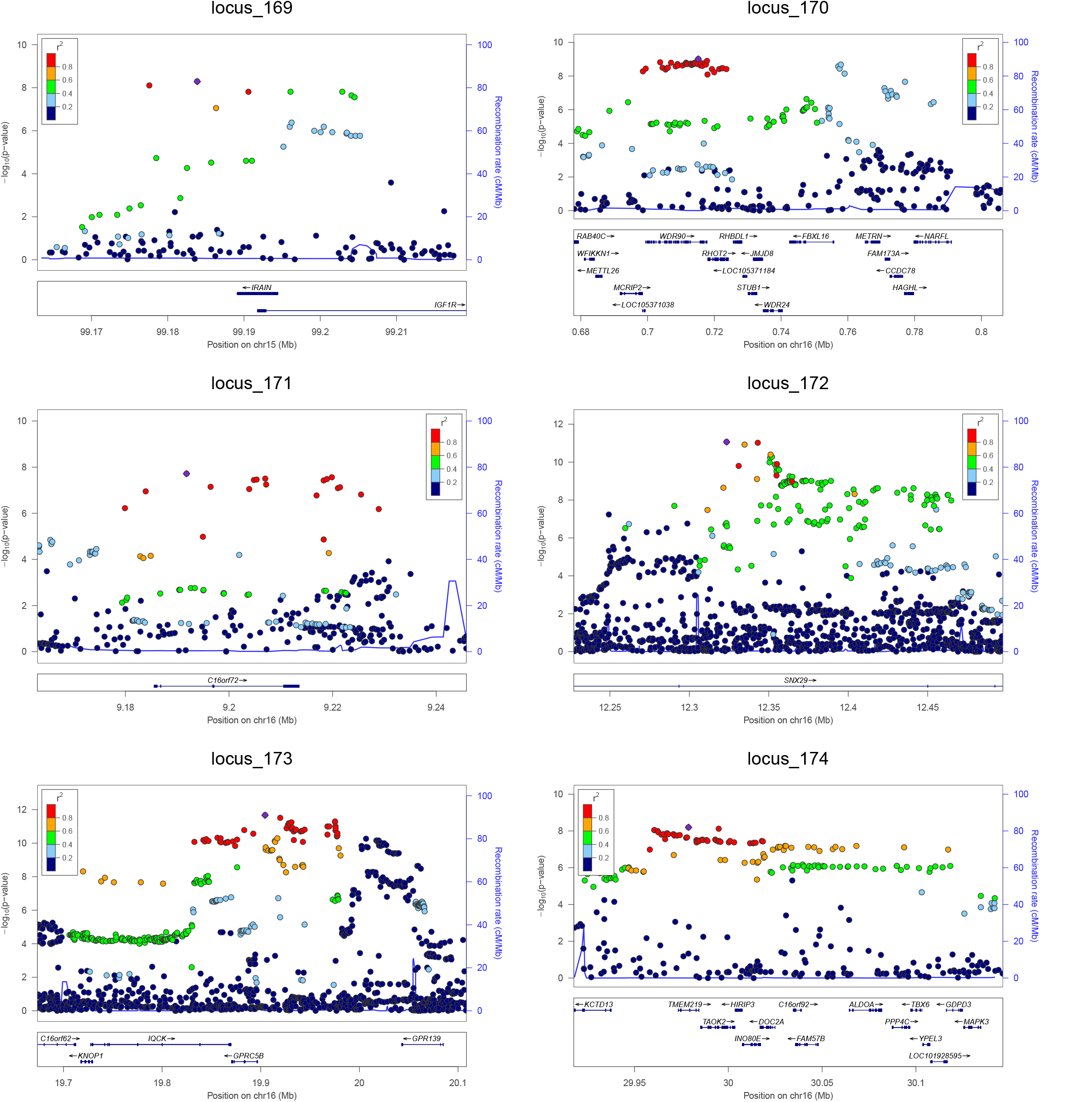
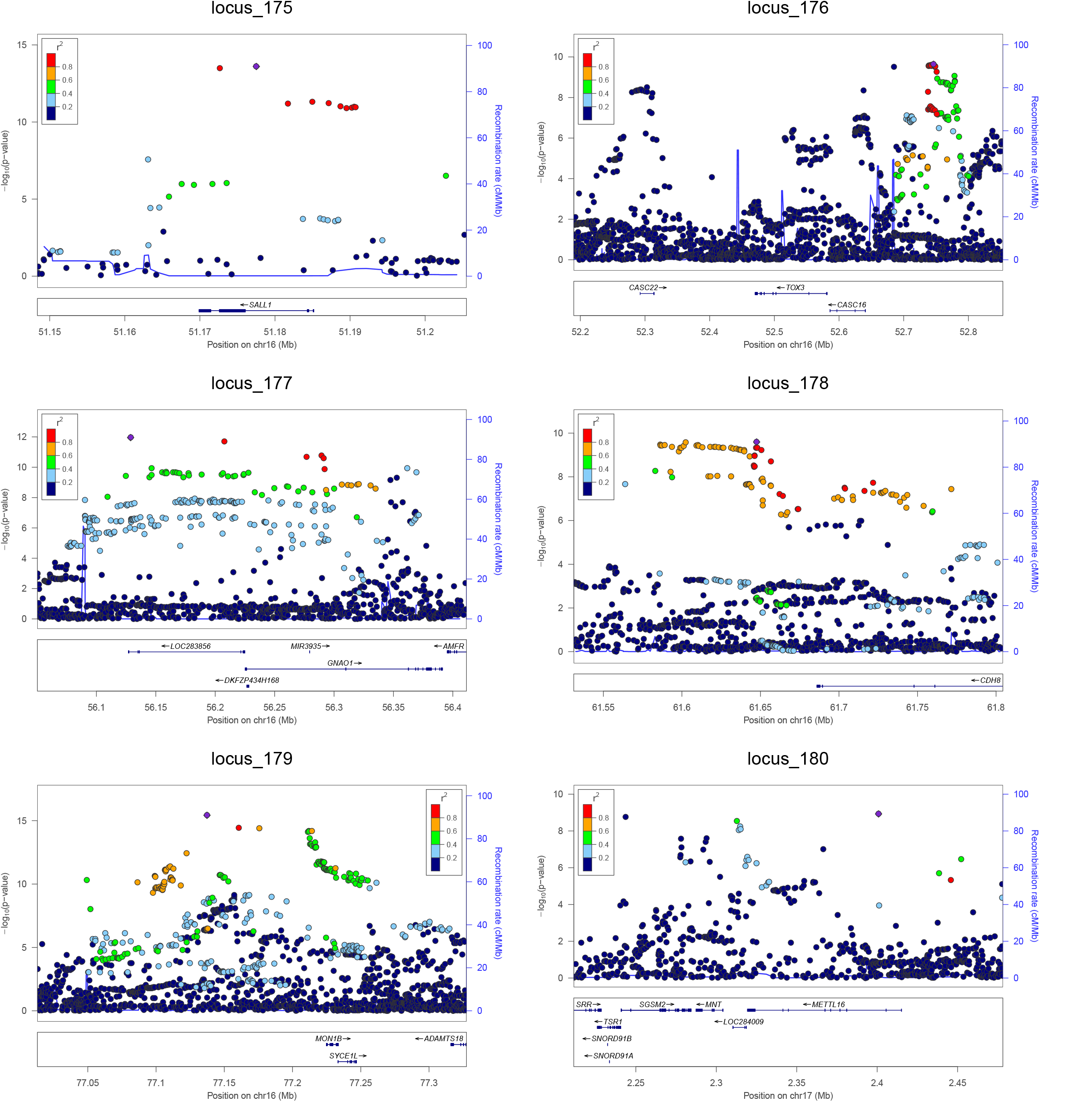
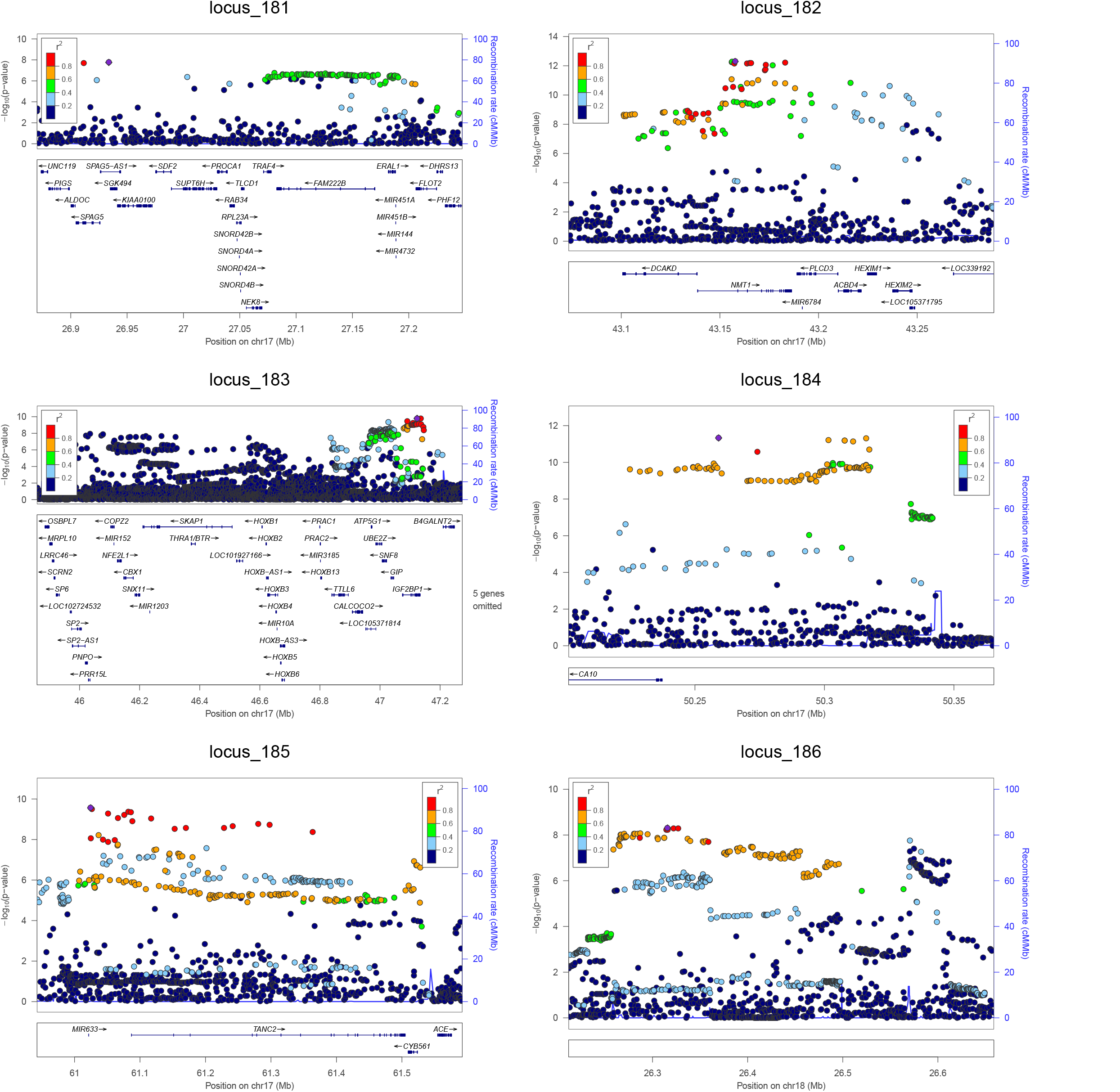
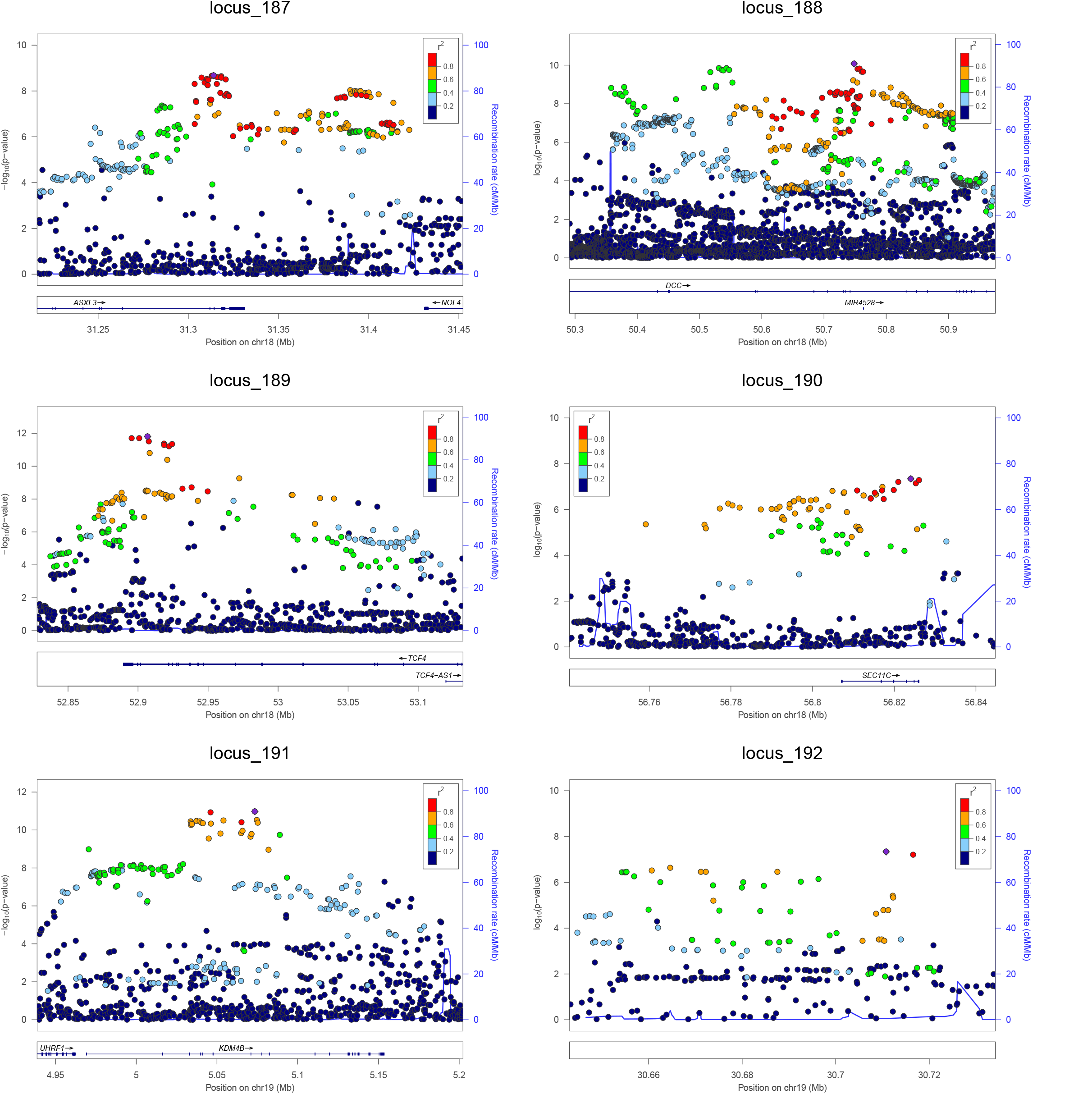
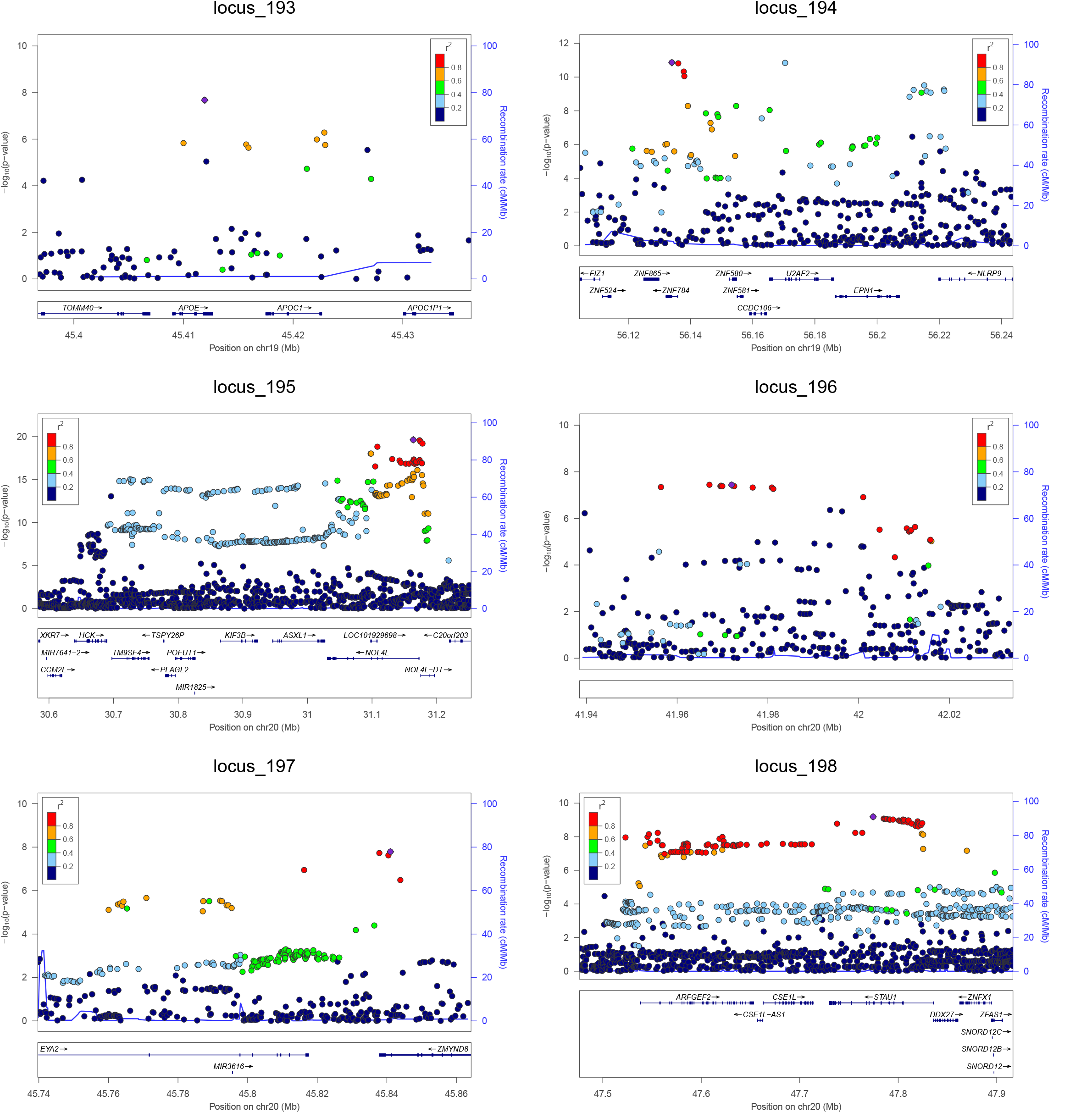
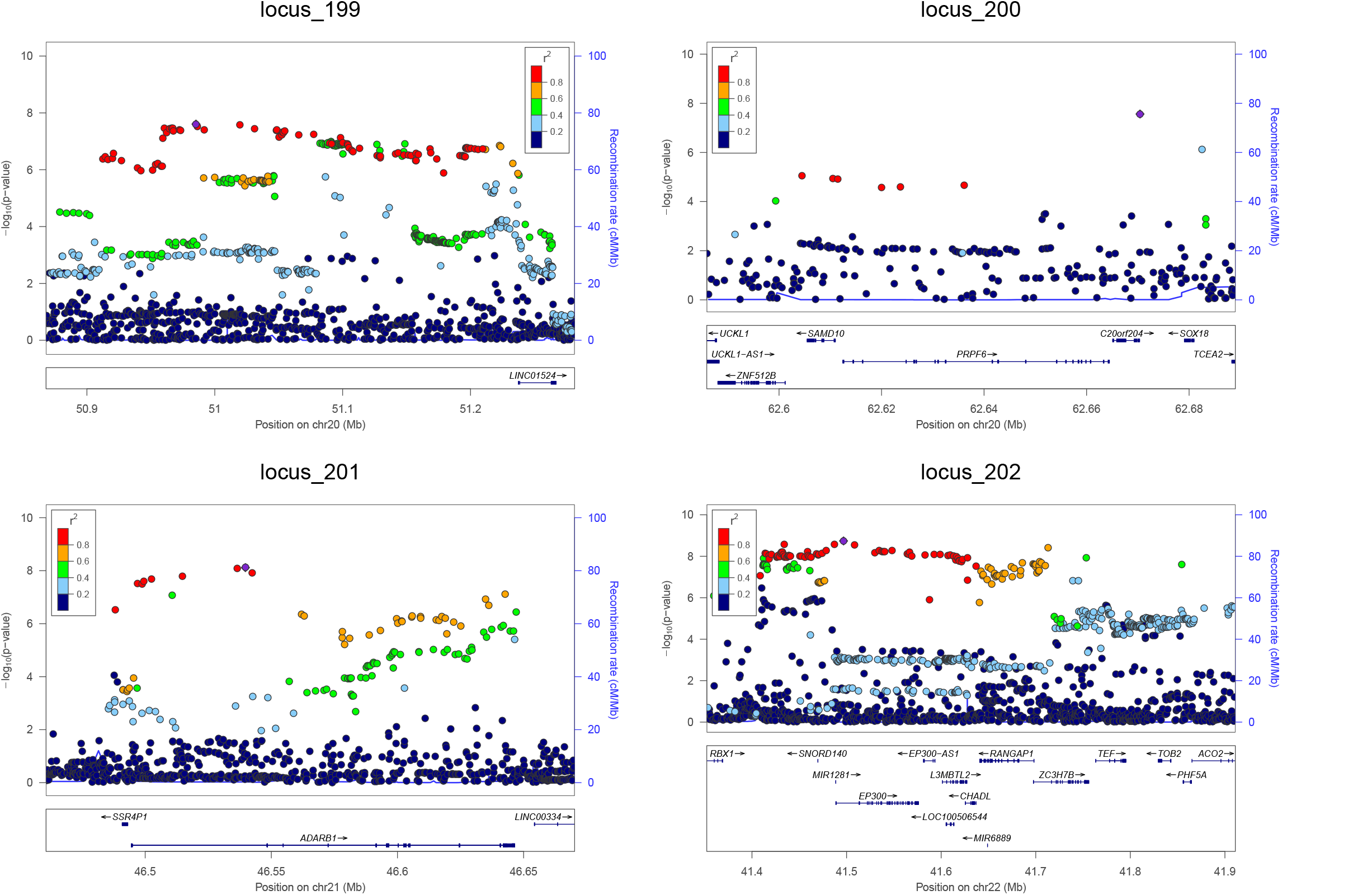
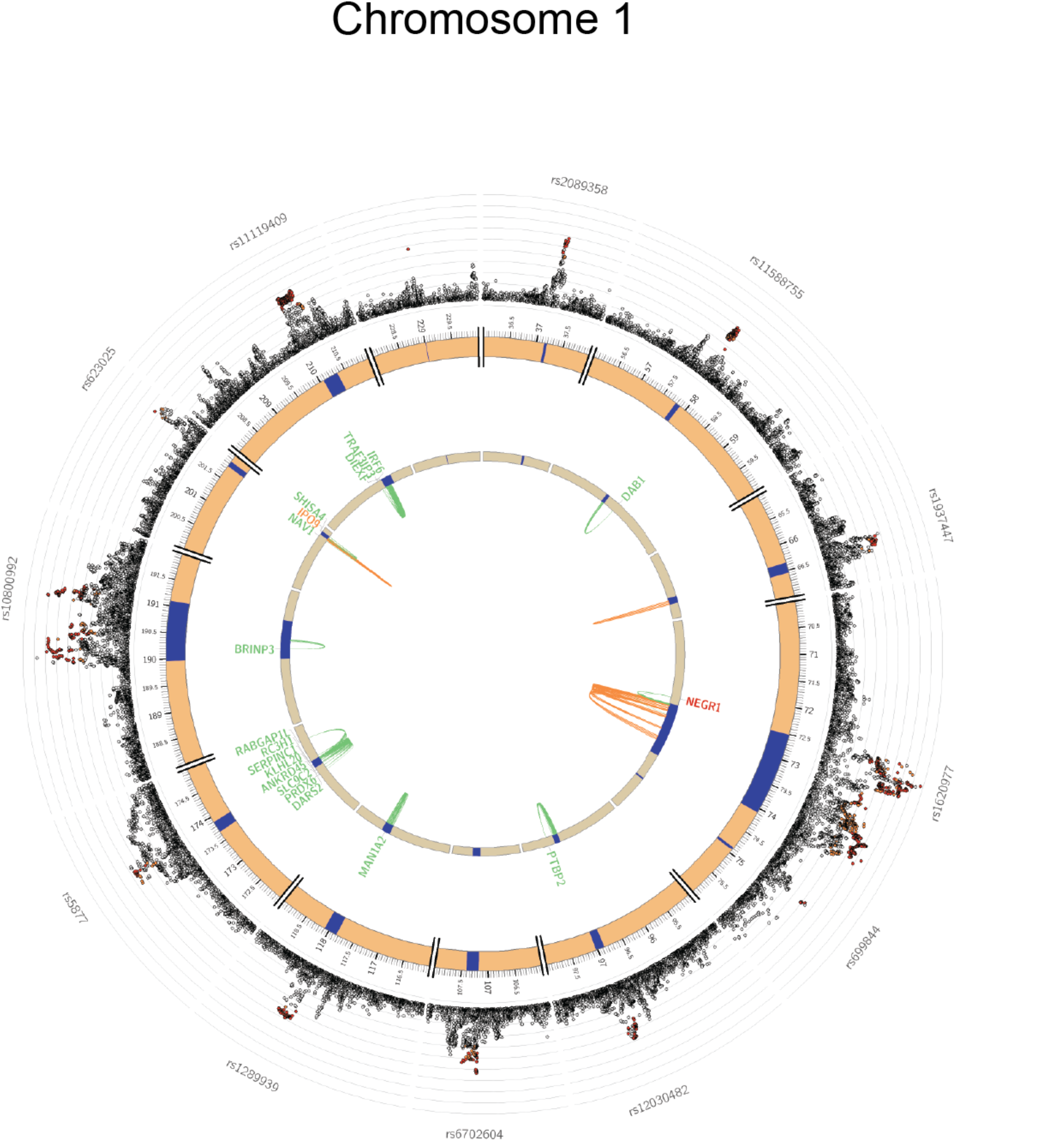
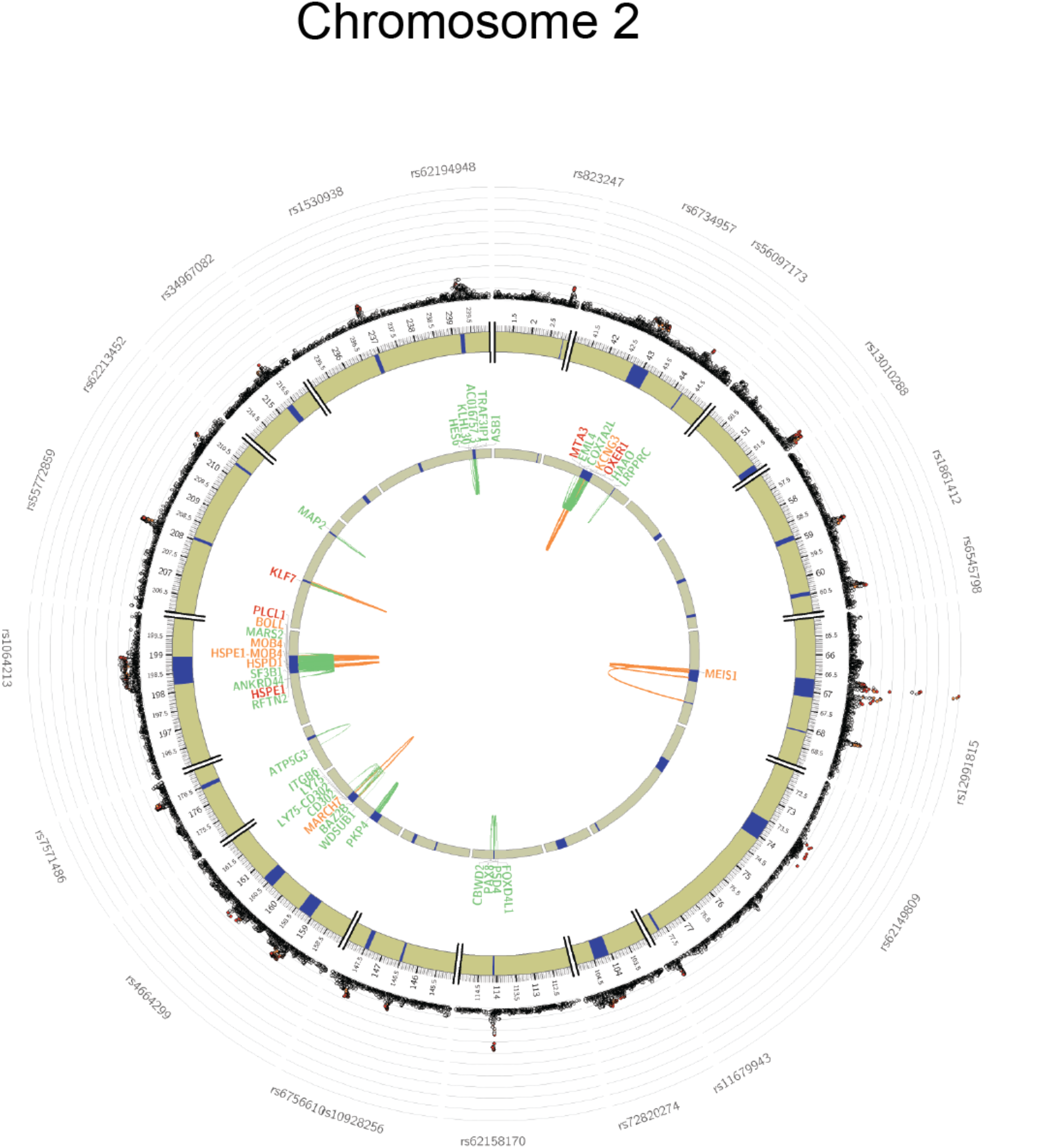
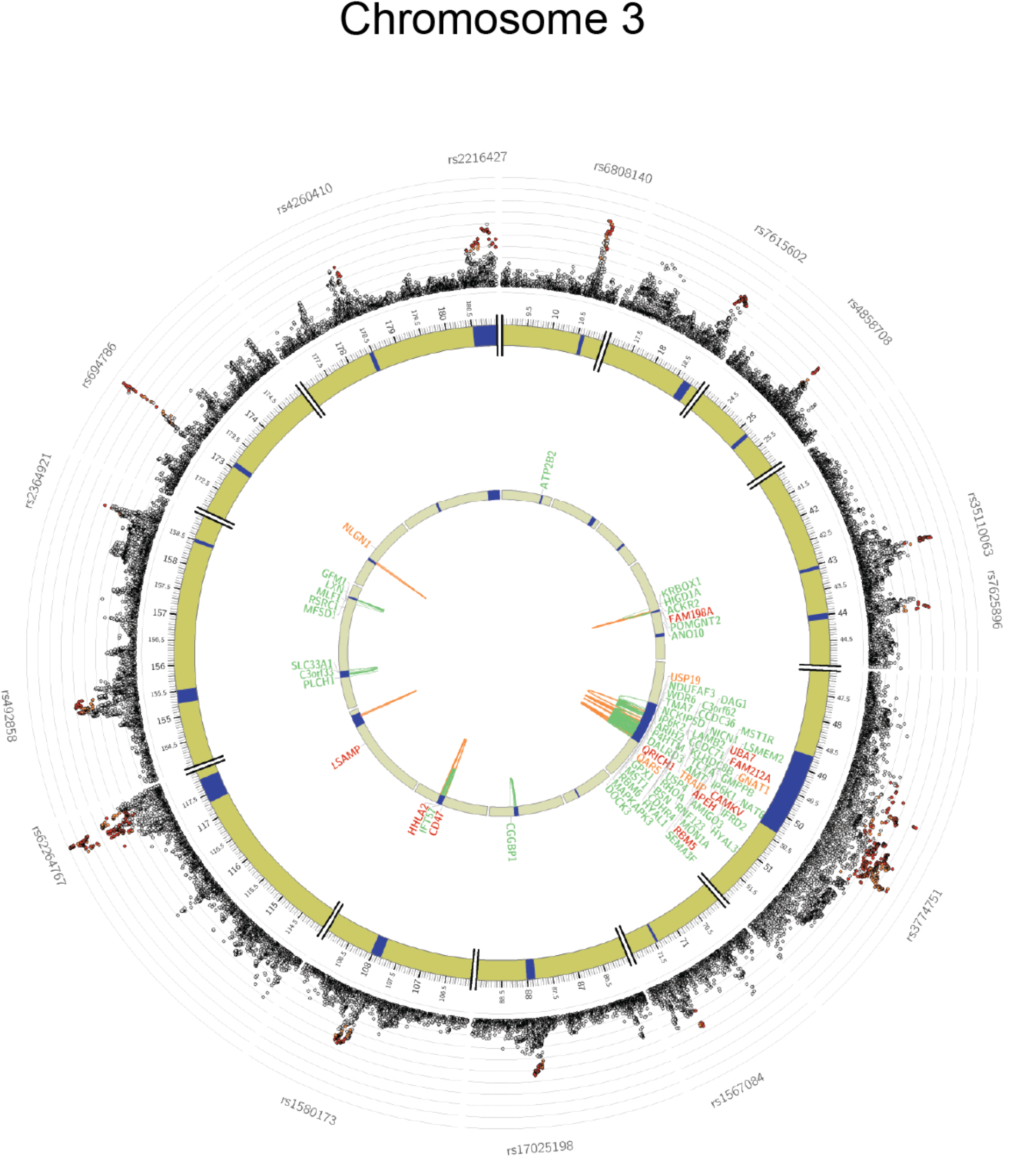
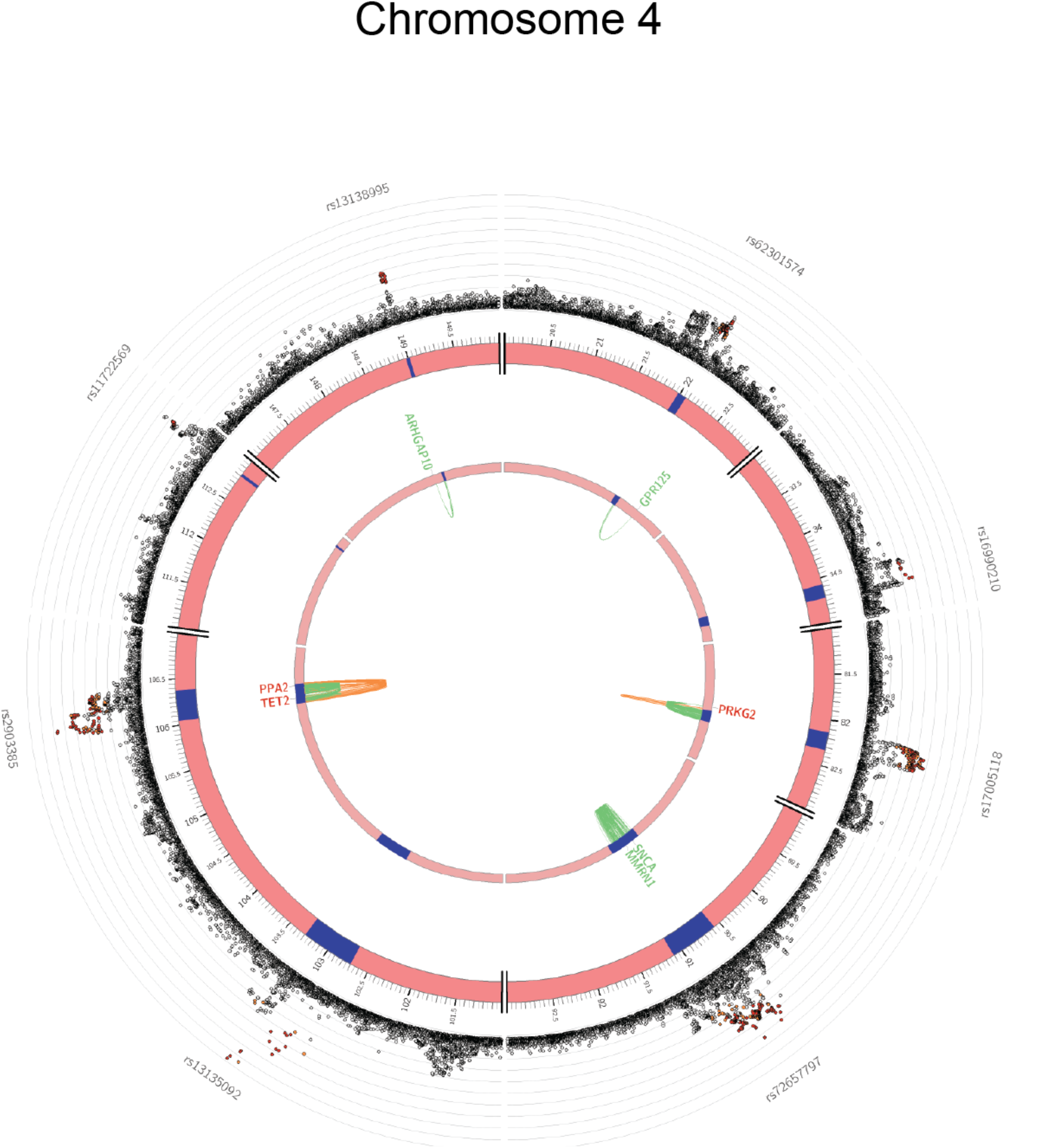
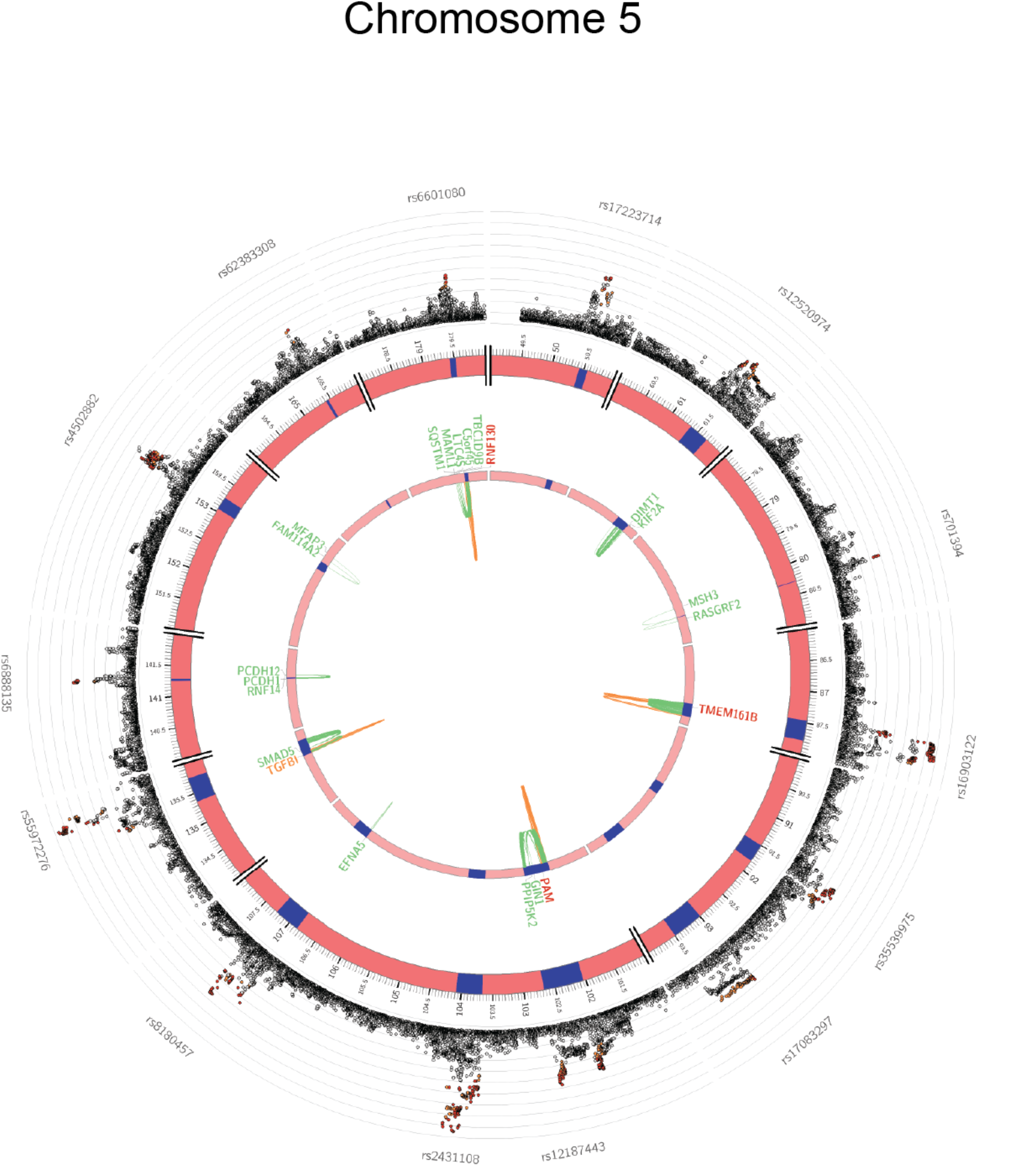
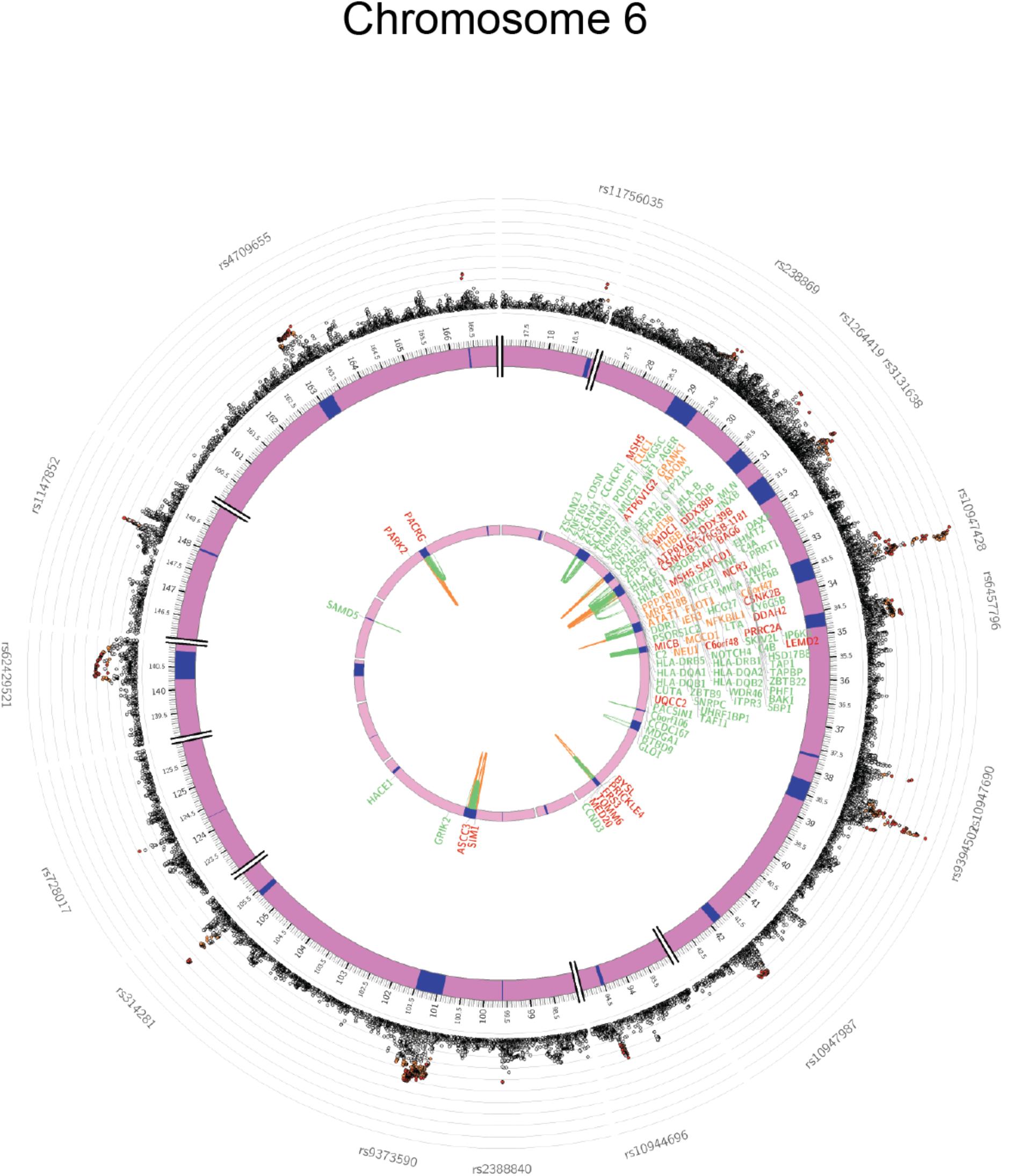
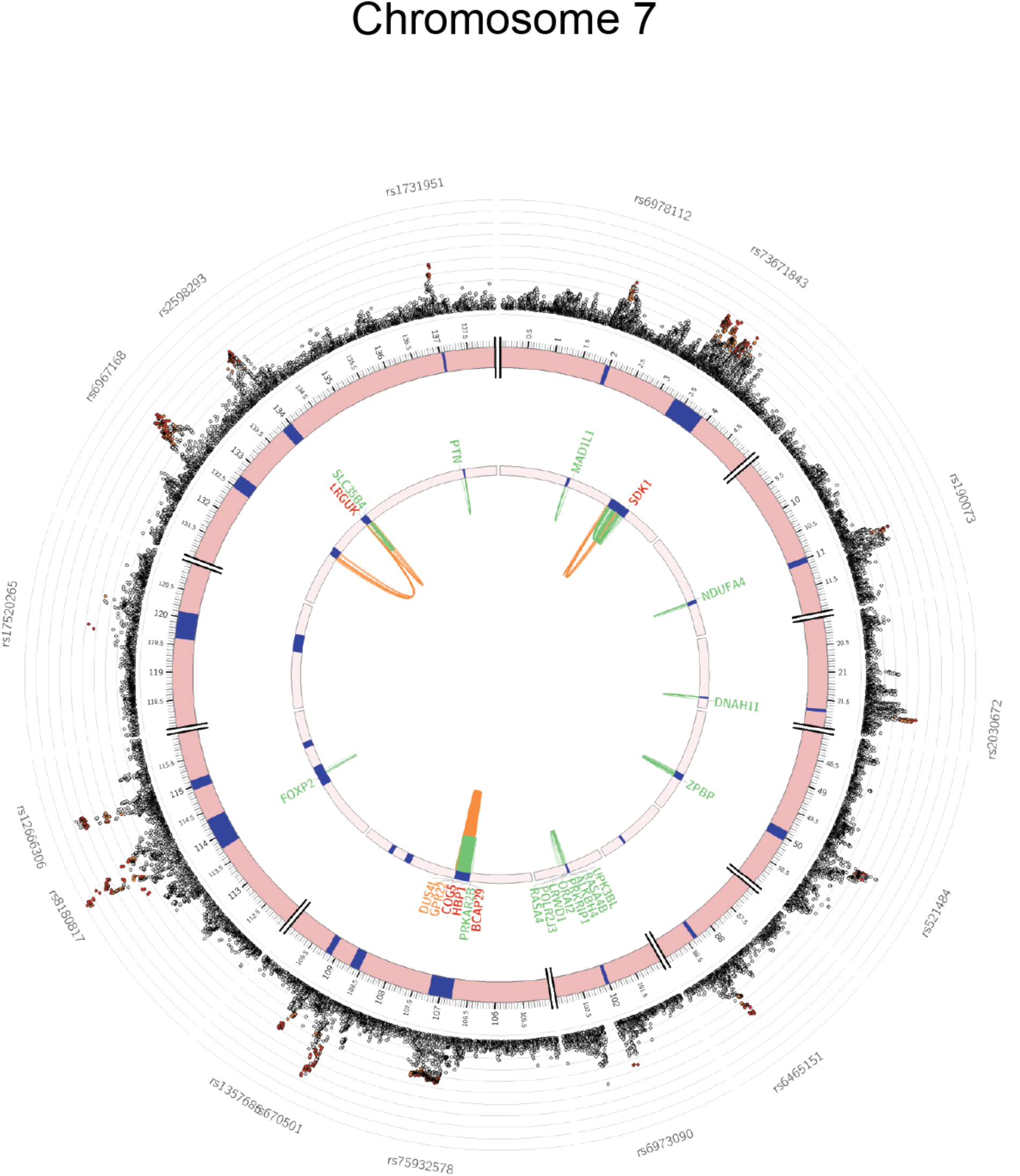
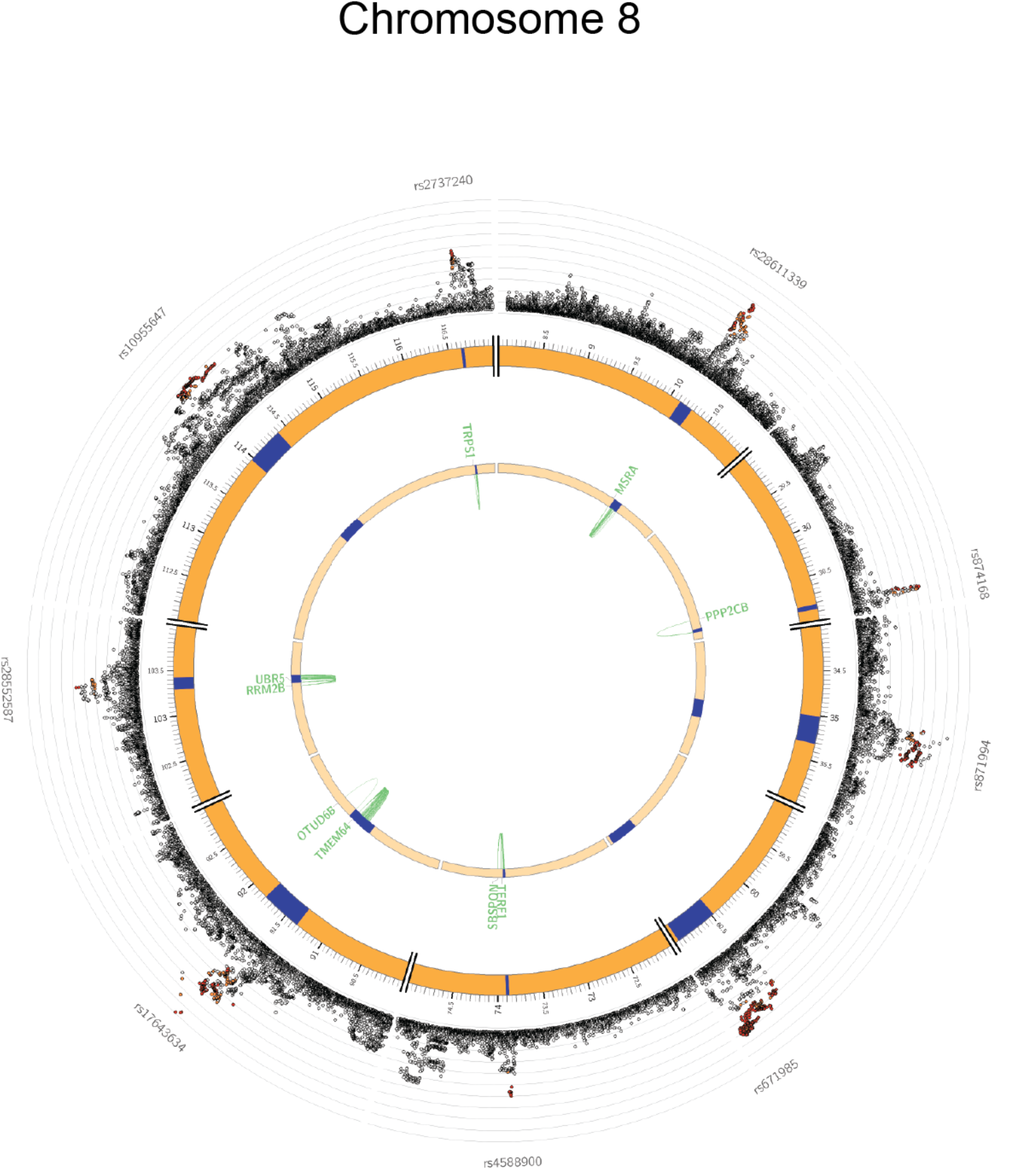
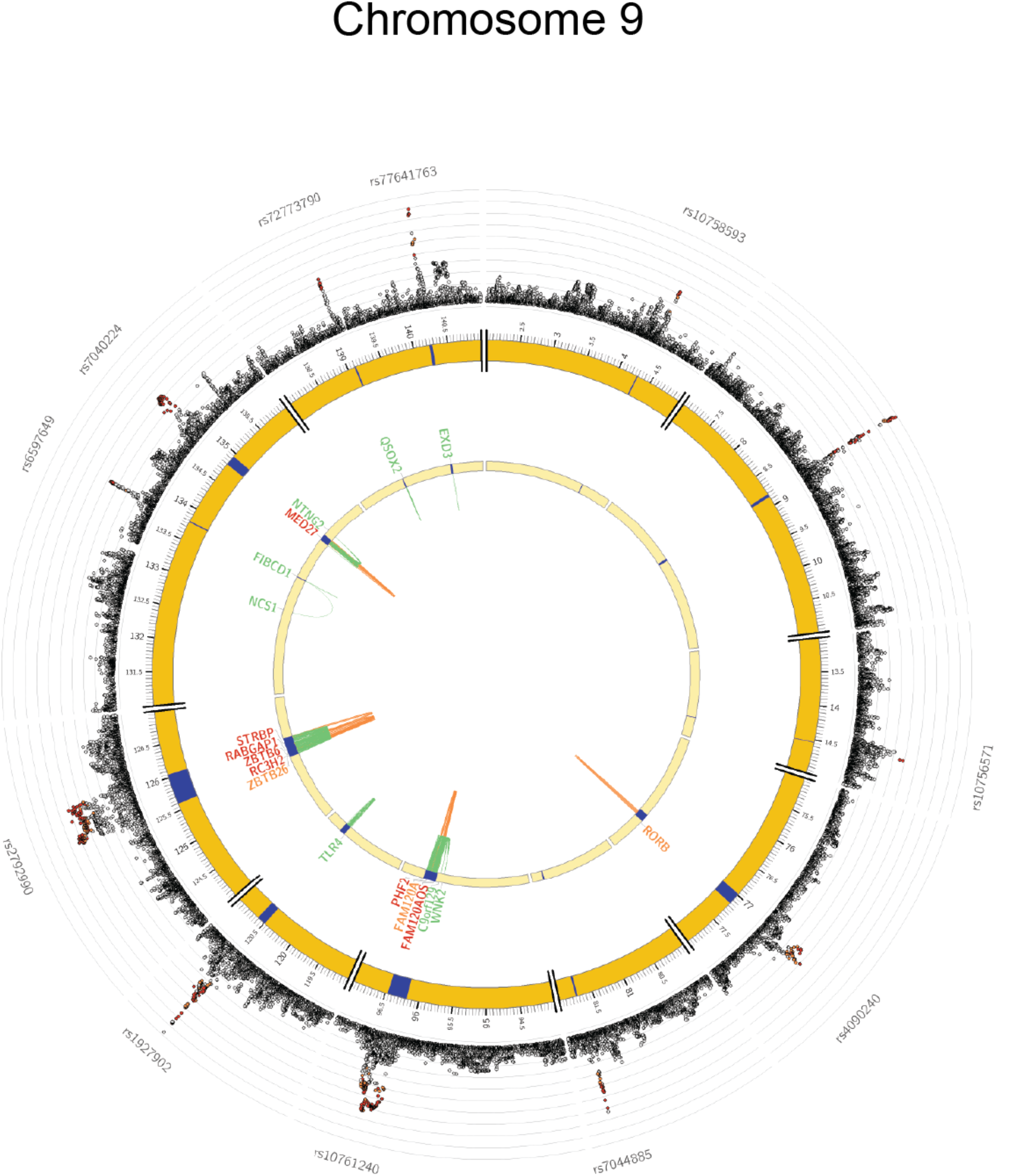
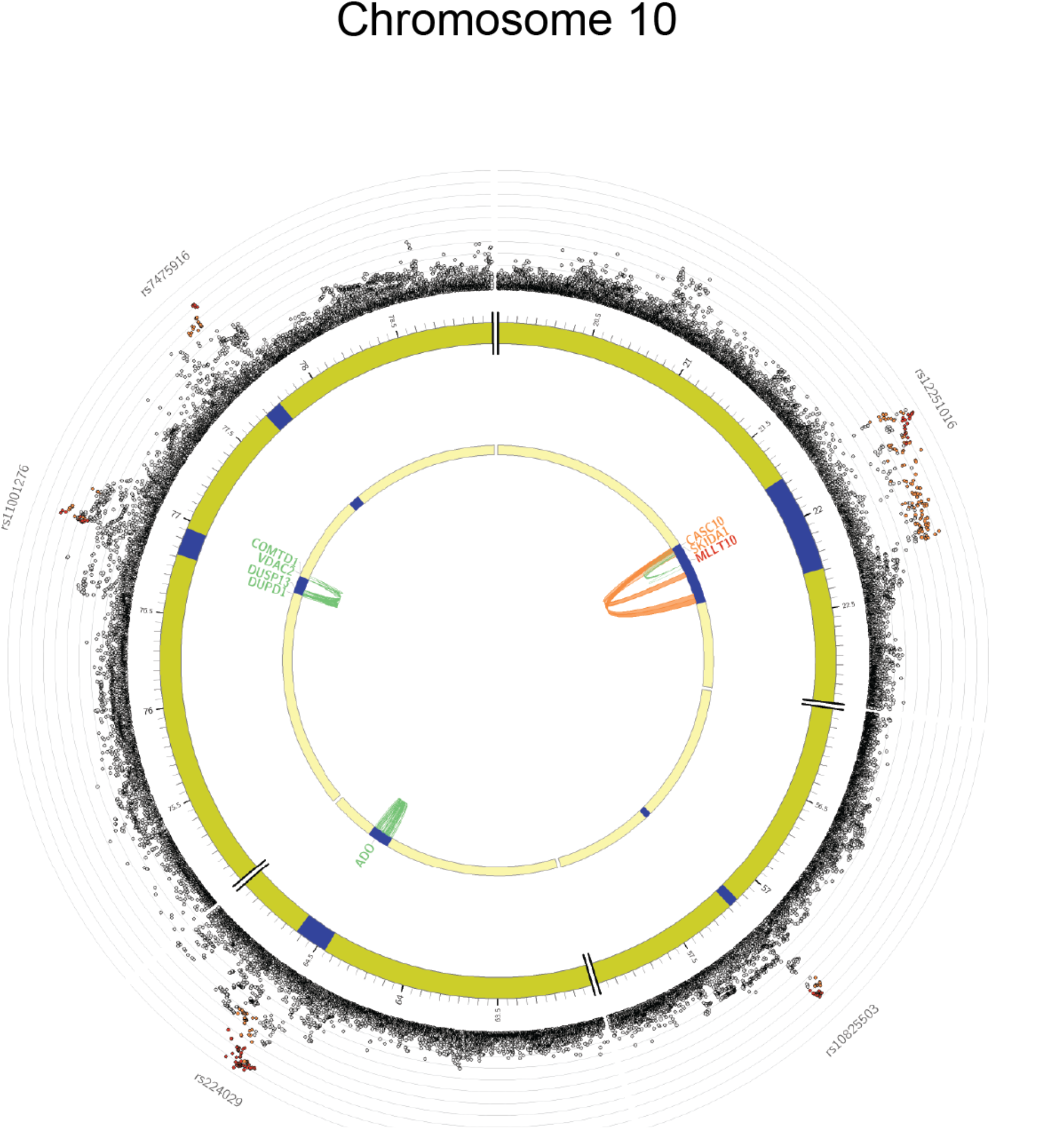
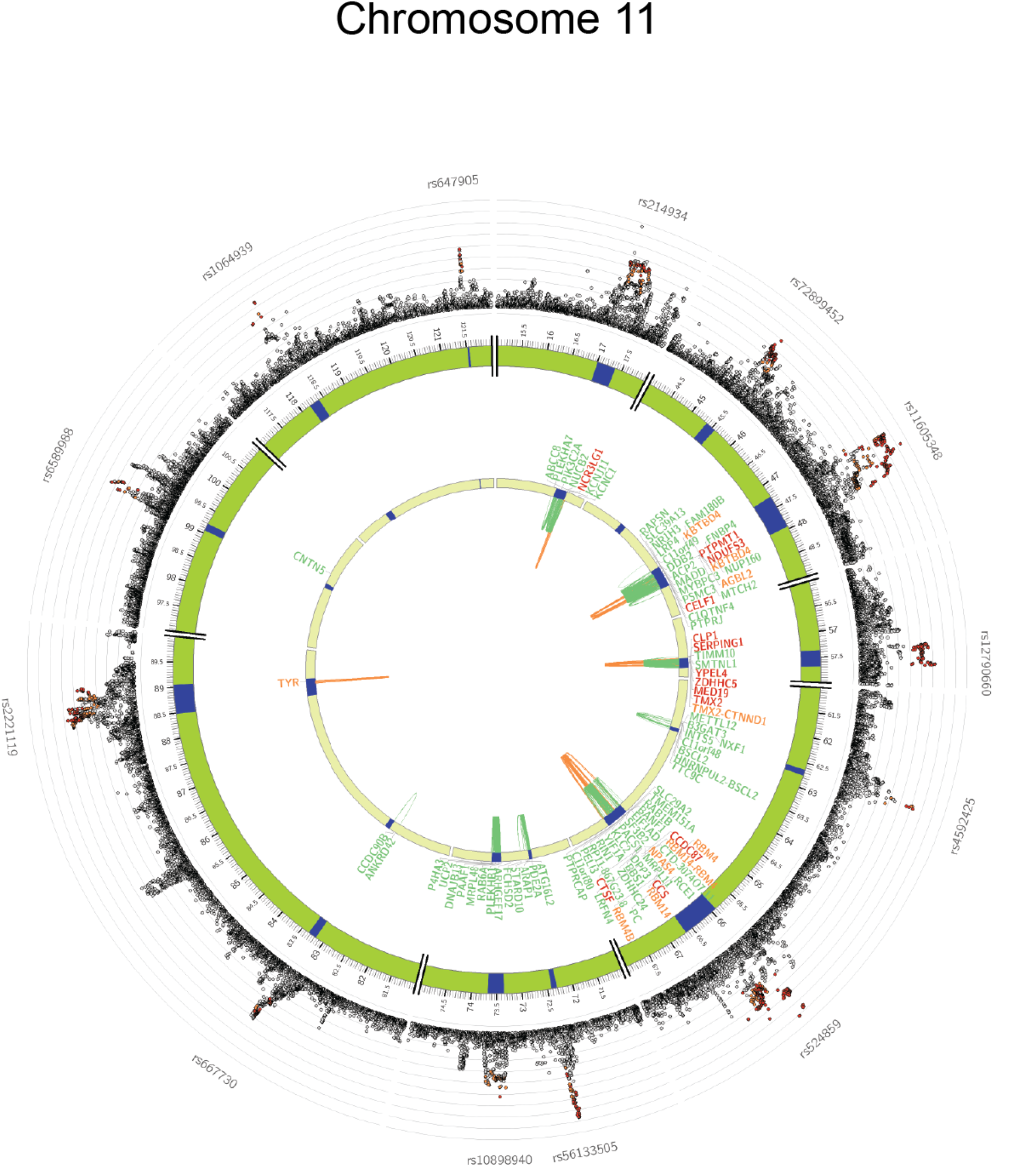
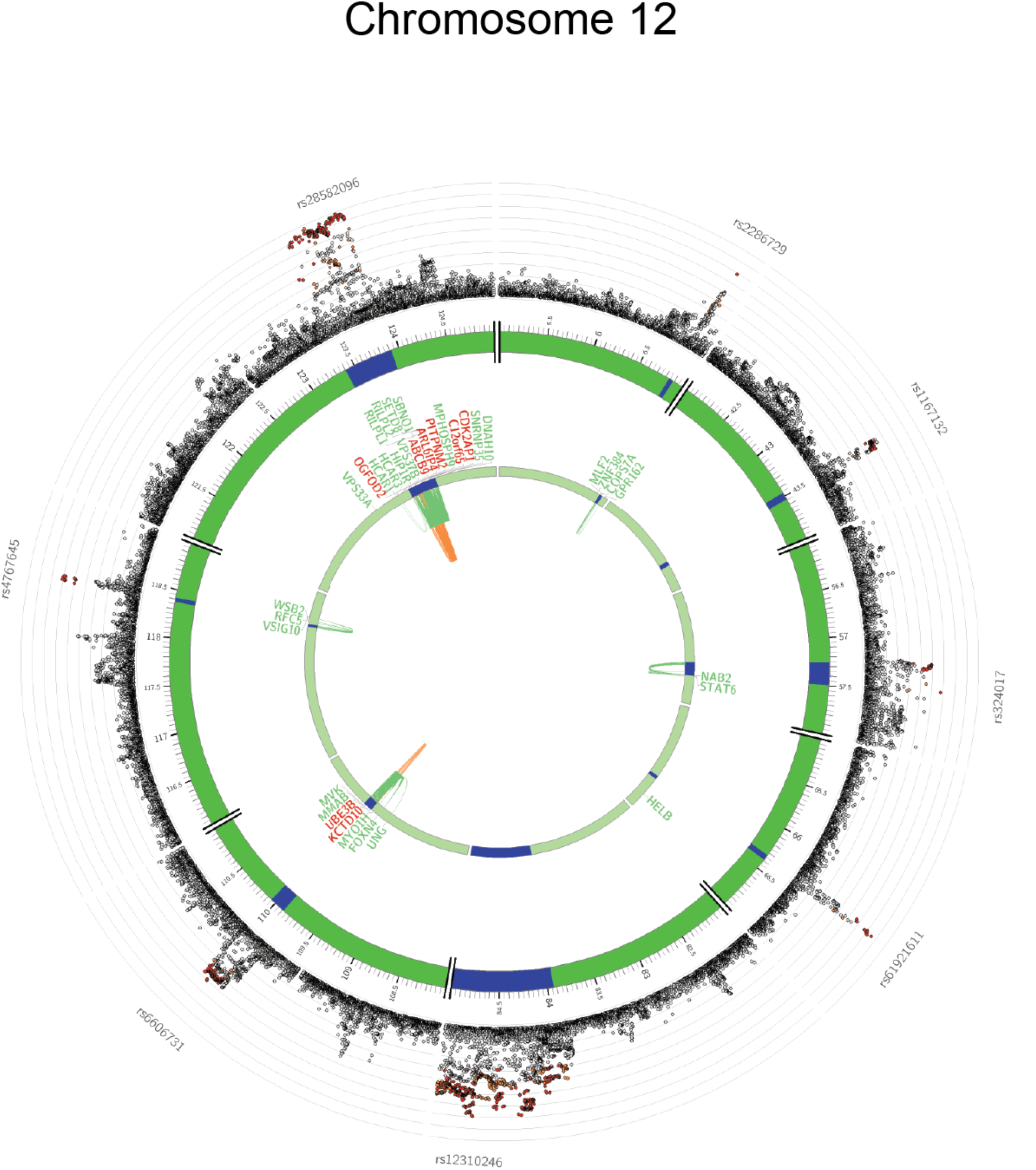
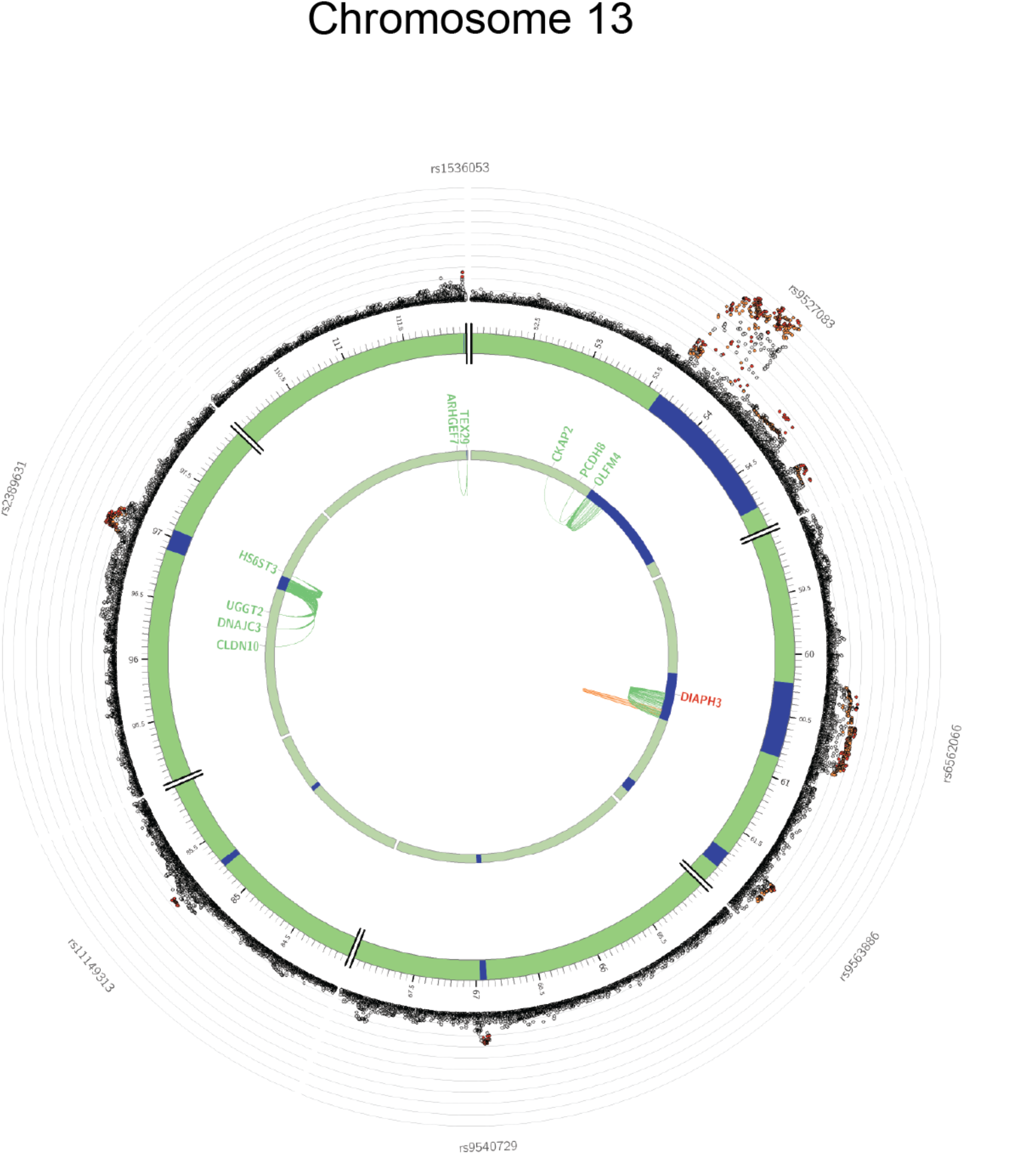
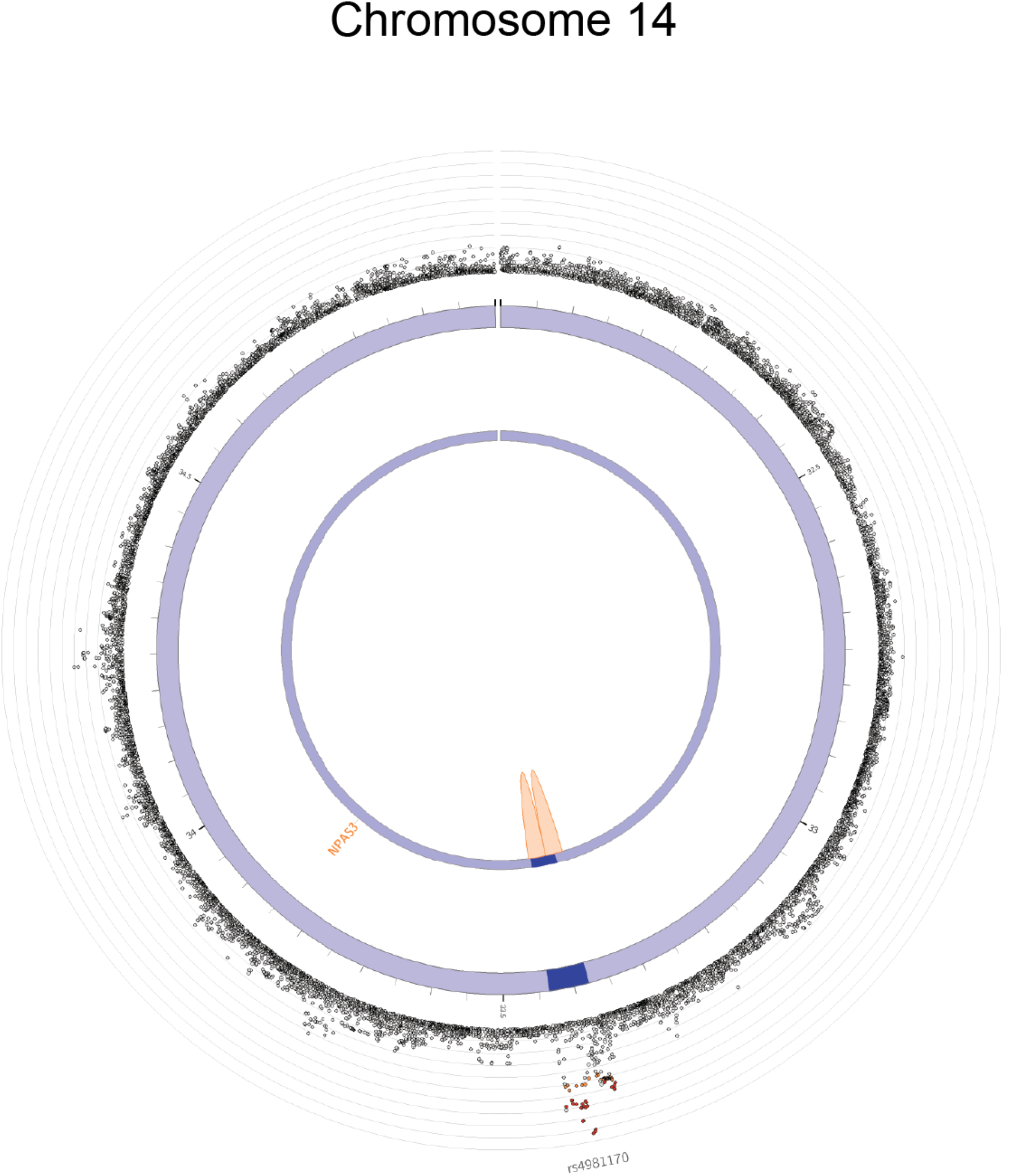
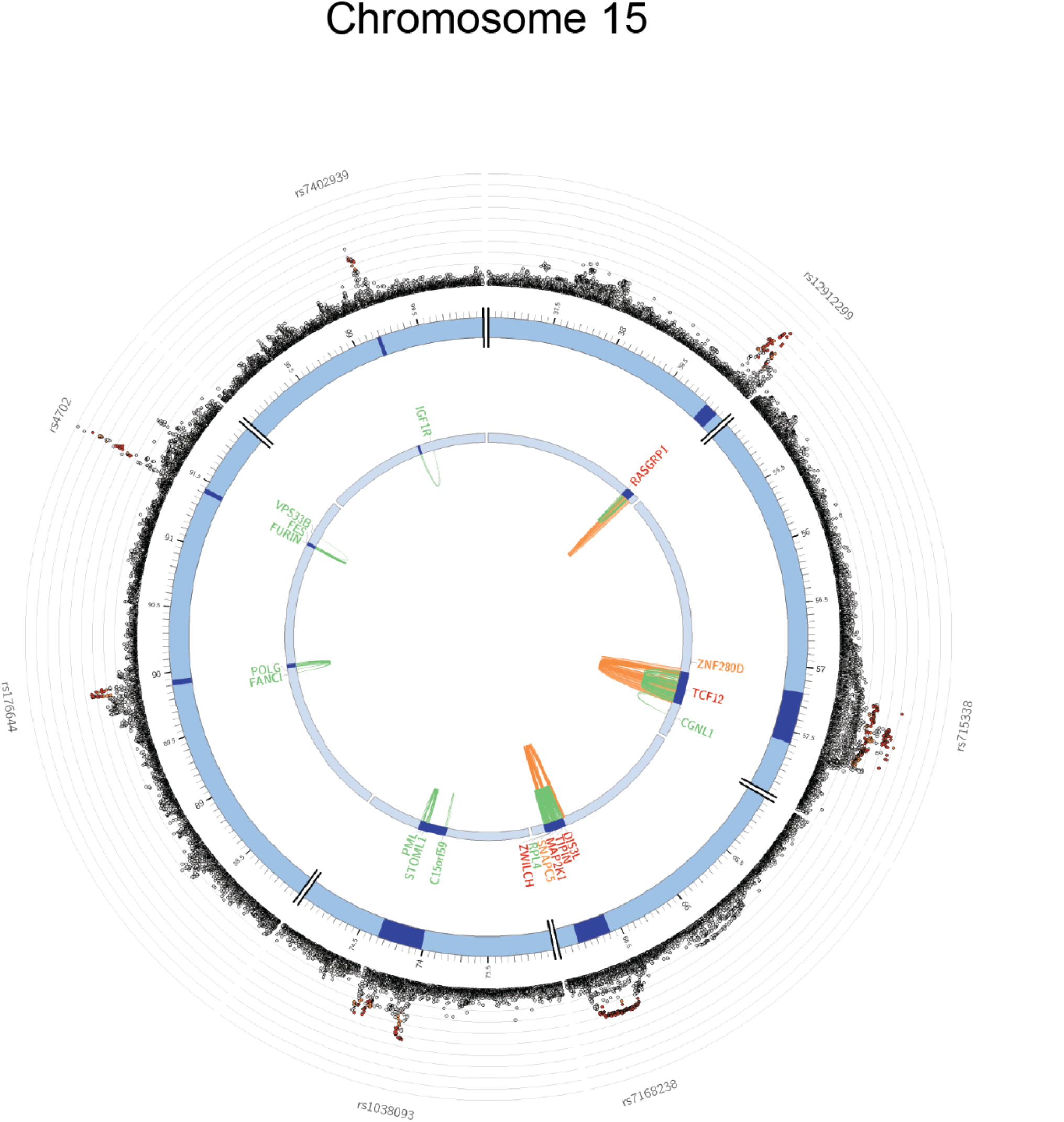
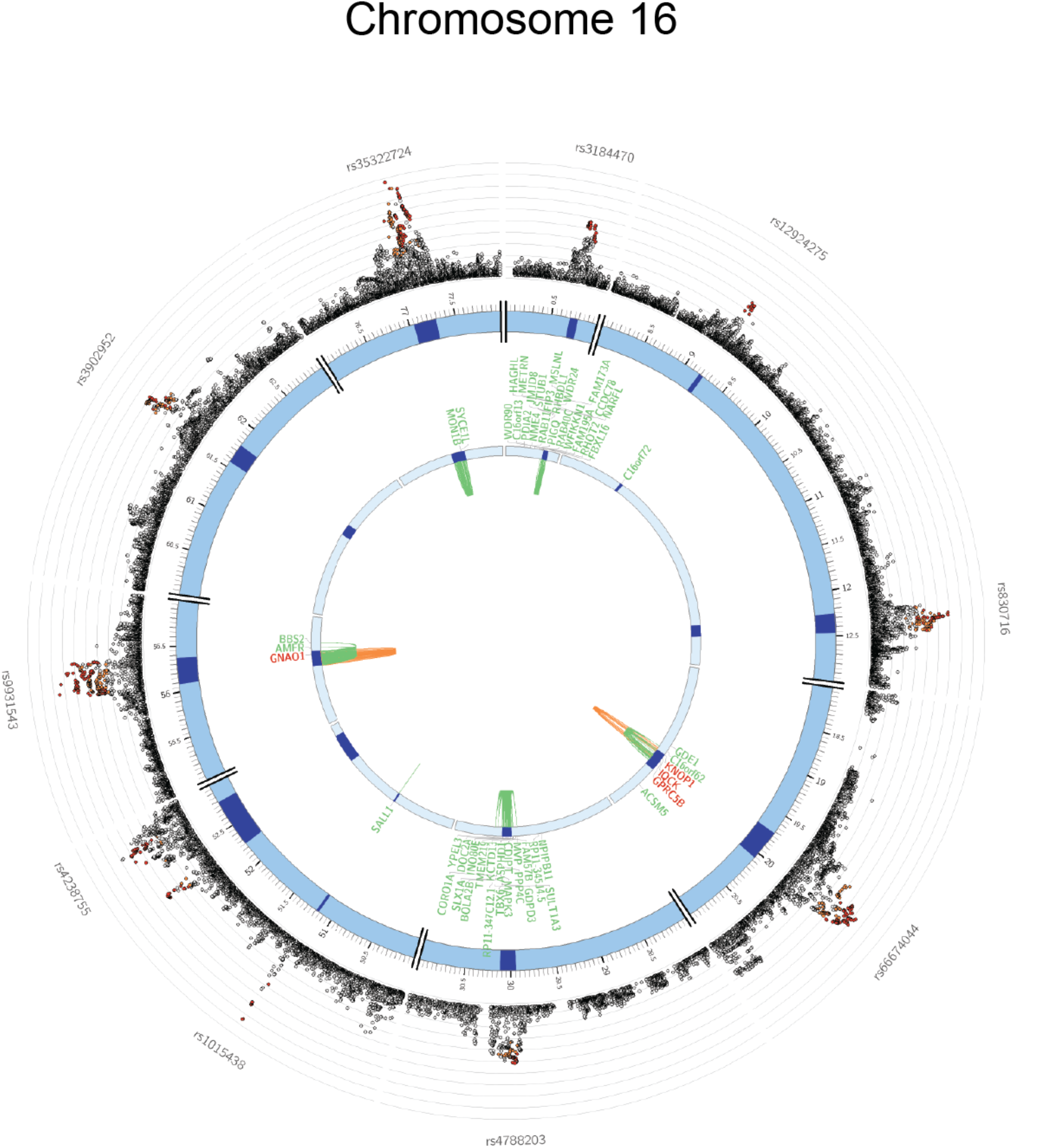
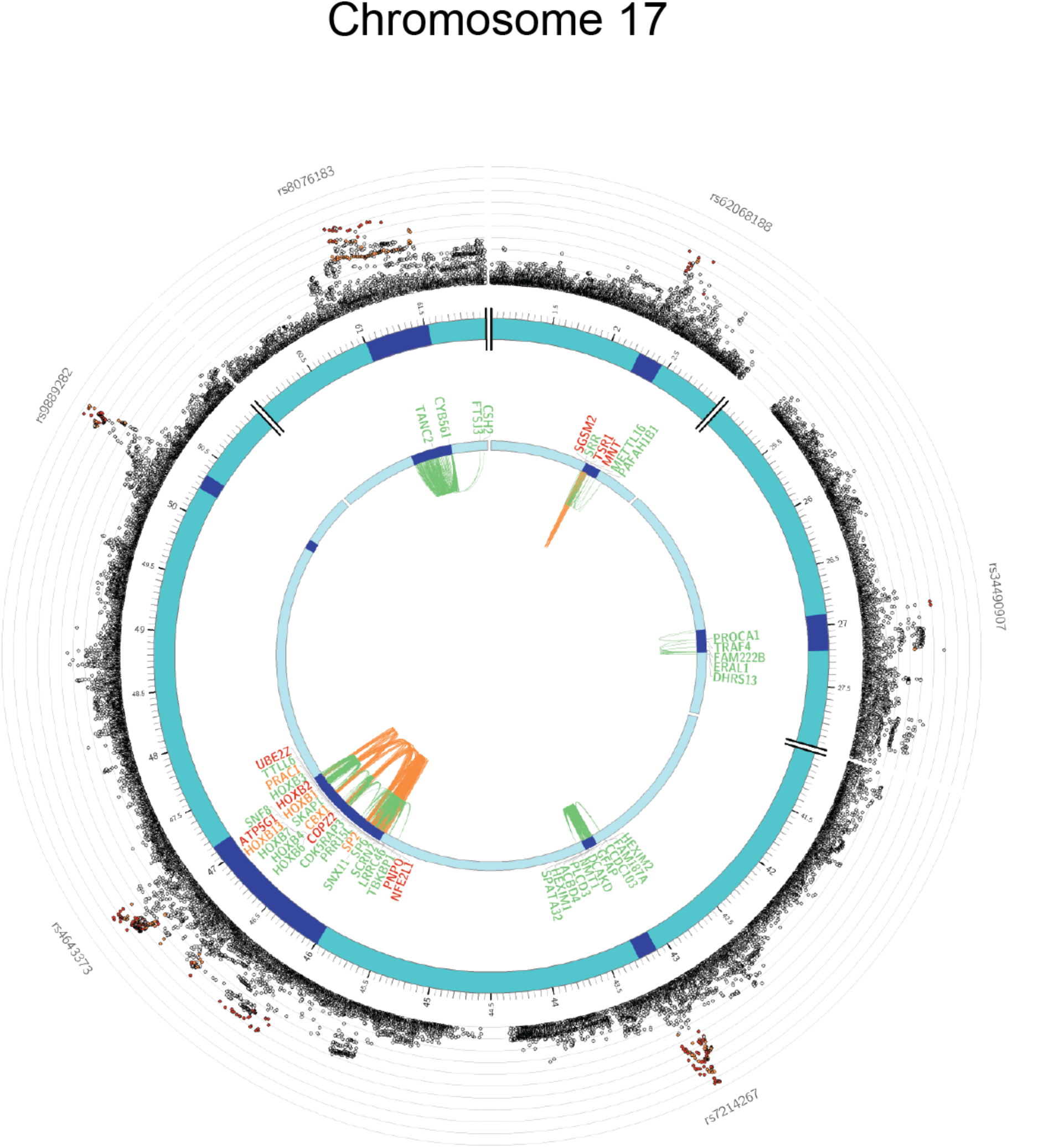
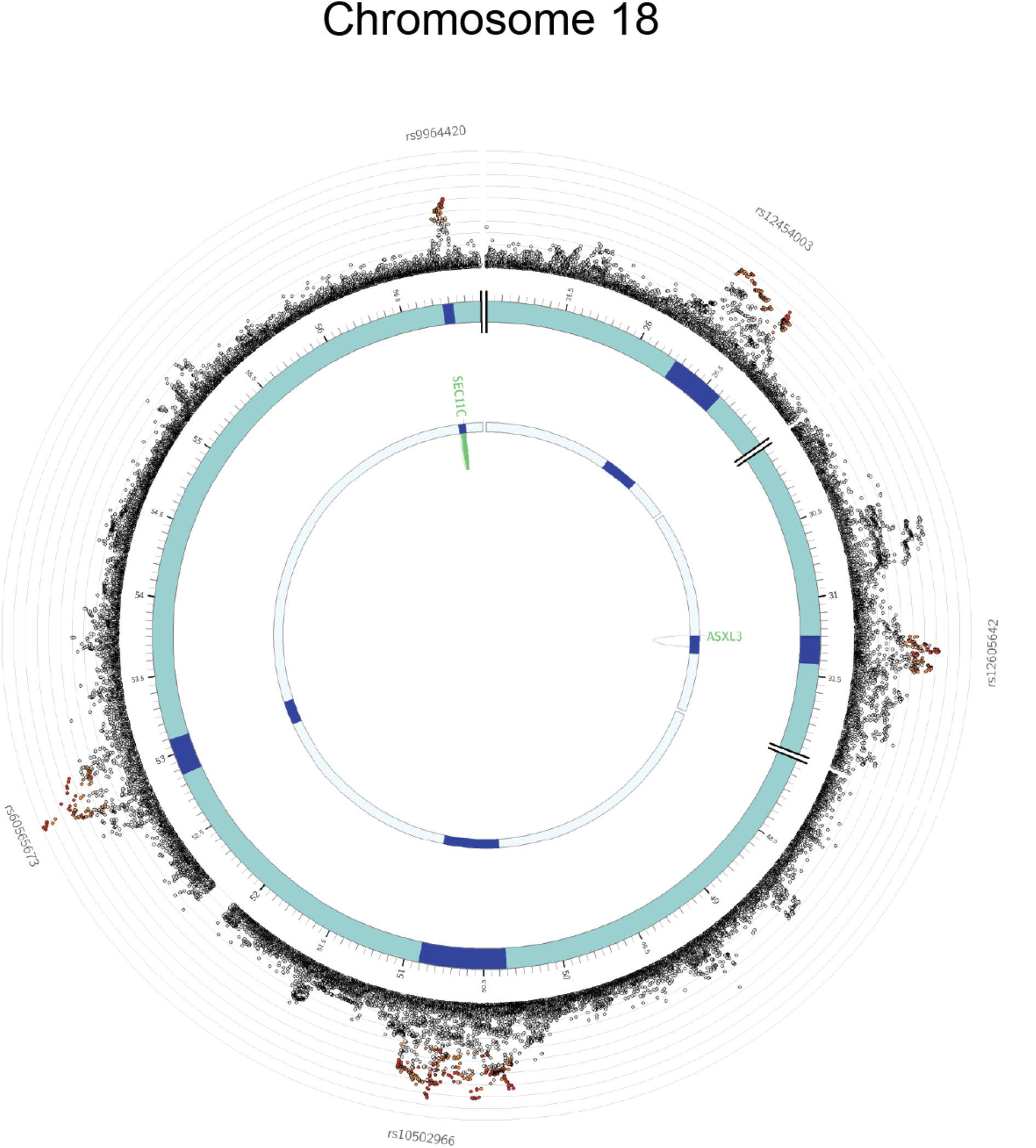
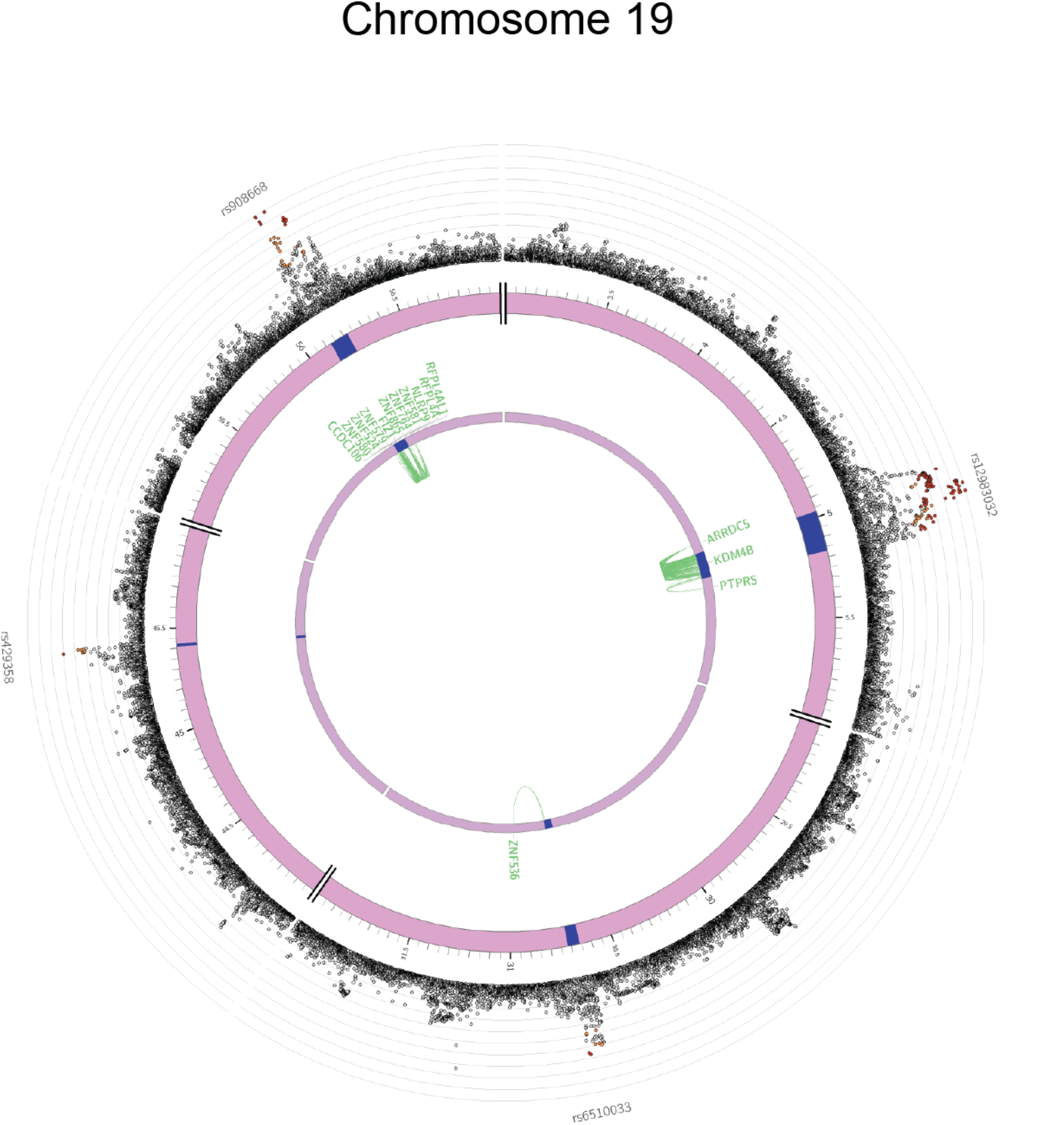
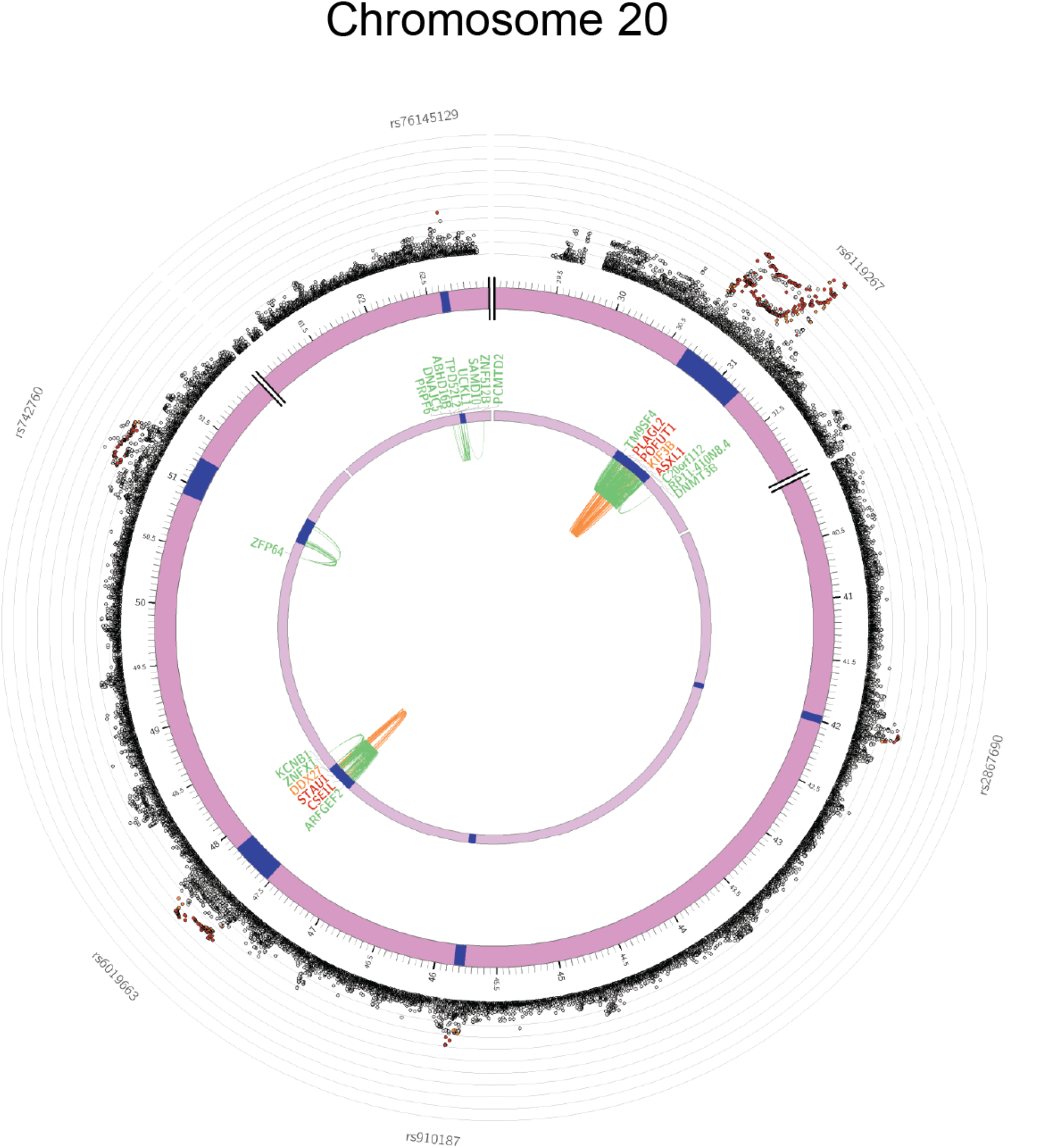
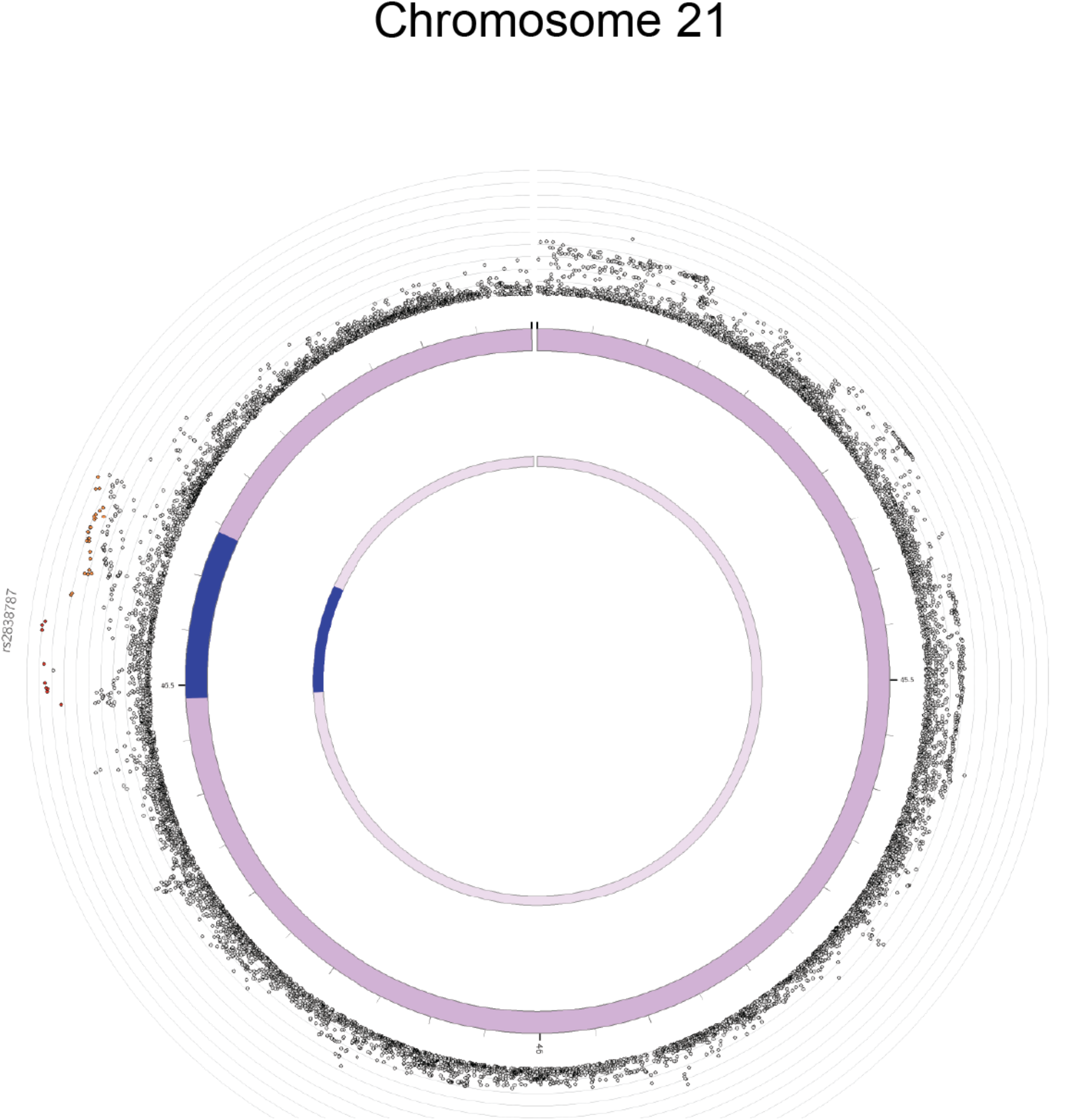
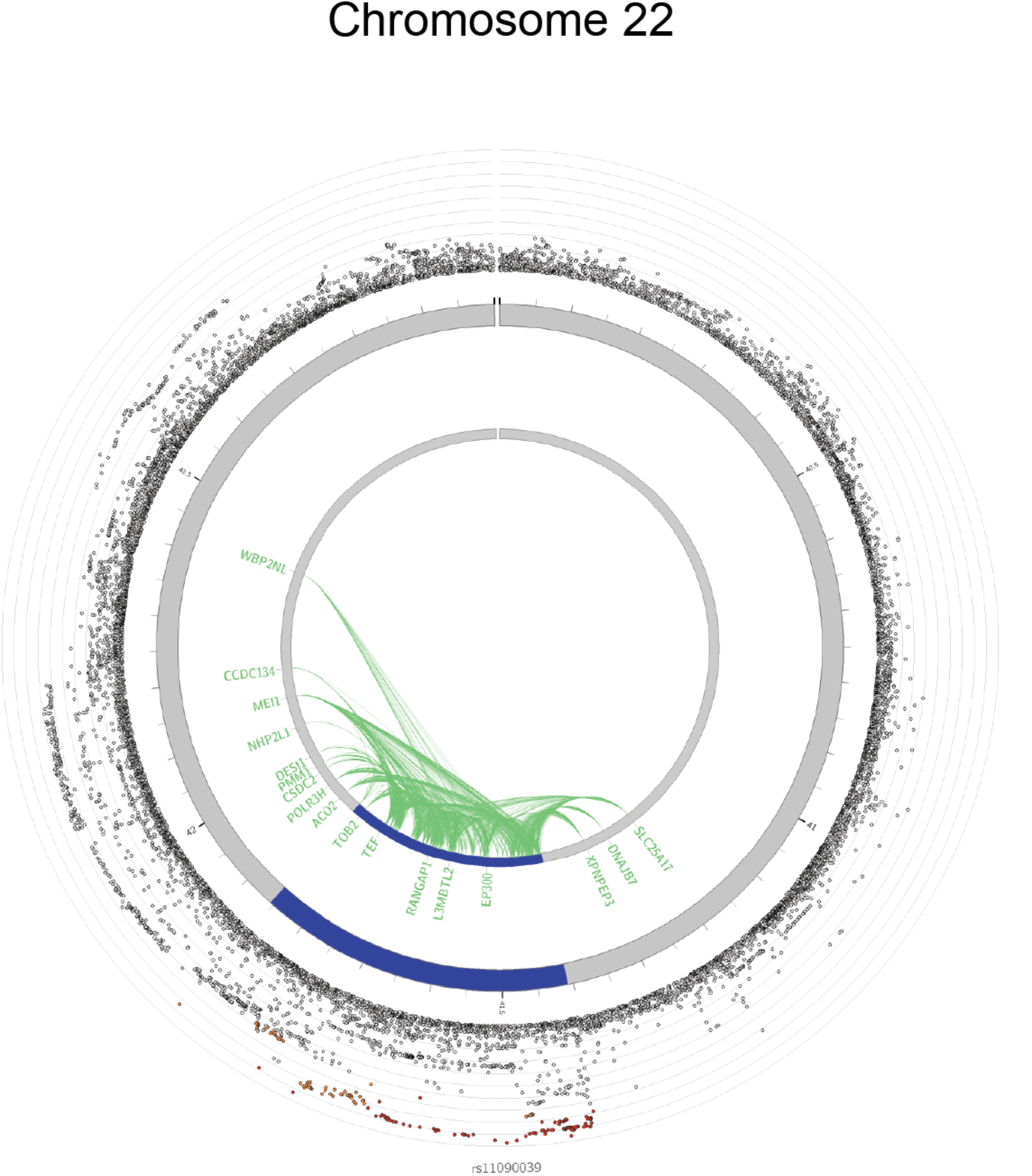

## Footnotes

1 We caution that only a limited set of brain tissues were included and thus we cannot rule out associations with many important areas such as pons, midbrain or thalamus based on this analysis.

2 We do note that for major depression the reverse MR could not be carried out due to an insufficient number of SNPs with a low P-value

